# ABHD17 enzymes regulate dynamic plasma membrane palmitoylation and N-Ras-dependent cancer growth

**DOI:** 10.1101/2020.05.21.108316

**Authors:** Jarrett R. Remsberg, Radu M. Suciu, Noemi A. Zambetti, Thomas W. Hanigan, Ari J. Firestone, Anagha Inguva, Amy Long, Nhi Ngo, Kenneth M. Lum, Cassandra L. Henry, Stewart K. Richardson, Marina Predovic, Ben Huang, Amy R. Howell, Micah J. Niphakis, Kevin Shannon, Benjamin F. Cravatt

**Author notes:** These authors contributed equally (Jarrett R. Remsberg, Radu M. Suciu, Noemi A. Zambetti).

## Abstract

A subset of Ras proteins, including N-Ras, depend on a palmitoylation/depalmitoylation cycle to regulate their subcellular trafficking and oncogenicity. General lipase inhibitors such as Palmostatin M block N-Ras depalmitoylation, but lack specificity and target several enzymes displaying depalmitoylase activity. Here, we describe ABD957, a potent and selective covalent inhibitor of the ABHD17 family of depalmitoylases, and show that this compound impairs N-Ras depalmitoylation in human acute myeloid leukemia (AML) cells. ABD957 produced partial effects on N-Ras palmitoylation compared to Palmostatin M, but was much more selective across the proteome, reflecting a plasma membrane-delineated action on dynamically palmitoylated proteins. Finally, ABD957 impaired N-Ras signaling and the growth of *NRAS*-mutant AML cells in a manner that synergizes with MEK inhibition. Our findings uncover a surprisingly restricted role for ABHD17 enzymes in modulating the N-Ras palmitoylation cycle and suggest that ABHD17 inhibitors may have value as targeted therapies for *NRAS*-mutant cancers.

---

*RAS* genes are the most common targets of dominant mutations in human cancer^1, 2^. There are three different *RAS* genes, which encode four highly homologous proteins (H-Ras, N-Ras, K-Ras4a, and K-Ras4b), and they are preferentially mutated in distinct tumor types^1, 2^. An elegant approach for directly targeting Ras oncoproteins involves designing covalent inhibitors targeting cysteine 12 of K-Ras^G12C 3-5^. However, it is more challenging to apply this approach to oncogenic amino acid substitutions that introduce side chains lacking reactive nucleophiles. Given the difficulties inherent in directly targeting oncogenic Ras, inhibitors of Ras effector molecules such as PI3K, Akt, mTOR, and MEK are being pursued as an alternative therapeutic strategy^6, 7^.

Ras proteins share very high homology throughout most of their protein sequence with the exception of the C-terminal “hypervariable region” (HVR), which contains signals that specify post-translational modifications required for proper subcellular localization^8^. The HVR of all four isoforms terminates with a CAAX motif, where the cysteine is prenylated by the farnesyltransferase FTase (or FNTA/FNTB). This lipid modification provides weak membrane binding affinity that is stabilized by a second signal motif. For K-Ras4b, this is provided by a polybasic lysine domain^9, 10^. By contrast, H-Ras, N-Ras, and K-Ras4a are *S*-palmitoylated at cysteine(s) adjacent to the CAAX motif ^8, 9^, and a dynamic cycle of palmitoylation/depalmitoylation mediated by palmitoyl acyl transferase (PAT) and serine hydrolase (SH) enzymes has been shown to regulate H-and N-Ras trafficking, subcellular localization, and function^11-13^.

Of the 23 mammalian PATs, only DHHC9 has been directly shown to catalyze N-Ras palmitoylation in conjunction with GOLGA7^14, 15^. The enzymes involved in depalmitoylating H- and N-Ras are presumed to be serine hydrolases based on studies showing that N-Ras depalmitoylation is blocked by the beta-lactone small molecules palmostatin B and M (Palm B and M)^12, 16^ or hexadecylfluorophosphonate (HDFP)^17^, as well as earlier work demonstrating that the yeast serine hydrolase APT1 (or, in humans, LYPLA1) can depalmitoylate H-Ras^18^. While LYPLA1 and LYPLA2 are inhibited by Palm B and M^19^, selective inhibitors or genetic knockdown of these serine hydrolases do not block the palmitoylation dynamics of N-Ras in cancer cells^20^, pointing to other enzymes being involved in this process. Using activity-based protein profiling (ABPP) methods, Lin and Conibear recently identified ABHD17 proteins, a poorly characterized group of three sequence-related serine hydrolases (ABHD17A, B, and C), as additional targets of Palm M and HDFP and showed that these enzymes can depalmitoylate N-Ras^20^. The ABHD17 enzymes are themselves palmitoylated on their N-termini, which promotes localization to the plasma membrane of cells^21^. Importantly, overexpression of ABHD17A was found to enhance N-Ras depalmitoylation and altered N-Ras subcellular localization in COS-7 cells, and, conversely, triple knockdown of ABHD17A/B/C by RNA interference impaired N-Ras palmitoylation turnover, albeit less effectively than Palm B, in HEK293T cells^20^.

Previous findings, taken together, support a potential role for ABHD17 enzymes in regulating N-Ras palmitoylation. However, ABHD17A, B, and C are all broadly expressed in most cell types, which makes characterization of their collective contribution to N-Ras palmitoylation challenging using genetic methods. It remains unclear, for instance, whether the partial blockade of N-Ras depalmitoylation observed in triple ABHD17A/B/C knockdown cells reflects an incomplete reduction in the expression of these enzymes or a contribution of other Palm B and M targets to the N-Ras palmitoylation cycle. Additionally, the studies performed to date relating ABHD17 enzymes to N-Ras depalmitoylation have occurred in heterologous cell types that do not depend on N-Ras as an oncogenic driver. Accordingly, it remains unknown whether ABHD17 enzymes regulate N-Ras palmitoylation in relevant biological systems and if disrupting these enzymes will affect the growth of *NRAS* mutant cancer cells. Finally, the broader potential substrate scope of ABHD17 enzymes beyond N-Ras and a handful of other palmitoylated proteins (e.g. PSD-95^20, 22^) remains unknown.

To address the aforementioned knowledge gaps, we describe herein a potent, selective, and cell-active covalent inhibitor of the ABHD17 enzymes and the characterization of its pharmacological effects in *NRAS* mutant cancer cell lines. We find that this compound – termed ABD957 – inhibits all three ABHD17 enzymes and partially impairs N-Ras depalmitoylation, ERK phosphorylation, and the growth of *NRAS* mutant cancer cells. ABD957 stabilizes N-Ras palmitoylation less completely than Palm M, but is also much more selective across the proteome, being restricted in its effects to plasma membrane-associated, dynamically palmitoylated proteins. ABD957 and Palm M also differentially affect the subcellular distribution of N-Ras, leading to the accumulation of this protein on the plasma membrane and intracellular membranes, respectively. Finally, we show that ABD957 synergizes with MEK inhibition to block *NRAS* mutant cancer cell growth. These data, taken together, indicate that ABHD17 enzymes perform a specialized function as plasma membrane-delineated *S*-depalmitoylases and that the chemical inhibition of these enzymes attenuates dynamic N-Ras depalmitoylation, resulting in impaired signaling and growth of *NRAS* mutant cancer cells.

## Results

### Discovery and characterization of ABHD17 inhibitors

The palmostatins (Palm B and Palm M)^12, 16^ and HDFP^17^ (**Fig. 1a**) are useful pharmacological tools for studying protein palmitoylation through blockade of depalmitoylase enzymes and, to date, remain the only reported inhibitors of ABHD17A/B/C^20^. HDFP is a broad-spectrum serine hydrolase inhibitor, targeting multiple depalmitoylases including LYPLA1/2, PPT1, ABHD10 and ABHD17A/B/C^17^. This compound^13, 17, 23^, along with other long-chain FPs^24, 25^, have been shown to preserve the palmitoylation state of several dynamically palmitoylated proteins, including Ras proteins^17, 20^, in mammalian cells. Palm B and M also inhibit several other serine hydrolases^19, 20^ and this lack of selectivity, along with presence of a metabolically labile beta-lactone, has limited the broad utility of these compounds in biological systems. Such factors motivated us to develop a more advanced chemical probe that selectively inhibits ABHD17 enzymes.

**Figure 1.**
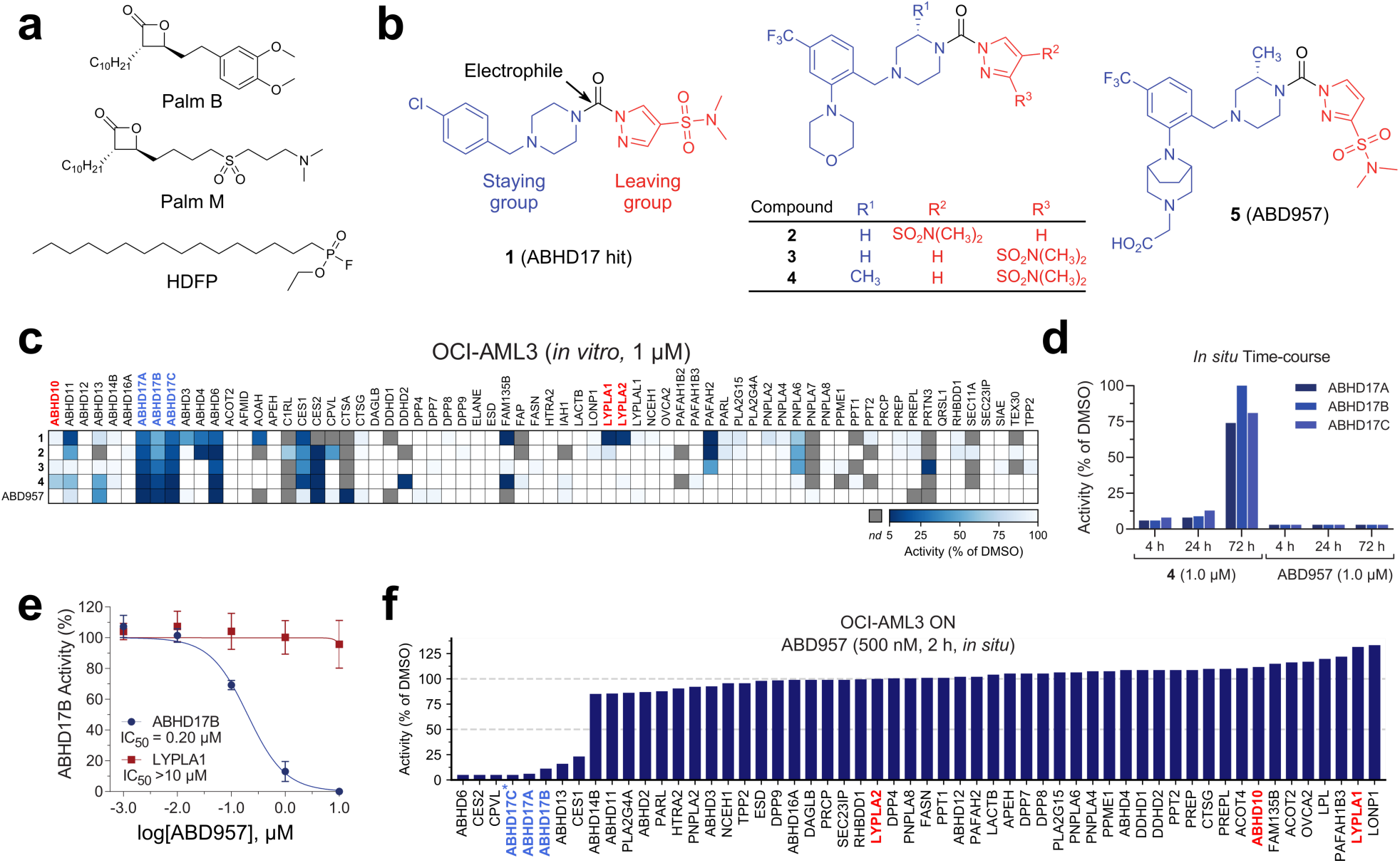
Discovery and characterization of ABD957 - a potent and selective inhibitor of the ABHD17 enzymes. **a**, Structures of broad-spectrum serine hydrolase inhibitors Palm B, Palm M and HDFP. **b**, Structures of pyrazole urea class of ABHD17 inhibitors discovered herein. **c**, MS-ABPP data of serine hydrolase activities in the particulate fraction of OCI-AML3 proteomes treated with compounds **1**-**5** (1 µM, 30 min). See **Supplementary Figure 1b** for MS-ABPP data of compounds tested at 10 µM and **Supplementary Table 1** for proteomic data. MS-ABPP data are from single experiments performed at the indicated concentrations for each compound. **d**, MS-ABPP data for an *in situ* time-course of ABHD17A/B/C inhibition by compounds **4** and ABD957 (1 µM) in THP1 cells, revealing maintenance of inhibition of ABHD17A/B/C over 72 h in cells treated with ABD957, but not with compound **4**. Data are from a single experiment representative of three independent experiments. **e**, IC_50_ curve for ABD957 inhibition of human ABHD17B (recombinantly expressed) and LYPLA1 (endogenous) activity in lysates of HEK293T cells (*in vitro*) measured by gel-ABPP. Data represent average values ± s.d. (n = 3 independent experiments). **f**, *In situ* MS-ABPP data for ABD957 (500 nM, 2 h) in GFP-N-Ras^G12D^-expressing OCI-AML3 (ON) cells confirming ABHD17A/B inhibition and selectivity across the majority of quantified serine hydrolases, including LYPLA1, LYPLA2, and ABHD10. Data represent three experiments corresponding to independent compound treatmentof cells, after which cell lysates were combined together for analysis by MS-ABPP. * One unique peptide used in quantification.

HDFP and palmostatins feature reactive electrophilic centers that inhibit serine hydrolases through covalent modification of the catalytic serine residue. While this feature likely contributes to a lack of specificity across the serine hydrolase class, more tempered electrophilic chemotypes, such as carbamates and ureas, have been shown to covalently inhibit individual serine hydrolases with excellent potency, selectivity, and cellular and *in vivo* activity^26-32^. The evaluation of candidate inhibitors of serine hydrolases has also benefited from the chemical proteomic technology activity-based protein profiling (ABPP), wherein activity-based probes showing broad reactivity with serine hydrolases are used to measure target engagement and selectivity of inhibitors in native biological systems^33-35^. We accordingly screened a serine hydrolase-directed compound library internally developed at Lundbeck La Jolla Research Center by gel-based ABPP in native mouse brain proteomes, where ABHD17 and other serine hydrolases can be visualized using a fluorescently (rhodamine) tagged fluorophosphonate (FP) activity-based probe. This screen furnished a piperazine-based pyrazole urea hit (**1, Fig. 1b**) that displayed moderate potency (IC_50_ [95% CI] = 0.93 [0.79-1.1] µM, **Table 1** and **Supplementary Fig. 1a, b**) against human ABHD17B, assayed by gel-based ABPP in stably tranduced HEK293T cells.

**Table 1.**
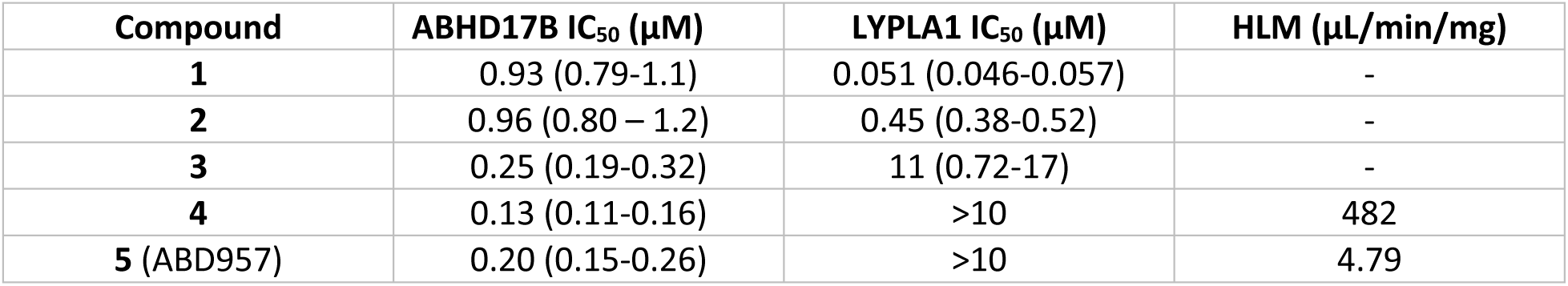
Properties of ABHD17 inhibitors. Potency values for inhibiting human ABHD17B and LYPLA1 were measured by gel-ABPP in lysates of HEK293T cells stably expressing recombinant human ABHD17B following a 30 min inhibitor preincubation. Inhibition of endogenously expressed LYPLA1 was also measured in these experiments in parallel. Data represent average values and 95% confidence intervals from three independent experiments. Intrinsic clearance was measured in human liver microsomes (HLM).

Compound **1** represented an attractive starting point as an ABHD17 inhibitor due to its simple, chemically tractable core and the presence of a moderately electrophilic pyrazole urea group resembling those found in other advanced chemical probes that covalently inhibit serine hydrolases^26, 30, 31, 36^. We next used a biotinylated FP probe^37^ and quantitative, mass spectrometry (MS)-based ABPP^31^ to assess the selectivity of **1** in the proteome of the human *NRAS* mutant acute myeloid leukemia (AML) cell line OCI-AML3^38^. Experiments performed with 1 and 10 µM of **1** confirmed simultaneous inhibition of all three A, B and C isoforms of ABHD17, as well as several additional serine hydrolases, including LYPLA1 and LYPLA2 (**Fig. 1c** and **Supplementary Fig. 1c**). We focused our initial optimization efforts on improving selectivity for ABHD17 enzymes, using cross-reactivity with human LYPLA1 as a guide, since this enzyme could be readily visualized by gel-based ABPP in ABHD17B-transduced HEK293T proteome (**Supplementary Fig. 1b**). We found that three modifications to **1** were sufficient to eliminate LYPLA1/2 activity while simultaneously improving inhibitory potency and selectivity for ABHD17s. First, introduction of a morpholino group to the benzyl moiety (compound **2, Fig. 1b**) resulted in a ∼10-fold loss of potency for LYPLA1 activity without impairing activity against ABHD17B (**Table 1**). Secondly, shifting the sulfonamide from the 4- to the 3-position on the pyrazole leaving group (compound **3**; **Fig. 1b**) diminished LYPLA1 activity further (IC_50_ ∼10 µM) and concurrently enhanced hABHD17B potency by ∼4-fold (IC_50_ [95% CI] = 0.25 [0.19-0.32] µM) (**Table 1**). MS-based ABPP experiments revealed that these modifications also had a favorable impact on global selectivity across the serine hydrolase family, removing off-targets such as ABHD11 (**Fig. 1c** and **Supplementary Fig. 1c**). Reasoning that increasing the steric demand around the electrophile could promote selectivity, as has been seen for other serine hydrolases^28, 30^, we introduced a methyl substituent on the piperazine staying group adjacent to the urea (compound **4, Fig. 1b**), which further boosted potency (IC_50_ [95% CI] = 0.13 [0.11-0.16] µM, **Table 1**) and shifted the selectivity profile - eliminating PAFAH2, but introducing ABHD10, a recently identified mitochondrial depalmitoylase^25^, as an off-target (**Fig. 1c** and **Supplementary Fig. 1c**). Furthermore, **4** suffered from poor metabolic stability in human liver microsomes (482 µL/min/mg, **Table 1**) and, when tested in cells, did not produce sustained inhibition of ABHD17s over 72 h (**Fig. 1d**), a common endpoint used for cell-based assays that measure, for instance, proliferation.

Additional modifications to **4**, specifically the introduction of a carboxymethyl-substituted diazobicyclo[3.2.1]octane in place of the morpholino group, furnished the more advanced chemical probe ABD957 (compound **5, Fig. 1b**) that displayed much lower microsomal clearance (4.8 µL/min/mg, **Table 1**) while maintaining good potency for ABHD17B (IC_50_ [95% CI] = 0.20 [0.15-0.26] µM, **Fig. 1e, Supplementary Fig. 1b**, and **Table 1**). MS-based ABPP revealed that ABD957 exhibits excellent selectivity across the serine hydrolase class, including avoidance of ABHD10 as an off-target (**Fig. 1c** and **Supplementary Fig. 1c**). ABD957 did cross-react with a handful of other off-targets across the >60+ serine hydrolases assayed by MS-based ABPP, but, to our knowledge, none of these enzymes have been reported to display protein depalmitoylation activity. Recognizing that cellular potency and selectivity were more pertinent parameters for assessing the suitability of ABD957 as a chemical probe, we performed MS-based ABPP experiments on OCI-AML3 “ON” (described below) cells treated *in situ* with ABD957 (500 nM) for 2 h (**Fig. 1f**). ABD957 blocked the activities of ABHD17A, B, and C under these conditions with good overall selectivity, including no cross-reactivity with LYPLA1, LYPLA2, or ABHD10. While ABD957 cross-reacted with a handful of other serine hydrolases, including CES1/2, ABHD6 and ABHD13, the global selectivity was markedly improved compared to previous ABHD17 inhibitors, such as Palm M and HDFP, which, consistent with previous studies^17, 20^, caused widespread serine hydrolase inhibition in cells (**Supplementary Fig. 1d**). Importantly, ABD957 (1 µM) produced sustained cellular inhibition of ABHD17A/B/C for 72 h (**Fig. 1d**), indicating that this compound was well-suited for a diverse array of cell-based experiments.

### ABHD17 inhibition attenuates N-Ras depalmitoylation in AML cells

To assess the impact of ABHD17 inhibition on N-Ras palmitoylation, we adapted a pulse-chase assay for measuring dynamic protein palmitoylation previously described by our lab^17^. We first generated sublines of OCI-AML3 cells in which we used RNA interference to reduce the expression of endogenous human N-Ras and then introduced into these cells either i) a murine GFP-N-Ras^G12D^ protein, or ii) a murine GFP-N-Ras^G12D^ variant where the palmitoylated HVR sequence was replaced with the non-palmitoylated HVR sequence of K-Ras4b (**Supplementary Fig. 2**). The two engineered cell lines expressing GFP-N-Ras^G12D^ and GFP-N-Ras^G12D, KRAS HVR^ proteins were termed “ON” and “ONK”, respectively. We next evaluated the effect of ABD957 and other compounds on N-Ras palmitoylation dynamics by pre-incubating ON or ONK cells with compounds or DMSO control for 1 h, then treating cells with the clickable palmitate analog 17-octadecynoic acid (17-ODYA^21, 39, 40^; 20 µM) for 1 h, followed by a 1 h chase period where the media was changed to remove 17-ODYA and supplemented with fresh inhibitor (**Fig. 2a**, left and middle panels). The palmitoylation state of N-Ras was then determined by anti-GFP immunoprecipitation, on-bead copper-catalyzed azide-alkyne cycloaddition (CuAAC)^41, 42^ to a rhodamine-azide (Rh-N_3_) reporter tag, and visualization of N-Ras palmitoylation signals by SDS-PAGE and in-gel fluorescence scanning (**Fig. 2a**, right lower panel). In this assay format, a compound that inhibited N-Ras depalmitoylation could be identified by comparing 17-ODYA labeling signals post-chase to a DMSO control sample. We also used a similar experimental protocol to survey the proteome-wide effects of candidate depalmitoylation inhibitors by performing CuAAC with Rh-N_3_ on lysates from 17-ODYA-treated cells (**Fig. 2a**, right upper panel).

**Figure 2.**
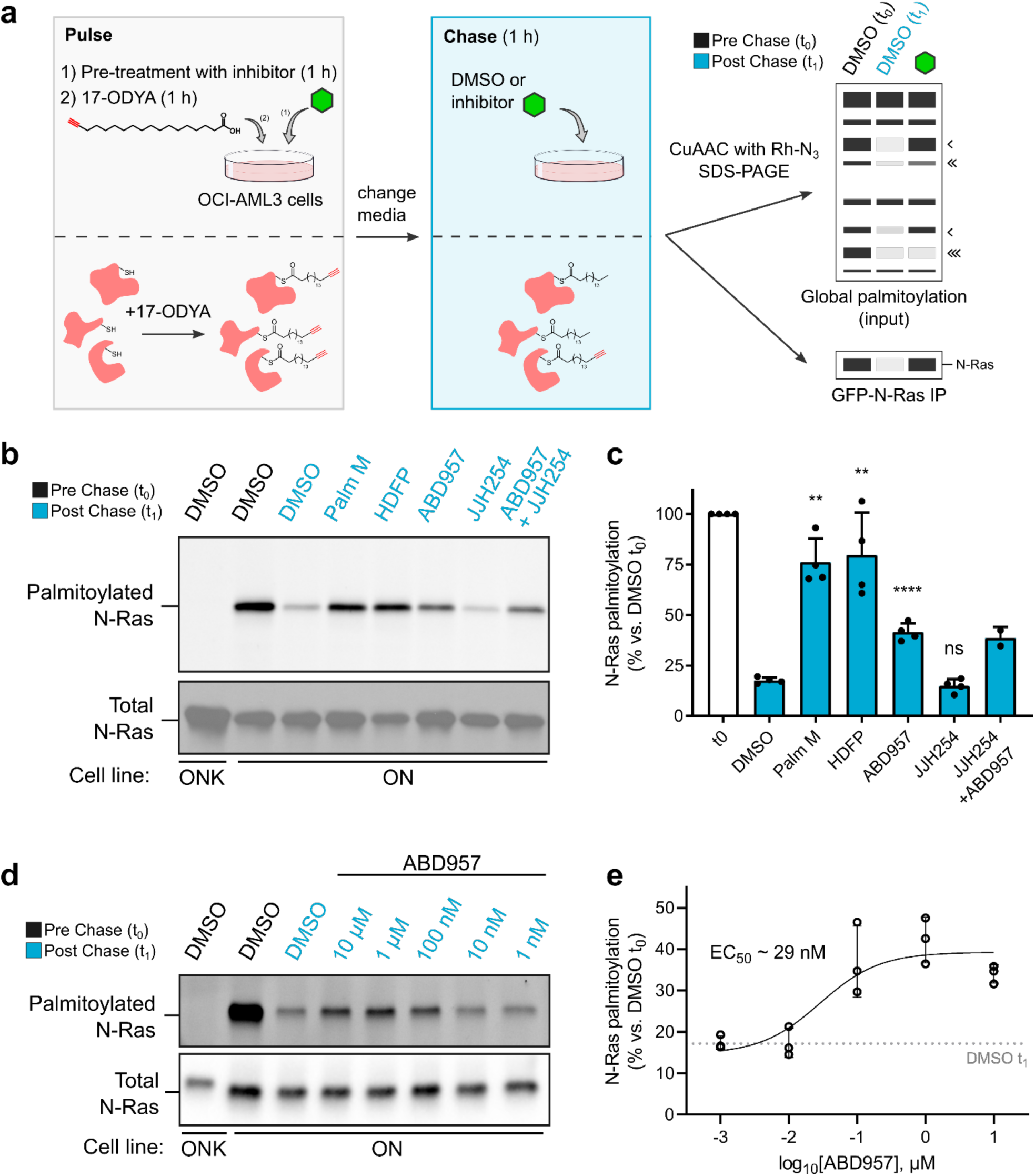
Effects of inhibitor treatment on the dynamic palmitoylation state of N-Ras. **a**, Schematic of a dynamic palmitoylation assay using metabolic labeling with 17-ODYA. Black arrowheads next to mock gel mark proteins that show dynamic palmitoylation fully (one arrowhead), partially (two arrowheads), or not (three arrowheads) preserved by serine hydrolase inhibition. **b**, Representative gel measuring N-Ras palmitoylation by 17-ODYA comparing effect of treatment with Palm M (10 µM), HDFP (20 µM), ABD957 (500 nM), and JJH254 (1 µM) in OCI-AML3 cells stably overexpressing GFP-N-Ras^G12D^ (ON) with GFP-N-Ras^G12D-KRAS-HVR^ (ONK) as a control (upper panel). N-Ras was immunoprecipitated via GFP and the degree of palmitoylation visualized by rhodamine attached via CuAAC to the alkyne of 17-ODYA. Total N-Ras content was measured by western blotting of GFP enrichments (lower panel). **c**, Quantification of inhibitor effects on dynamic palmitoylation. Data present average values ± s.d. (n = 4 independent experiments for all samples except ABD957/JJH254 for which n = 2). Statistical significance was calculated with unpaired two-tailed Student’s *t*-tests. **d**, Representative gel measuring N-Ras palmitoylation by 17-ODYA in the presence of varying concentrations of ABD957 (upper panel). Dotted line represents mean of residual post-chase DMSO (t_1_) signal. N-Ras was enriched and visualized as described in **b**. Total N-Ras content was measured by western blotting of GFP enrichments (lower panel). **e**, Quantification of concentration-dependent effects of ABD957 on N-Ras palmitoylation. Data represent average values ±s.d. (n = 3 independent experiments).

Consistent with previous studies^12, 16, 17, 20^, we found that the promiscuous lipase inhibitors Palm M and HDFP near-completely preserved N-Ras palmitoylation following the chase period (**Fig. 2b, c**). ABD957 also protected N-Ras from depalmitoylation, but to a lesser degree than Palm M or HDFP (**Fig. 2b, c**). In contrast, the dual LYPLA1 and LYPLA2 inhibitor JJH254^43^ did not affect N-Ras palmitoylation dynamics, either when administered on its own or in combination with ABD957 (**Fig 2b, c**). As expected, palmitoylation signals were not observed in ONK cells (**Fig. 2b, d**), indicating that the GFP-N-Ras^G12D, KRAS HVR^ protein does not undergo palmitoylation in the pulse-chase experiment.

The partial stabilization of N-Ras palmitoylation observed with ABD957 was concentration-dependent and showed an estimated IC_50_ value of 29 nM (**Fig. 2d, e**), which is consistent with the inhibition of ABHD17A/B/C in cells treated with 500 nM of this compound, as measured by MS-ABPP (**Fig. 1f**). Proteome-wide surveys of palmitoylation dynamics by SDS-PAGE revealed further that, while both Palm M and HDFP broadly affected the palmitoylation state of proteins in ON cells, ABD957 had a much more limited impact on dynamically palmitoylated proteins (**Supplementary Fig. 3a**). Intrigued by the distinct profiles of different depalmitoylation inhibitors, we next evaluated the effect of these compounds on global protein palmitoylation using quantitative MS-based proteomics.

### ABHD17 inhibition regulates the palmitoylation of plasma membrane proteins

We evaluated the impact of compounds on global palmitoylation dynamics using a similar pulse-chase protocol to that previously described^17^ and shown in **Fig. 2a**, where 17-ODYA-labeled proteins were conjugated by CuAAC to biotin-N_3_ to enable streptavidin enrichment of these labeled proteins followed by quantitative, multiplexed MS-based proteomics using tandem mass tags (TMT)^44^. We also required that the enrichment of 17-ODYA-labeled proteins show sensitivity to hydroxylamine treatment for designation as palmitoylated proteins, as reactivity with hydroxylamine is a characteristic feature of chemically labile thioester adducts formed between fatty acids and cysteine residues on proteins^45^. In total, 224 palmitoylated proteins were identified in ON cells (**Supplementary Table 1**). Consistent with past studies^17, 23^ only a modest subset (nineteen) of these proteins, which included N-Ras, displayed clear evidence of dynamic palmitoylation in pulse-chase experiments of ON cells, as reflected by at least two-fold reductions in 17-ODYA modification following the 1 h chase period (t_0_/t_1_ ≥ 2) (**Fig. 3a, b**, green and red proteins rightward of vertical dashed red line; and **Supplementary Table 1**). The palmitoylation states of most of these proteins were stabilized by treatment with Palm M (10 µM) (**Fig. 3a**, red proteins; and **Supplementary Table 1**). In contrast, ABD957 (500 nM) preserved the palmitoylation state of only a select few proteins, including N-Ras (**Fig. 3b**, red proteins). Another noteworthy difference between Palm M and ABD957-treated cells is that Palm M increased the apparent palmitoylation state of an additional set of proteins that otherwise did not show evidence of dynamic palmitoylation (**Fig. 3a**, blue proteins and **Supplementary Fig. 4a**). We are unsure of the mechanistic basis for these changes, but they point to a much broader effect of Palm M on the palmitoylated proteome compared to ABD957.

**Figure 3.**
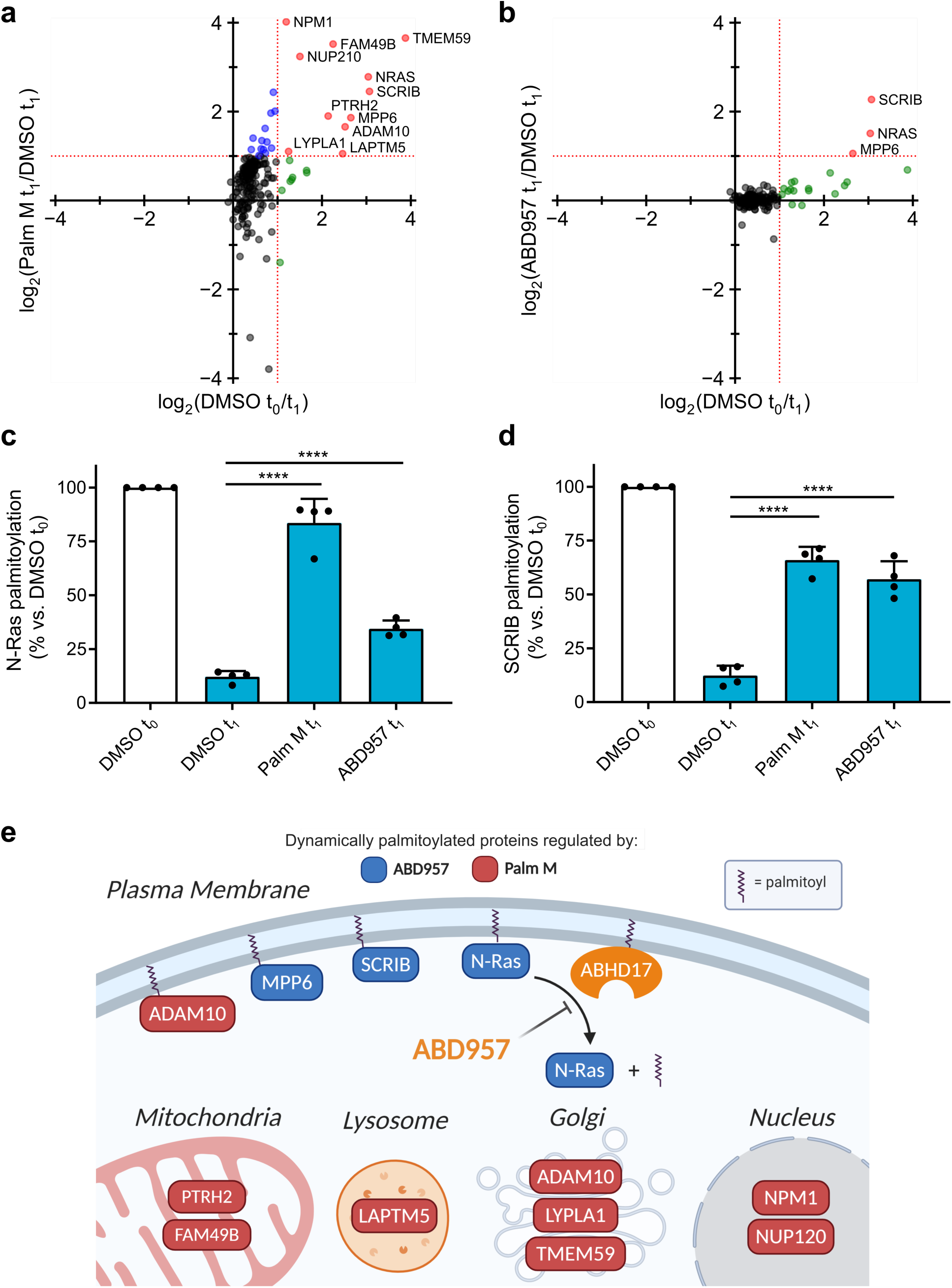
Global palmitoylation effects of Palm M and ABD957 in leukemia cells. **a, b**, Mass spectrometry (MS)-based profiling of ON cells as described in **Figure 2a**. Cells were preincubated with Palm M (10 µM) or ABD957 (500 nM) for 1 h, metabolically labeled with 20 µM 17-ODYA for 1 h (t_0_) and chased with media lacking 17-ODYA for 1 h (t_1_) containing Palm M (10 µM, **a**), ABD957 (500 nM, **b**), or DMSO control. Scatter plots compare log_2_ ratios of palmitoylated proteins in experiments measuring dynamic palmitoyation (DMSO t_o/_ t_1_; x-axis) verus effect of inhibitor treatment (inhibitor t_1_/DMSO t_1_; y-axis). Proteins shown were designated as palmitoylated based on showing hydroxylamine sensitivity (≥50% reduction in enrichment following hydroxylamine treatment) in other experiments (**Supplementary Table 2**). Proteins in red are both dynamically palmitoylated (DMSO t_0_/DMSO t_1_ ≥ 2 fold) and preserved in their palmitoylation state by inhibitor treatment (inhibitor t_1_/DMSO t_1_ ≥ 2 fold), green are dynamically palmitoylated, but not preserved by inhibitor treatment (inhibitor t_1_/DMSO t_1_ < 2 fold), and blue are proteins that did not display evidence of dynamic palmitoylation (DMSO t_0_/DMSO t_1_ < 2 fold), but showed higher palmitoylation signals following inhibitor treatment (inhibitor t_1_/DMSO t_1_ ≥ 2 fold). Red dotted lines represent 2-fold ratio values. Data represent average values (n = 4 from two biological replicates). **c, d**, Bar graphs quantifying N-Ras and SCRIB palmitoylation from MS-based proteomic experiments. Data represent average values relative to DMSO t_0_ ± s.d. (n = 4 from two biological replicates). Statistical significance was calculated with unpaired two-tailed Student’s *t*-tests. **e**, Cartoon diagram depicting the subcellular localization of proteins showing dynamic palmitoylation that was inhibited by Palm M (blue) or both Palm M and ABD957 (red).

In agreement with our focused palmitoylation assays on N-Ras (**Fig. 2b**), the palmitoylation state of this protein was more completely preserved by Palm M compared to ABD957 (**Fig. 3c**). Notably, the quantification of N-Ras signals could be performed in ON cells using both peptides that are unique to N-Ras (relative to other Ras isoforms) and peptides from the fused GFP protein as a surrogate, and both sets of peptides provided similar data supporting the preservation of N-Ras palmitoylation by Palm M and ABD957 (**Supplementary Fig. 4b** and **Supplementary Table 2**). In addition to N-Ras, ABD957 substantially preserved the palmitoylation state of SCRIB (**Fig. 3d**), a tumor suppressor protein that depends on palmitoylation for proper plasma membrane localization and function^46^, and MPP6, a palmitoylated guanylate kinase that has been shown to localize to the basolateral plasma membrane of intestinal epithelial cells^47^. In the case of SCRIB, ABD957 produced a degree of protection of palmitoylation that matched that observed for Palm M (**Fig. 3d**).

When considering possible reasons for the more restricted effect of ABD957 on the palmitoylated proteome compared to Palm M, we noted that the three main proteins affected by ABD957 all reside at the plasma membrane (**Fig. 3e**). In contrast, dynamically palmitoylated proteins that were affected by Palm M, but not ABD957, localized to other subcellular compartments (e.g., Golgi (LYPLA1, TMEM59)^48, 49^, nucleus (NPM1, NUP210)^50, 51^, lysosome (LAPTM5)^52^, mitochondria (FAM49B, PTRH2)^53, 54^) (**Fig. 3e**), where they may not interface with plasma membrane-localized ABHD17 enzymes. One possible exception is the metalloproteinase ADAM10, which localizes to both the Golgi apparatus and plasma membrane^55^, and, for which the palmitoylation state was preserved by Palm M, but not ABD957. Our quantitative proteomic data thus indicate that the ABHD17 enzymes regulate the dynamic palmitoylation state of N-Ras and a discrete set of plasma membrane-localized proteins in leukemia cells.

### ABD957 and Palm M differentially affect N-Ras localization in leukemia cells

Previous studies have shown that Palm B and M alter the subcellular localization of N-Ras, leading to redistribution of this protein from the plasma membrane to endomembranes^12, 16, 56^. This effect was also shown to depend on the presence of the N-Ras HVR in transduced primary hematopoietic cells^56^. Using live-cell confocal microscopy, we monitored GFP-N-Ras localization in ON cells following treatment with Palm M or ABD957. Consistent with previous studies^12, 16^, we found that Palm M promoted the time-dependent accumulation of N-Ras at endomembranes (**Fig. 4a**, upper panels). The intracellular N-Ras showed partially overlapping distribution with the Golgi marker N-acetylgalactosaminyltransferase (GALNT2, visualized as an RFP-tagged protein) (**Fig. 4b** and **Supplementary Fig. 5**). ABD957, on the other hand, promoted the retention of GFP-N-Ras at the plasma membrane without evidence of endomembrane redistribution (**Fig 4a-c** and **Supplementary Fig. 5**). The effect of ABD957 was selective for palmitoylated N-Ras, as it was observed in ON, but not ONK cells (**Fig. 4c**). These cell imaging results indicate that ABHD17 inhibition, in addition to partially protecting the palmitoylation state of N-Ras, leads to the accumulation of this protein at the plasma membrane, where ABHD17 enzymes are themselves localized^21^.

**Figure 4.**
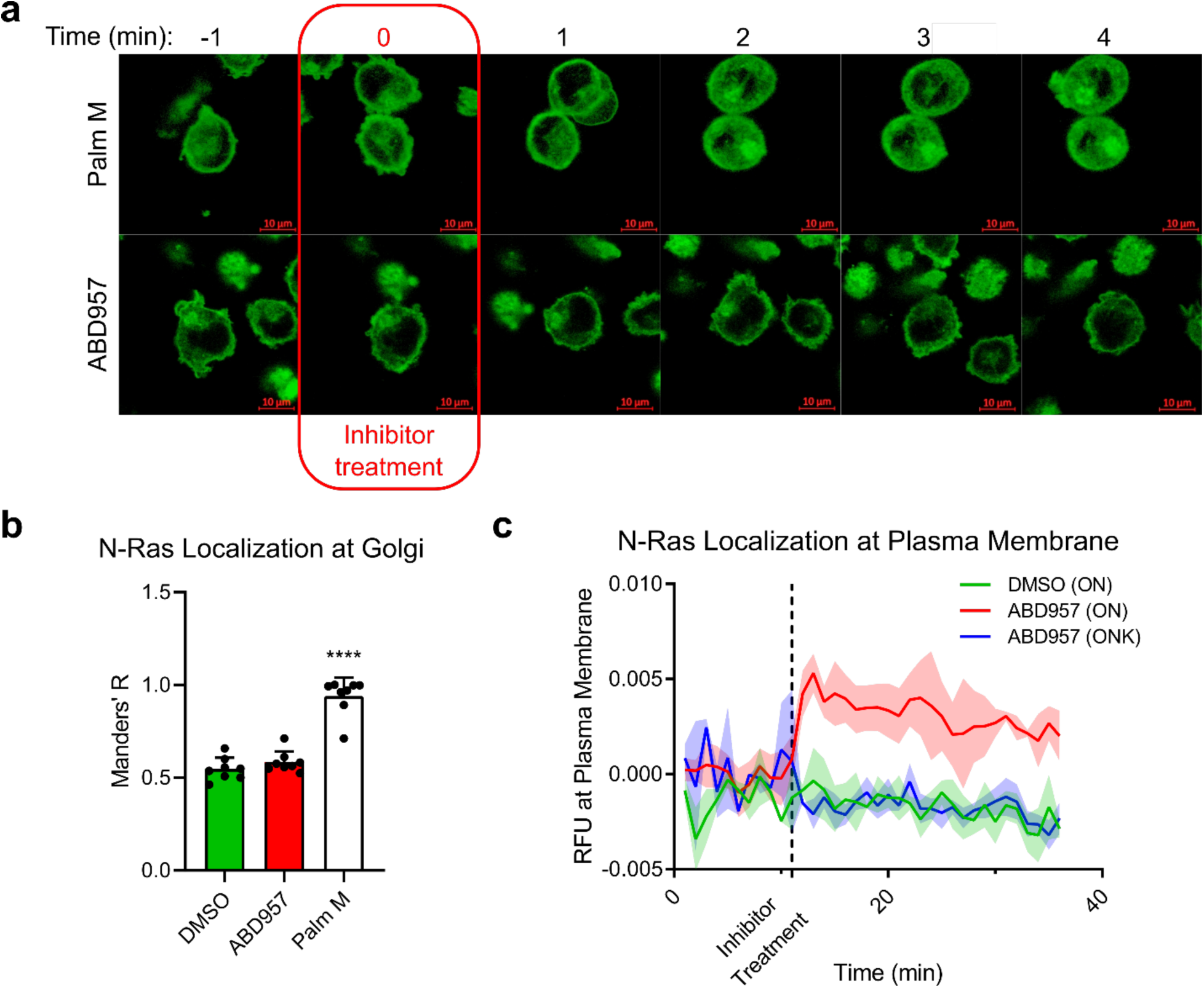
Effect of inhibitor treatment on GFP-N-Ras localization in ON and ONK cells. **a**, Time course images taken once every minute following administration of DMSO, Palm M (10 µM), or ABD957 (500 nM). Compound was infused into the medium at t = 0 (boxed in red). Green channel shows GFP-N-Ras. Note that only cells within the focal plane of the image collections were evaluated for N-Ras localization. Additional cells that were out of the focal plane are also seen in certain fields (identified by typically smaller sizes with entire cell showing green color). **b**, Fold-change in N-Ras localization to the Golgi apparatus, as a representative endomembrane compartment and measured by costaining with the Golgi marker N-acetylgalactosaminyltransferase (GALNT2), after 10-minute compound treatment. Data represent average values ± s.d. (n = 8 individual cells per group). Data shown are from representative experiments from three biological replicates. **c**, Quantification of GFP-N-Ras signal intensity at the plasma membrane in ON and ONK cells treated with ABD957 (500 nM). Data represent average values of individual cells ± s.d. as shaded area (n = 2-5 cells per image, with images collected each min). Data shown are from a representative experiment of five biological replicates.

### ABHD17 inhibition selectively blocks *NRAS* dependent cell growth and synergizes with the MEK inhibitor PD901

We next investigated how ABD957 affected N-Ras function and the growth of N-Ras-dependent cancer cells. We first addressed this question by measuring downstream signaling, specifically ERK phosphorylation, which we found was substantially blocked by both Palm M and ABD957 in *NRAS* mutant (and N-Ras-dependent) OCI-AML3 cells, but not in *KRAS* mutant (and N-Ras-independent) NB4 cells^57^ (**Fig. 5a, b**). In contrast, the MEK inhibitor, PD901, which functions downstream of Ras signaling, blocked ERK phosphorylation in both cell lines (**Fig. 5a, b**). The effect of Palm M on ERK phosphorylation was greater than ABD957, mirroring their respective impacts on N-Ras palmitoylation (**Fig. 2** and **3**).

**Figure 5.**
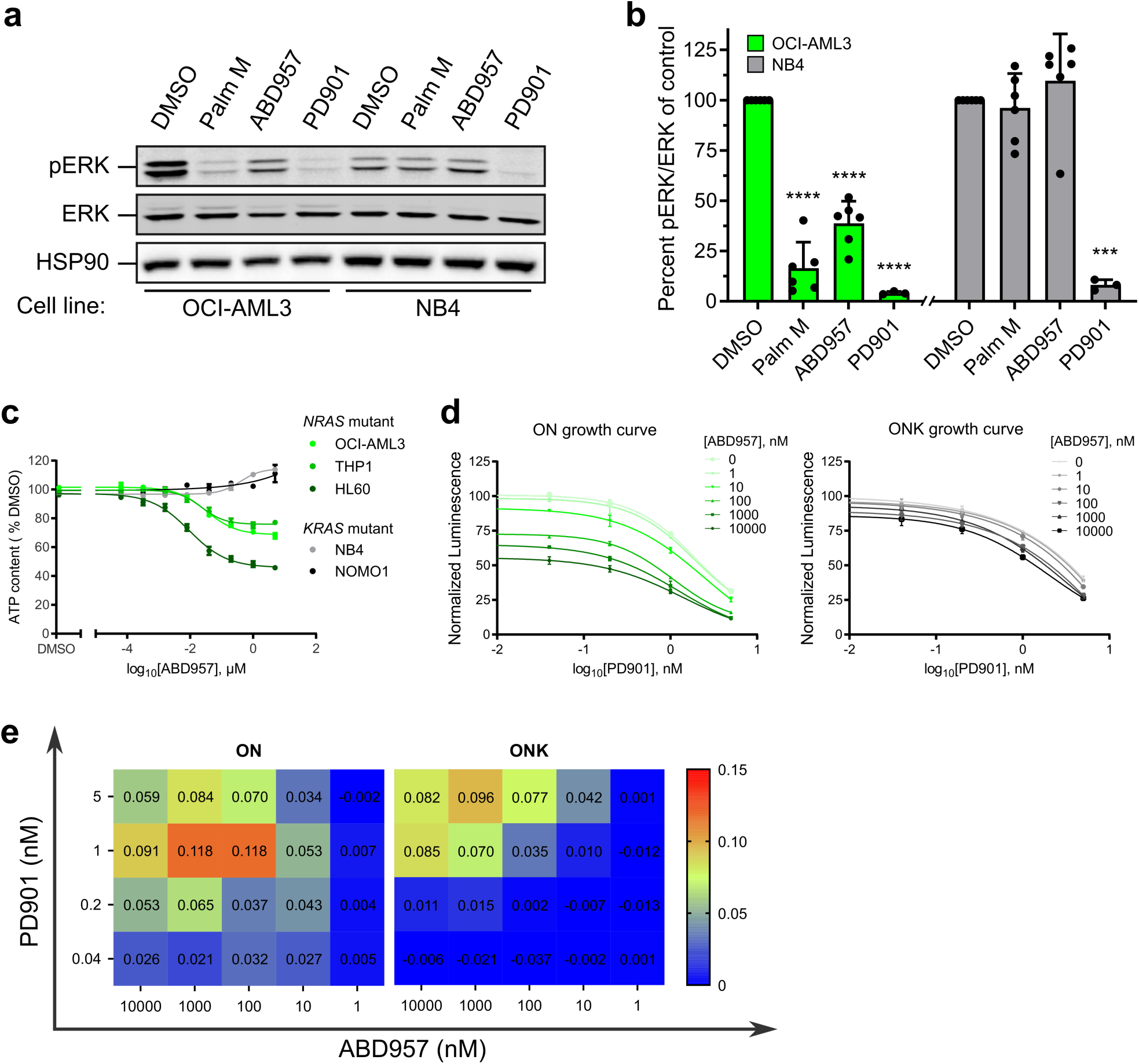
ABHD17 inhibition selectively impairs the signaling and proliferation of *NRAS* mutant leukemia cells. **a**, Representative western blot for phosphorylated ERK (pERK) following exposure to Palm M (10 µM), ABD957 (500 nM), or PD901 (10 nM) for 4 h in *NRAS* mutant OCI-AML3 cells or *KRAS* mutant NB4 cells. **b**, Quantification of data shown in **a**; average values ± s.d. (n = 6 independent experiments except for PD901, where n = 3 independent experiments). Statistical significance was calculated with unpaired two-tailed Student’s *t*-tests compared to DMSO control. **c, d**, Growth of the indicated human AML cell lines (**c**) and of isogenic ON and ONK cells (**d**) treated with varying concentrations of ABD957 for 72 h. Cell growth was measured by Cell Titer Glo. Data represent average values (n = 3 independent experiments). **e**, Bliss independence score quantifying synergy of ABD957 and PD901 in **d**.

The enhanced cellular stability of ABD957 (**Fig. 1d**) enabled further studies of its impact on cell proliferation, where we found that the compound reduced the growth of *NRAS* mutant AML cell lines (OCI-AML3, THP1, and HL60), but not *KRAS* mutant cell lines (NB4 and NOMO1) (**Fig. 5c**). The growth inhibitory effects of ABD957 were partial and plateaued at ∼500 nM (**Fig. 5c**) consistent with the near-complete inhibition of ABHD17 enzymes in AML cells treated with this concentration of the compound (see **Fig. 1f**). We next explored the impact of combining ABHD17 and MEK inhibitors by exposing ON and ONK cells to a range of ABD957 and PD901 concentrations. Using the Bliss Independence Model^58^ to assess potential drug interactions, we found that the combination of ABD957 and PD901 was substantially synergistic in ON cells, but not ONK cells (**Fig. 5d, e**), pointing to ABHD17 inhibition as a possible way to augment the therapeutic activity of MEK inhibitors in N-Ras-dependent cancers.

## Discussion

The fundamental role that C-terminal lipid modifications play in regulating Ras signaling has stimulated several historical efforts to target the post-translational processing of Ras onco-proteins^6, 59^. In the 1990s, potent and selective farnesyltransferase (FTase) inhibitors were developed; however, these compounds did not effectively block Ras modification or signaling due to alternative processing of N-Ras and K-Ras by geranylgeranyl transferases^60-64^. More recently, the importance of an additional lipid modification – dynamic cysteine *S*-palmitoylation – for certain Ras oncoproteins, including N-Ras, has pointed to an alternative set of pharmacological targets for controlling Ras signaling in cancer cells^7, 65^. Biochemical and pharmacological studies with first-generation inhibitors of N-Ras depalmitoylation have implicated several enzymes in this process^12^, with more recent studies revealing a possible key contribution from the ABHD17 sub-group of serine hydrolases^20^.

The high sequence identity shared by ABHD17A, ABHD17B, and ABHD17C (>70%) and their co-expression in most cell types presents technical challenges for studying the net contribution of these enzymes to N-Ras palmitoylation, which, to date, has not been assessed in *NRAS*-mutant cancer cells. We have addressed this problem by creating ABD957, a pan-ABHD17 inhibitor that displays good potency, proteomic selectivity, and metabolic stability. Using ABD957, we show that the ABHD17 enzymes make a substantial, but partial contribution to N-Ras depalmitoylation in *NRAS-*mutant AML cells, which is in agreement with previous work in heterologous cell systems showing that RNAi interference-mediated knockdown of ABHD17A/B/C failed to completely arrest N-Ras dynamic palmitoylation^20^. Whether other targets of the more promiscuous inhibitors Palm M and HDFP account for the putative ABHD17-independent N-Ras depalmitoylation activity remains unknown, but our pharmacological studies testing ABD957 in combination with JJH254 would argue against additional contributions from LYPLA1 and LYPLA2. We also note that our broader proteomic investigations of Palm M identified several proteins with heightened palmitoylation, including those that were not dynamically palmitoylated over the time course of our pulse-chase experiments. This result suggests that Palm M may affect other cellular metabolic processes beyond protein depalmitoylation that in turn impact the net palmitoylation state of proteins in cells.

One of our most striking findings was the selectivity that ABD957 displayed for N-Ras across the broader palmitoylated proteome. Unlike Palm M, which affected the palmitoylation state of many proteins, ABD957 only preserved the palmitoylation state of three plasma membrane-associated proteins – N-Ras, SCRIB, and MPP6. Considering that most of the other dynamically palmitoylated proteins affected by Palm M, but not ABD957, localize to intracellular membrane compartments (**Fig. 3e**), our data support a model where ABHD17 enzymes regulate palmitoylated proteins exclusively at the plasma membrane. We do not know yet whether individual ABHD17 enzymes (A, B, or C) will show differential substrate specificities in cells, but, regardless, it is likely that the substrate scope of this family of depalmitoylases will expand as more cell types are treated with ABD957 and examined for their palmitoylation profiles. Indeed, the palmitoylation of the plasma membrane-associated, post-synaptic density protein PSD-95 (DLG4), which is predominantly expressed in neurons and was not detected in the AML cells examined herein, has also been shown to be regulated by ABHD17 enzymes^20, 22^. Conversely, SCRIB has been found, in other cell types, to be regulated by the depalmitoylase LYPLA2^66^, indicating the potential for cellular context to also restrict the substrate scope of ABHD17 enzymes. With regards to other enzymes that may be responsible for regulating the palmitoylation state of intracellular proteins, we note that the serine hydrolase ABHD10 was recently identified as a depalmitoylase localized to the mitochondria^25^. Thus, emerging data point to the existence of multiple *S*-depalmitoylases that individually regulate dynamic protein palmitoylation in specific organelles of mammalian cells.

The general strategy of inhibiting the palmitoylation/depalmitoylation cycle of N-Ras has the advantage of selectively targeting cancer cells that depend on this oncoprotein for growth while preserving the functions of K-Ras4b in normal tissues. Our data indicate that ABHD17 inhibitors offer one potential way to achieve this objective. However, several important mechanistic questions about the interactions between ABHD17 enzymes and N-Ras palmitoylation and signaling remain to be answered. How, for instance, does ABD957-induced accumulation of palmitoylated N-Ras at the plasma membrane lead to perturbation in N-Ras signaling? Might, for instance, persistently palmitoylated N-Ras mislocalize, over time, to plasma membrane microdomains that are less productive for signal transduction? We also wonder whether the partial blockade of N-Ras signaling and cancer growth caused by ABHD17 inhibitors reflect maximal possible effects of the plasma membrane-delineated stabilization of N-Ras palmitoylation. Considering that Palm M more greatly affected N-Ras signaling (as measured by ERK phosphorylation), it is possible that redirecting N-Ras to intracellular membranes is required to more completely block signal transduction, an outcome that, based on our data, may not be achievable with ABHD17 blockade alone. On the other hand, the remarkably restricted impact of ABHD17 inhibitors on the palmitoylated proteome suggests that these compounds may perturb N-Ras signaling without gross effects on normal cell physiology. Such a profile may enable ABHD17 inhibitors to safely augment the activity of other anti-cancer agents, as we have shown for the MEK inhibitor PD901, to provide a compelling path for the targeted therapy of *NRAS* mutant cancers.

## Supporting information

Biology Methods

Chemistry Methods

## Contributions

J.R.R., R.M.S., N.A.Z., M.J.N., K.S., and B.F.C. conceived the project and wrote the paper. J.R.R. performed palmitoyl-proteomics experiments. R.M.S. perfomed NRAS palmitoylation assays. N.A.Z., J.R.R., A.J.F., A.I., A.L., M.P., and B.H. performed proliferation and NRAS signaling assays. T.W.H. performed imaging experiments. A.J.F. developed syngenic AML cell lines. N.N., C.L.H., and R.M.S. performed gel- and MS-ABPP experiments. N.N. and C.L.H. performed compound characterization. K.M.L. performed and analyzed MS-ABPP experiments and analyzed spectra. S.K.R. synthesized Palmostatin M. A.R.H. provided synthetic expertise and contributed to related studies not detailed in this paper. J.R.R., R.M.S., N.A.Z., T.W.H., K.M.L., and M.J.N. performed data analysis and visualization.

## Acknowledgements

This work was supported by grants from the NIH (CA193994, CA231991, and CA72614), an American Cancer Society postdoctoral fellowship PF-18-217-01-CDD (J.R.R.), the Leukemia and Lymphoma Society (LLS Fellowship 5465-18 to N.A.Z.), and the Damon Runyon Cancer Research Foundation (Fellowship DRG-2149-13 to A.J.F).

## Supplementary Inforrmation

Included are Supplementary Methods, **Supplementary Figures 1-6**, including full-length versions of cropped gels and blots shown in **Figures 1-5** and **Supplementary Figures 1-5, Supplementary Tables 1** and **2**.

---

**Supplementary Figure 1.**
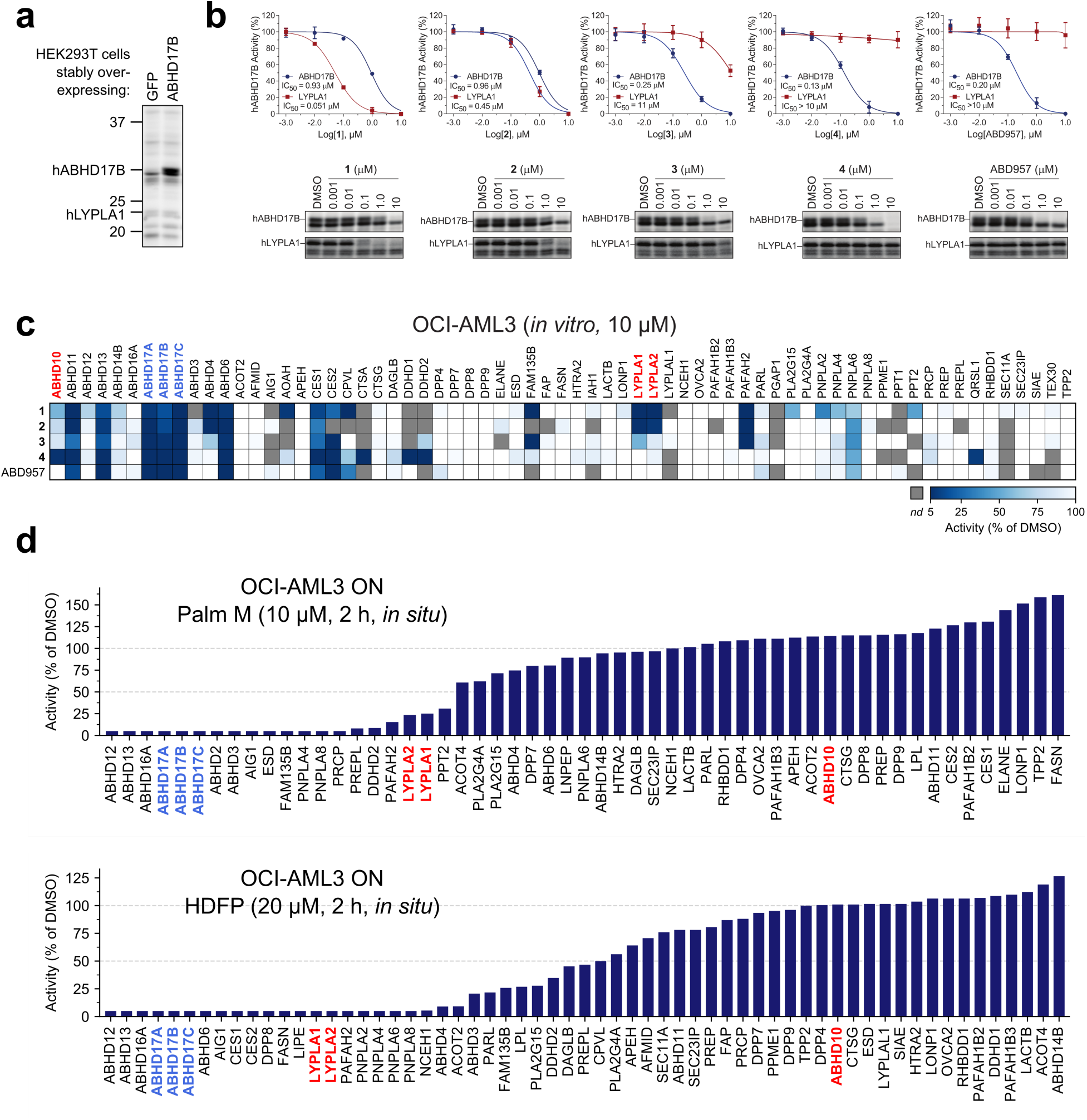
Characterization of ABHD17 inhibitors. **a**, Representative gel-ABPP image showing FP-rhodamine labeling of recombinant human ABHD17B (hABHD17B) in proteomic lysates of stably transfected HEK293T cells. **b**, IC_50_ curves and representative gel-ABPP images of human ABHD17B and LYPLA1 activity in HEK293T cell lysate treated with compounds **1**-**5**. Data represent average values ± s.d. (n = 3 independent experiments). **c**, MS-ABPP data of serine hydrolase activities in the particulate fraction of OCI-AML3 proteomes treated with compounds **1**-**5** (10 µM, 30 min). Data are from single experiments performed at the indicated concentrations for each compound. **d**, MS-ABPP data of serine hydrolase activities in OCI-AML3 ON cells treated *in situ* with Palm M (10 µM) or HDFP (20 µM) for 2 h. Data represent three experiments corresponding to independent treatments of cells with compound (Palm M or HDFP), after which cell lysates were combined together for analysis by MS-ABPP.

**Supplementary Figure 2.**
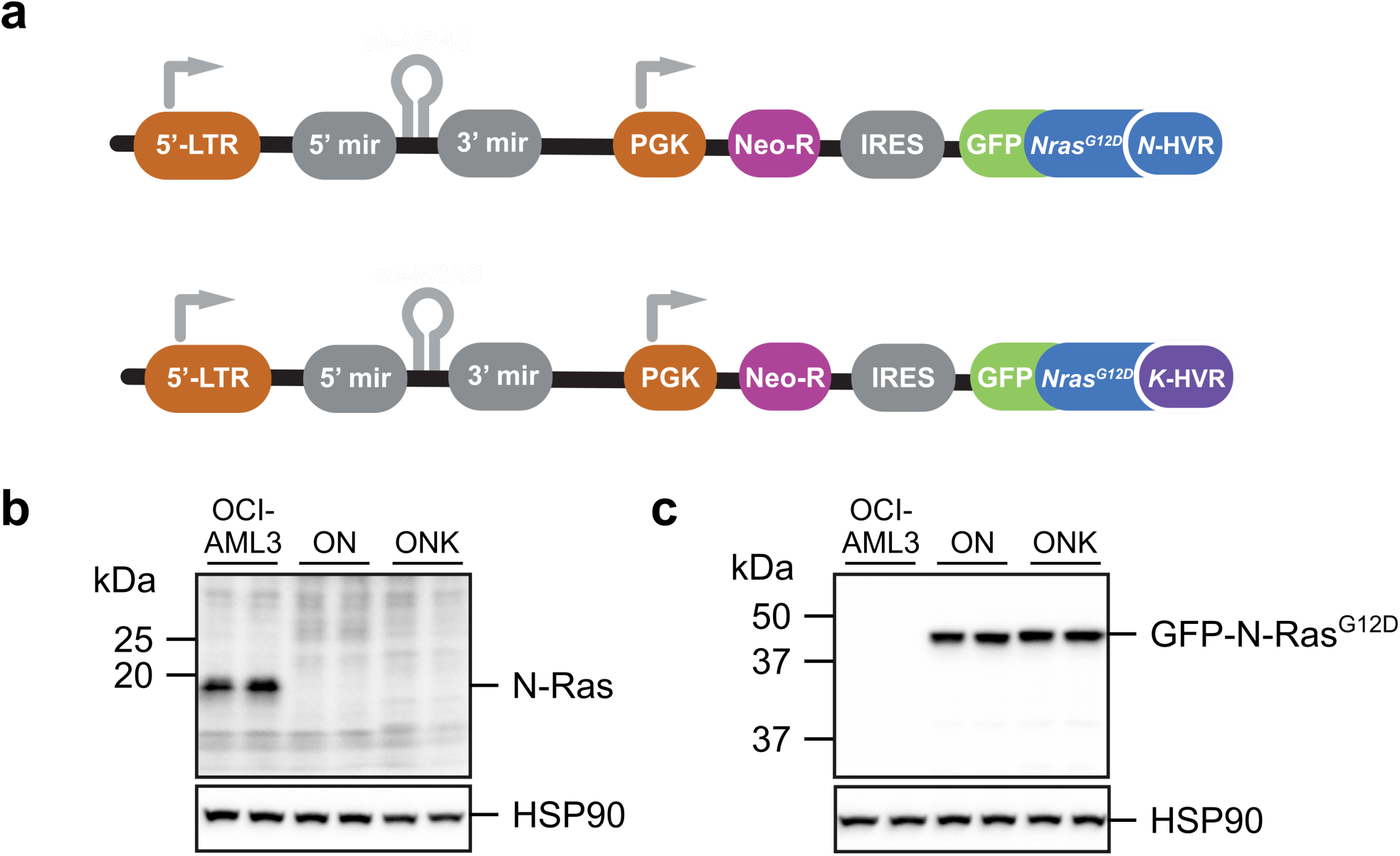
Establishment of OCI-AML3 sublines expressing GFP-N-RasG12D (ON cells) or GFP-N-Ras^G12D-KRAS-HVR^ (ONK cells). **a**, Schematic of viral construct containing an shRNA for N-Ras and cDNAs for GFP-N-Ras^G12D^ or GFP-N-Ras^G12D-KRAS-HVR^. **b**, Western blot confirming endogenous N-Ras knockdown in stably infected ON and ONK cells using N-Ras specific antibody^67^. **c**, Western blot analysis with an anti-Ras (G12D mutant) antibody confirming expression of GFP-N-Ras^G12D^ or GFP-N-Ras^G12D-KRAS-HVR^ in ON and ONK cells, respectively. For western blots, two biological replicates are shown for each group.

**Supplementary Figure 3.**
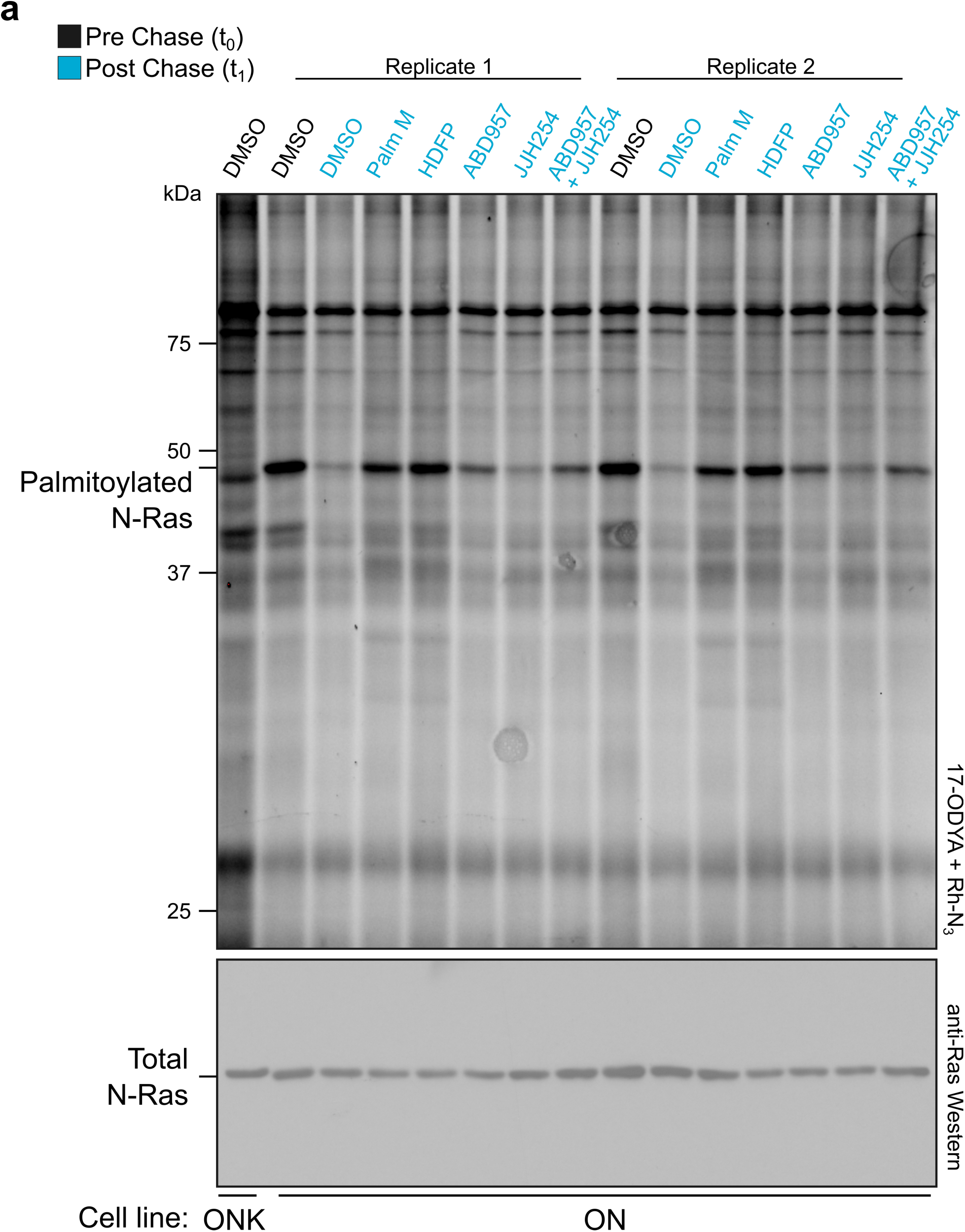
Effects of inhibitor treatment on global palmitoylation as assessed by SDS-PAGE and in-gel fluorescence scanning. Representative SDS-PAGE analysis of protein modification by 17-ODYA comparing effects of treatment with Palm M (10 µM), HDFP (20 µM), ABD957 (500 nM), and JJH254 (1 µM) in ON and ONK cells. Palmitoylation visualized by rhodamine attached via CuAAC to the alkyne of 17-ODYA (top gel). Total N-Ras quantity in each sample was visualized by western blotting with a Ras antibody following anti-GFP enrichments.

**Supplementary Figure 4.**
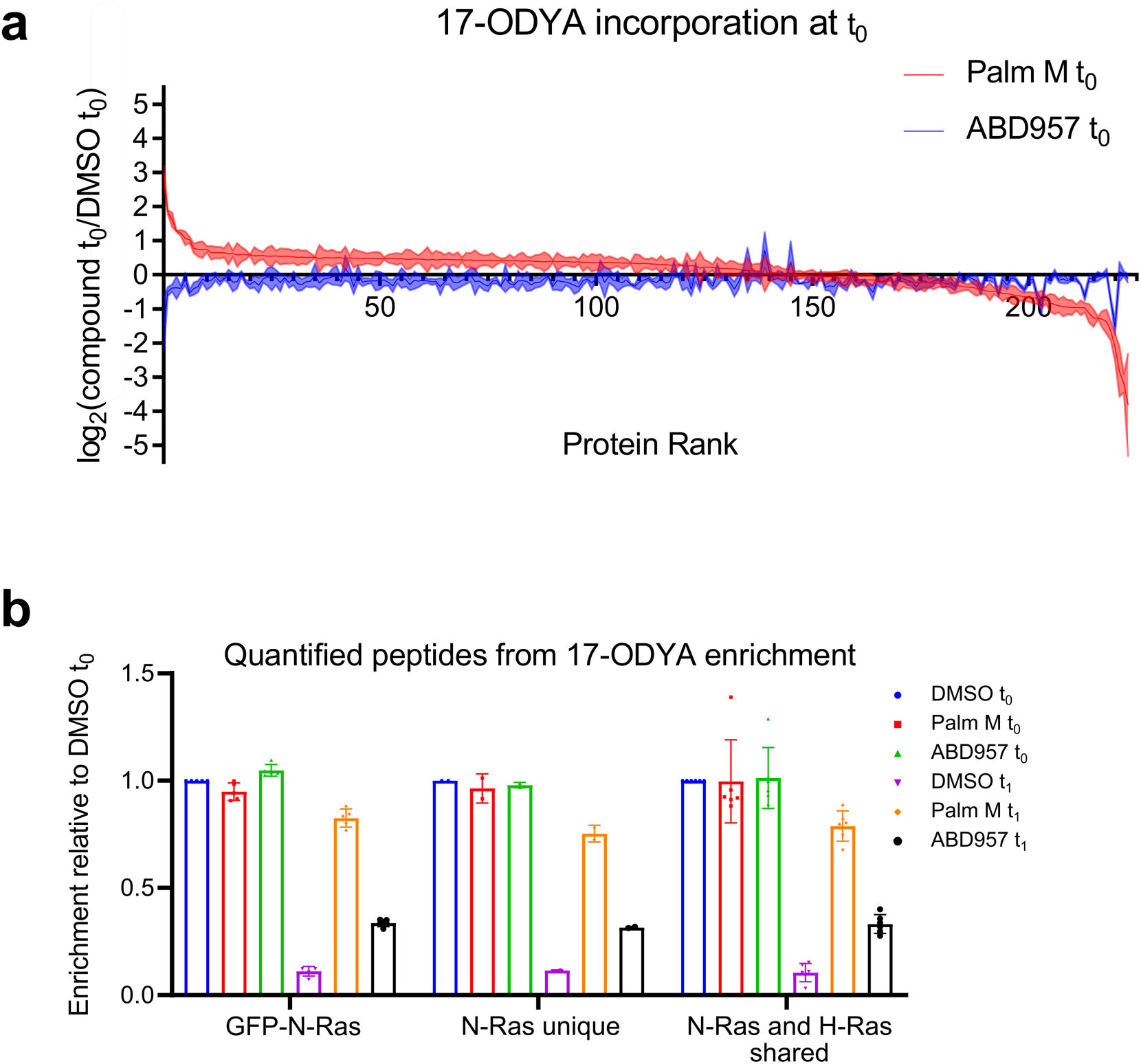
Global palmitoylation effects of Palm M and ABD957 in leukemia cells. **a**, MS-based proteomics of ON cells metabolically labeled with 20 µM 17-ODYA as described in **Fig. 3**, showing palmitoylated protein signals for Palm M (10 µM) or ABD957 (500 nM) treated cells versus DMSO-treated cells at t_0_. The results indicate that Palm M increases the apparent palmitoylation state of several proteins prior to the chase period and, as shown in **Fig. 3a**, some of these proteins are not dynamically palmitoylated (blue proteins in **Fig. 3a**). **b**, Bar graphs quantifying different categories of N-Ras peptides from MS-based proteomics experiments (also see **Supplementary Table 1**).

**Supplementary Figure 5.**
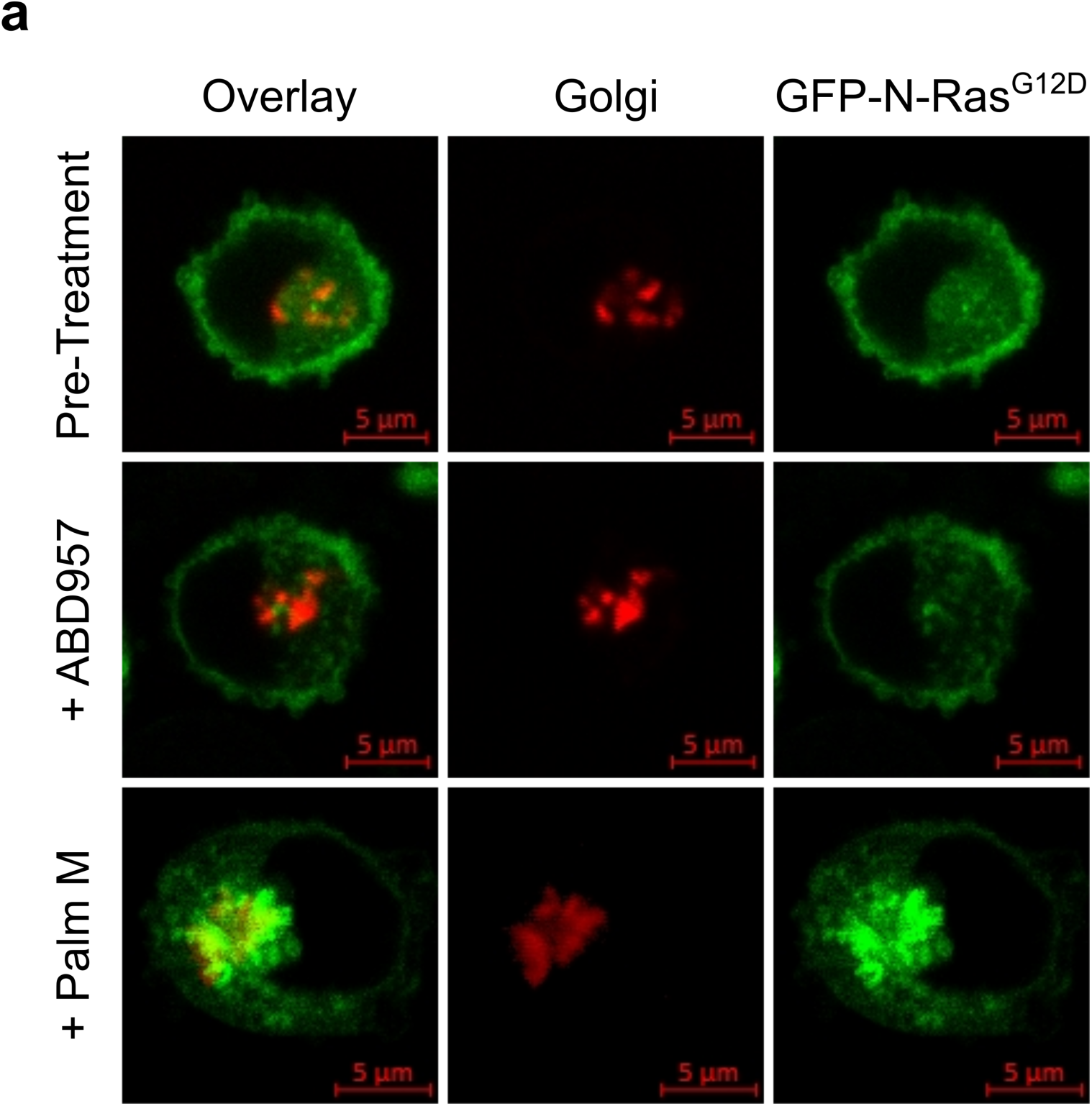
Partial colocalization of N-Ras with Golgi markers in Palm M-treated cells. Representative images from cells co-stained with the Golgi marker N-acetylgalactosaminyltransferase (GALNT2) before and after treatment with Palm M (10 µM) or ABD957 (500 nM). Red channel shows Golgi marker (middle), green channel shows GFP-N-Ras (right), and overlay of the two markers (left).

**Supplementary Figure 6.** Full-length gels and blots corresponding to the indicated cropped gels and blots in other figures.

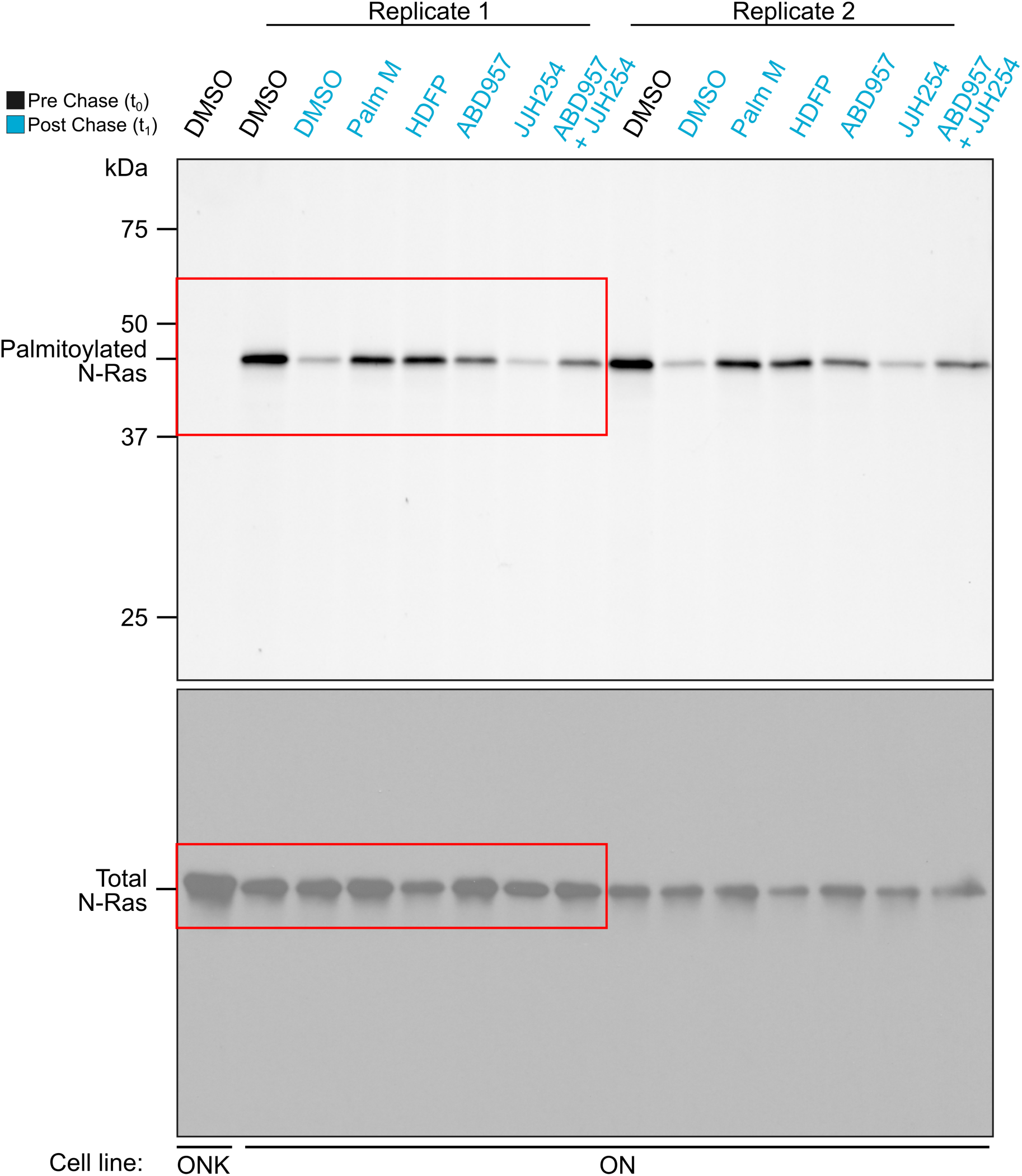

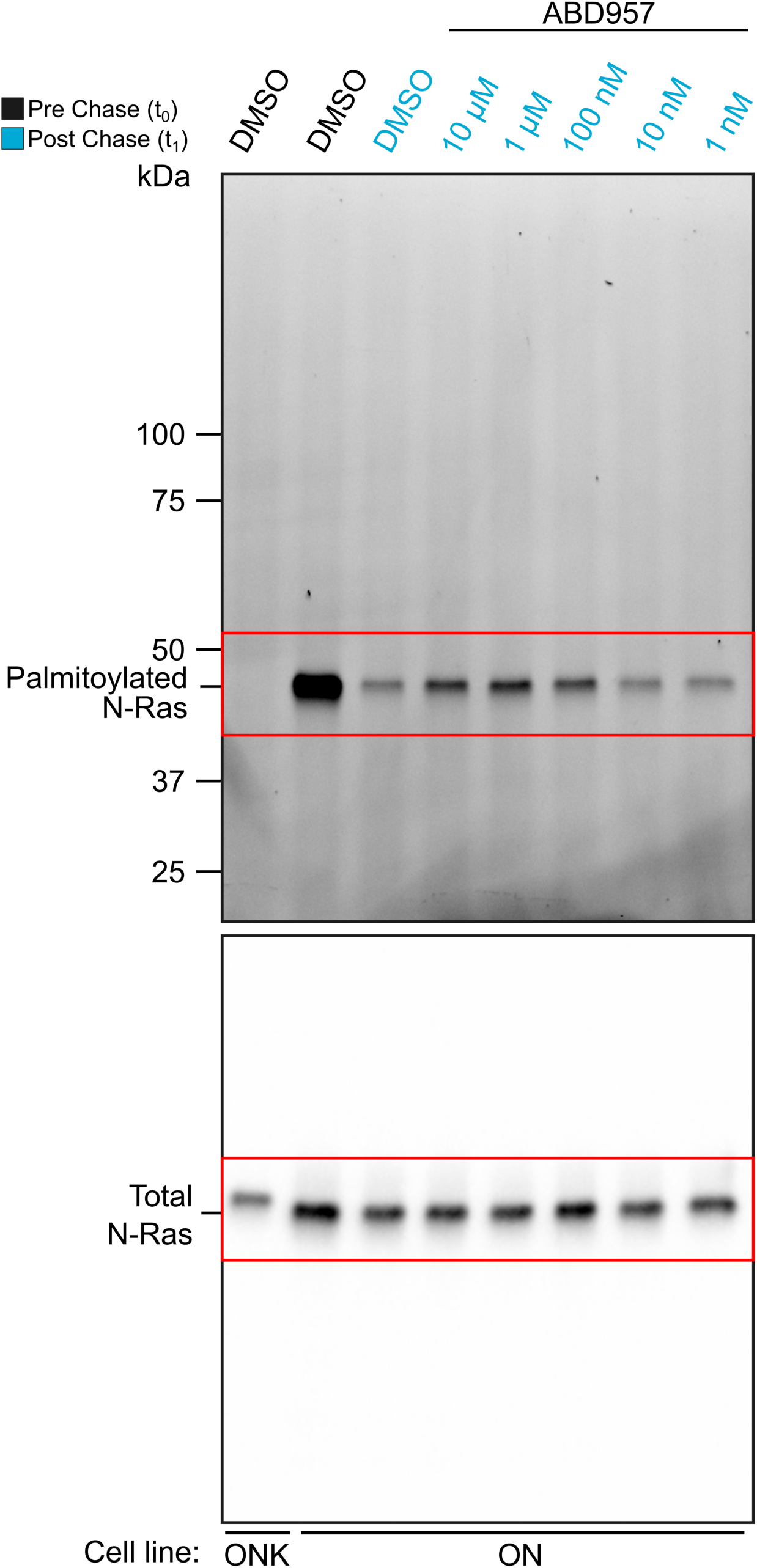

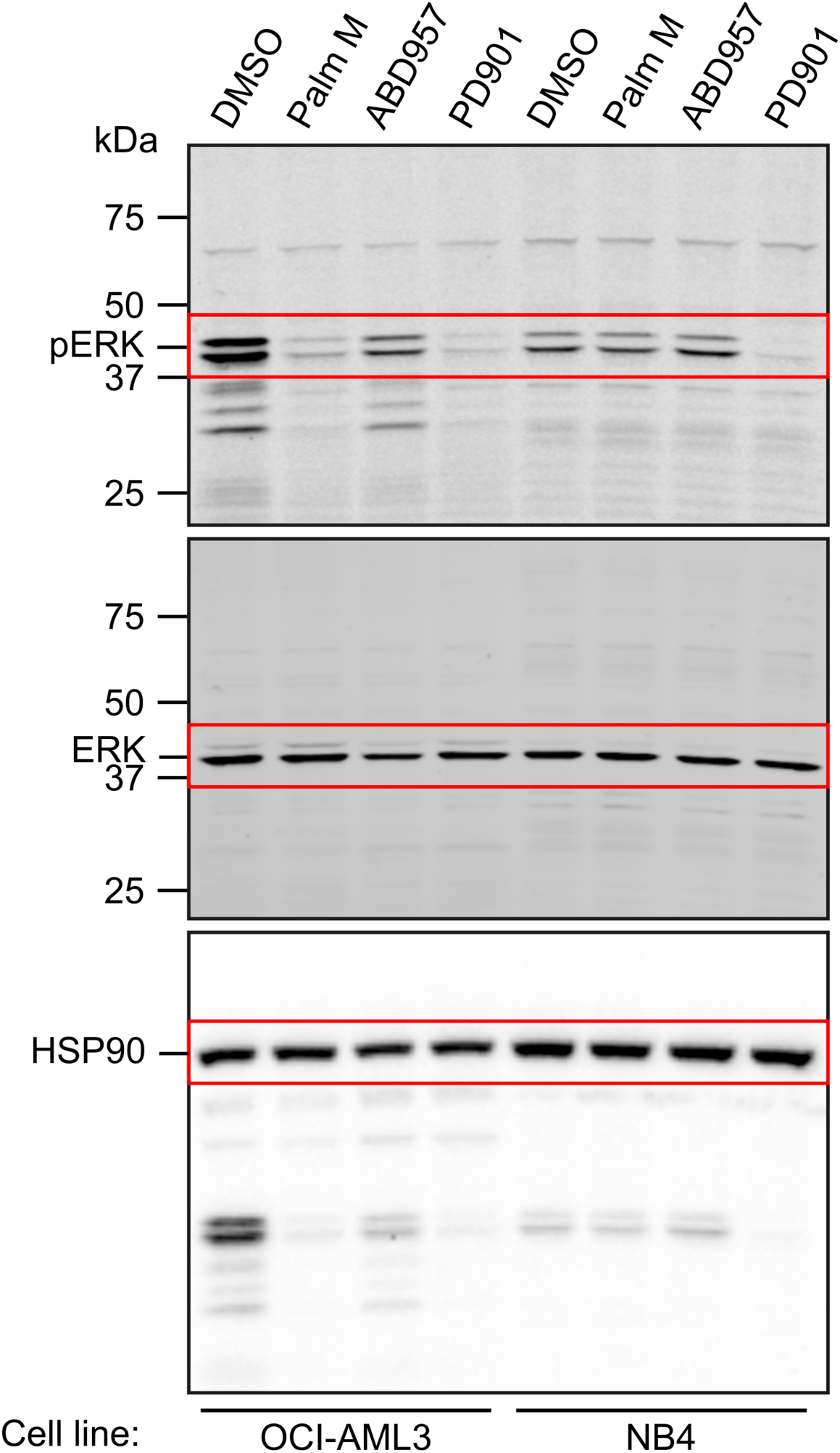

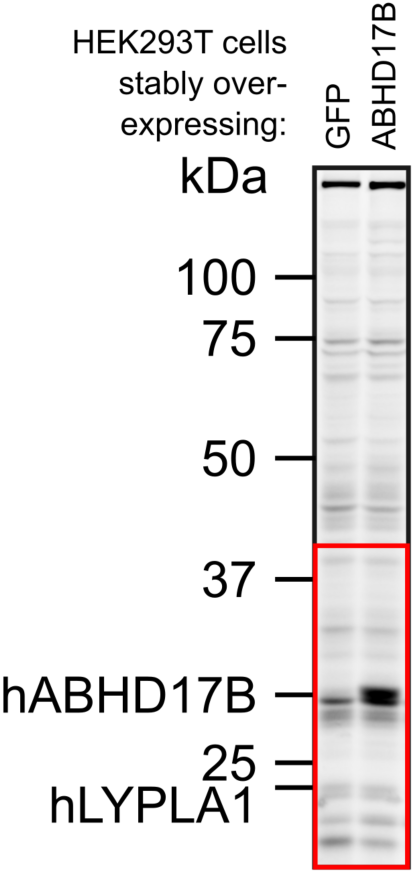

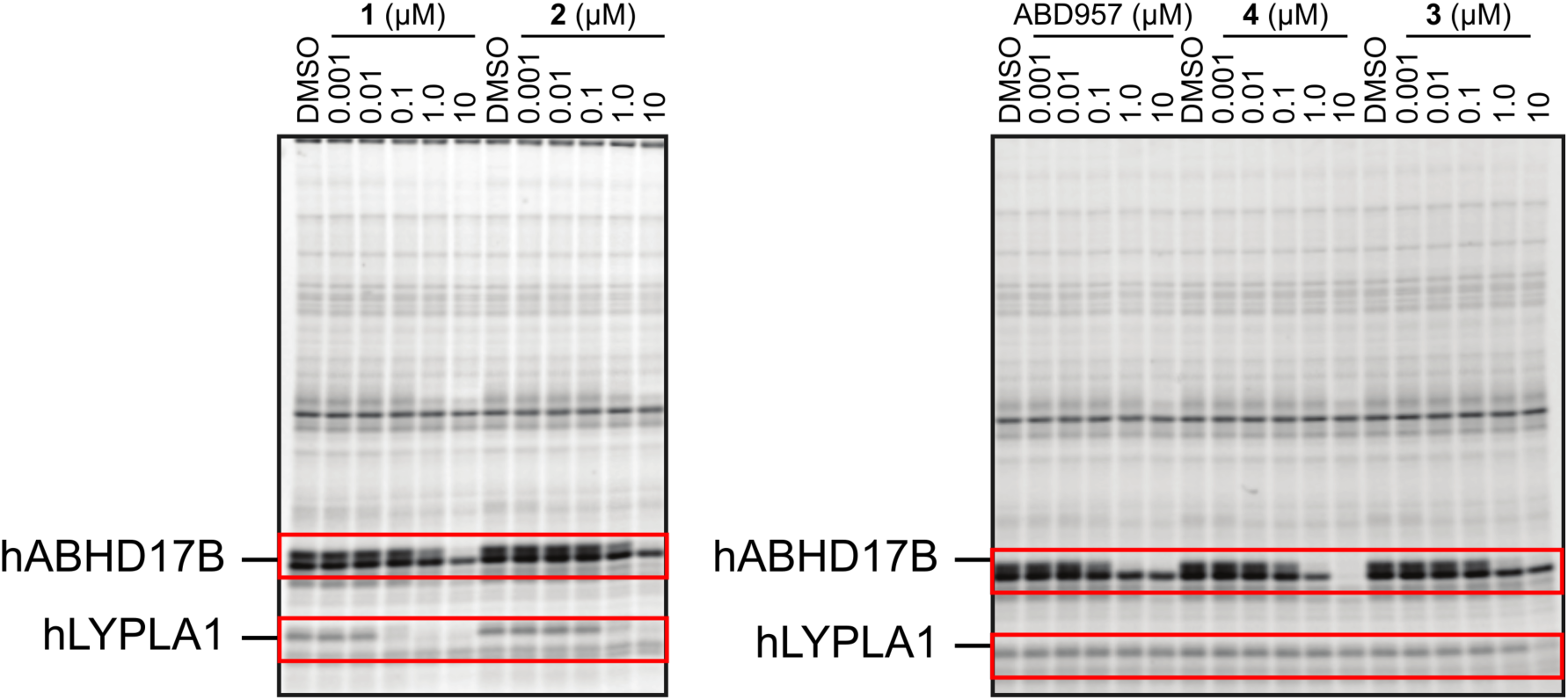

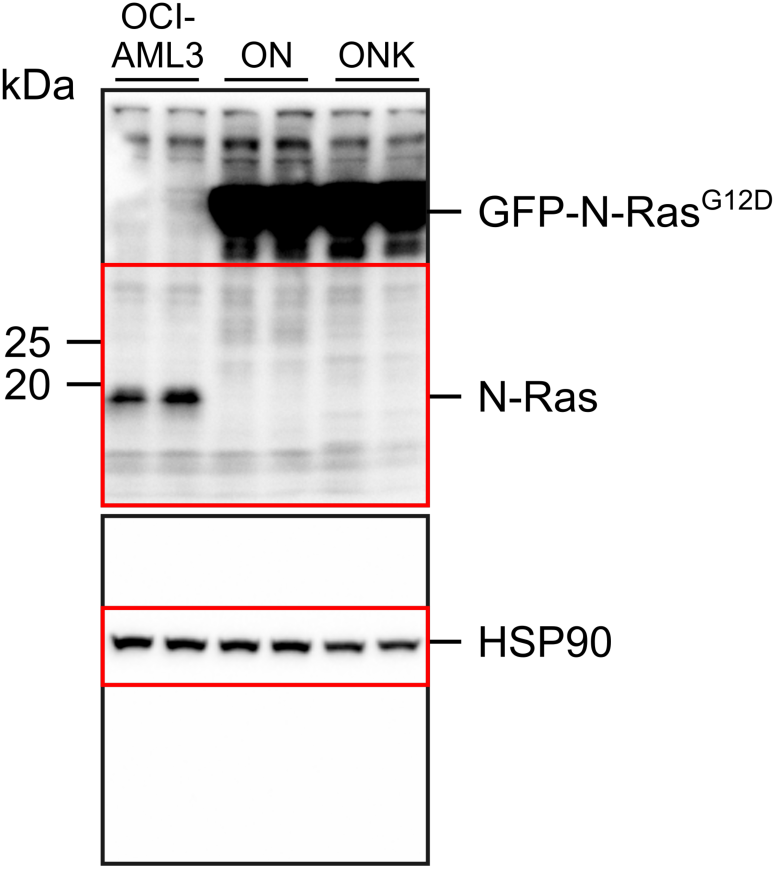

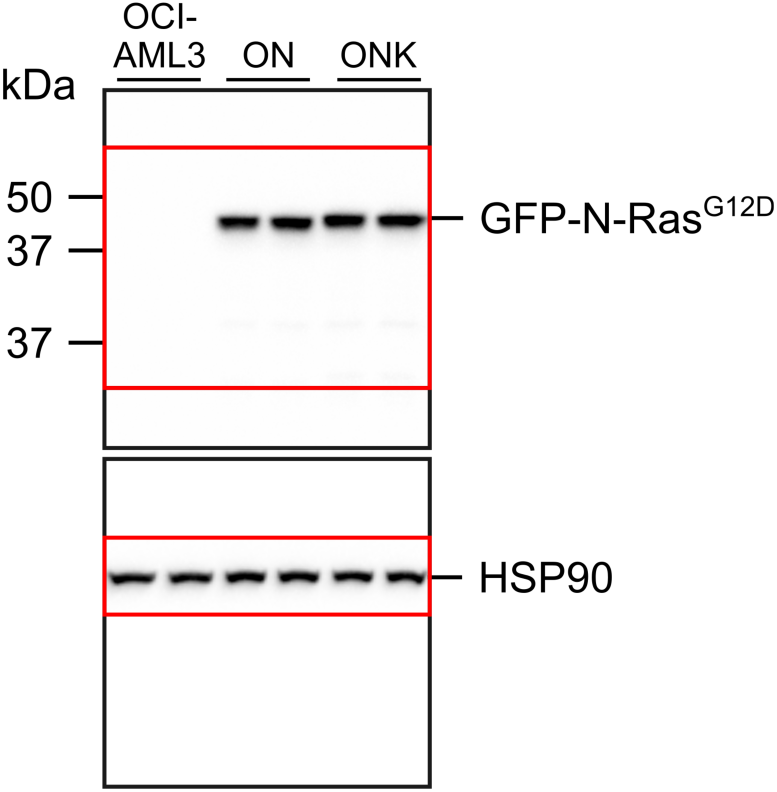

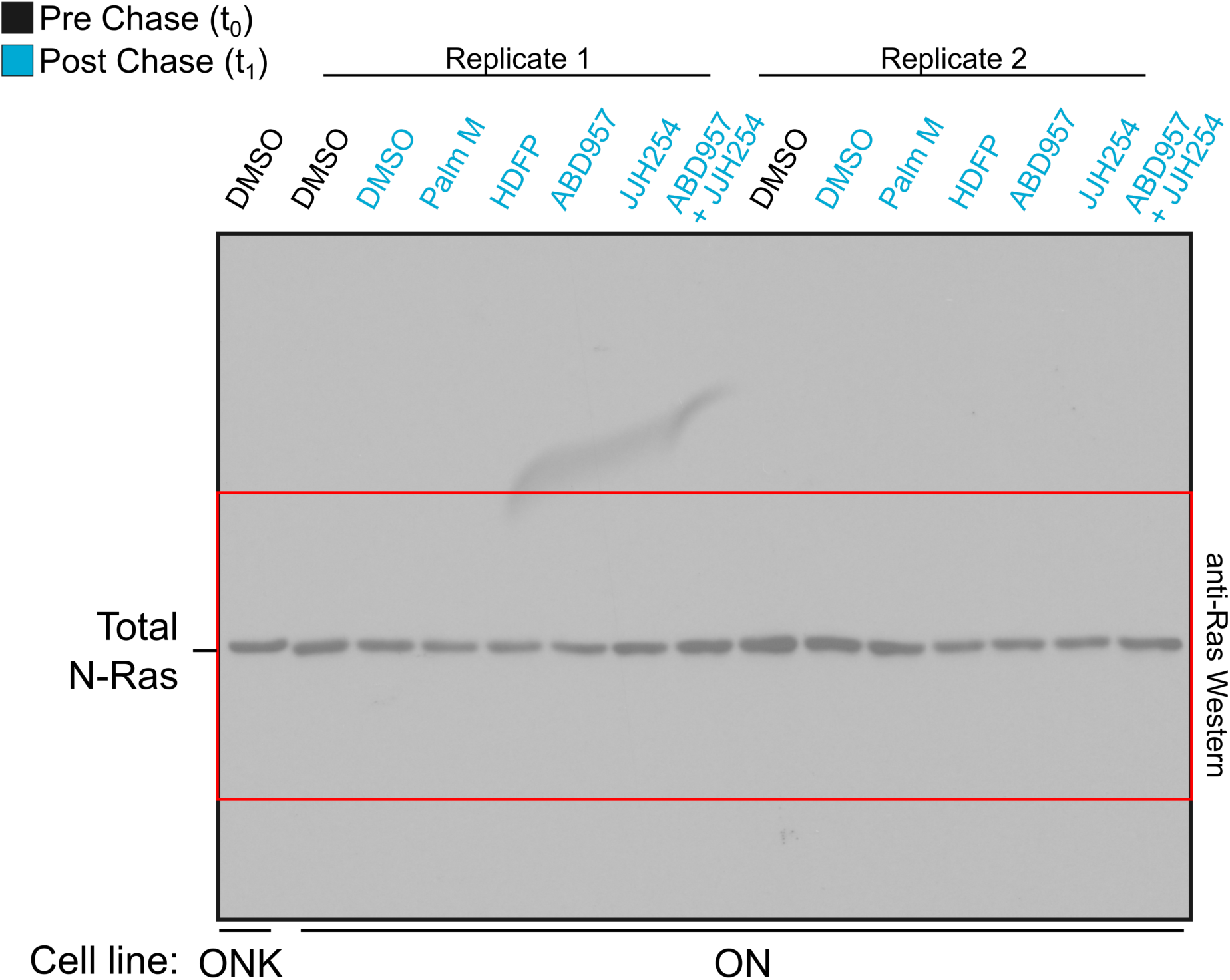

## Notes

### Competing Interest Statement

The authors have declared no competing interest.

