## Supplementary material for "ABHD17 enzymes regulate dynamic plasma membrane palmitoylation and N-Ras-dependent cancer growth": Biology Methods

**Cell lines and tissue culture**

OCI-AML3 (DSMZ catalog number: ACC-582) cells were grown in RPMI supplemented with 10% fetal bovine serum (FBS), L-glutamine (2 mM), penicillin (100 U/mL), streptomycin (100 μg/mL), and 50 μM β-mercaptoethanol and grown at densities between 0.3e6 and 2e6 cells/mL. NB-4 (DSMZ catalog number: ACC-207), NOMO1 (DSMZ catalog number: ACC-542), HL60 (DSMZ catalog number: ACC-3), and THP1 (ATCC: TIB-202) were grown in RPMI and HEK293T (ATCC: CRL-3216) were grown in DMEM, supplemented with 10% fetal bovine serum (FBS), L-glutamine (2 mM), penicillin (100 U/mL), and streptomycin (100 μg/mL). All cells were maintained at 37 °C with 5% $CO_2$.

ABHD17B (MHS1010-202726047, 4748883) clone in pCMD-SPORT6 vector was purchased from GE Dharmacon (now known as Dharmacon Inc).

The following primers were used for subcloning into an untagged pCL vector for viral transfection:

| **Primer Name** | **Sequence (5' to 3')** |
|---|---|
| FAM108B1_F_pCL_HindIII | TTT AAG CTT GCC ACC ATG AAT AAT CTT TCA TTT AGT G |
| FAM108B1_R_pCL_XhoI | AAA CTC GAG TTA CAA ATT TAC CAG TTC CTG |

**Generation of HEK293T transgenic lines**

HEK293T cells stably expressing various constructs were made using protocol described by Bar-Peled and Kemper.[1] In brief, 1.8e6 HEK293T cells were plated in 6 cm plates and allowed to settle overnight. To 200 μL of serum-free DMEM the following were added: 1 μg plasmid containing gene of interest in lentivirus vector, 900 ng ΔVPR, 100 ng VSV-G and 6 μL of XtremeGene HP transfection reagent. Reagents were flicked to mix, and after 15 minutes, the transfection mixture was added dropwise to plates containing cells. Virus containing supernatants were collected 48 hours later and concentrated using a Lenti-X Concentrator (Clontech), and then used to infect HEK293T cells (plated the day prior at 0.25e6 per well of 12-well plate) in the presence of 10 μg/mL Polybrene (Santa Cruz). The media was replaced 24 hours later, cells were allowed to recover for an additional 24 hours, and then puromycin was added for selection.

**Gel-based activity-based protein profiling**

Inhibitor potency ($IC_{50}$ values) against hABHD17B and hLYPLA1 was determined by competitive gel-based ABPP using FP-Rh competition[2].

Cell lysates were fractionated by ultra-centrifugation (100,000*g* for 45 min at 4ºC) and membrane pellets were resuspended in PBS and diluted to a final protein concentration of 1.0 mg/mL. Cell proteomes (50 μg) were treated with inhibitor (0.001-10 μM) or DMSO for 30 min at 37ºC and subsequently treated with FP-Rh (1.0 μM) for an additional 30 min at room temperature. Reactions were quenched with 4X SDS-PAGE loading buffer and FP-Rh-labeled enzymes were resolved by SDS-PAGE (10% acrylamide). In-gel fluorescence was visualized using a Bio-Rad ChemiDoc™ XRS imager. Fluorescence is shown in gray scale. Quantification of enzyme activities was performed by densitometric analysis using ImageJ software (NIH). Integrated peak intensities were generated for bands corresponding to ABHD17B and LYPLA1. $IC_{50}$ values were calculated through curve fitting semi-log-transformed data (*x*-axis) by non-linear regression with a four-parameter, sigmoidal dose response function (variable slope) in Prism software (GraphPad).

**MS-ABPP sample preparation**

For *in situ* treatments, OCI-AML3, THP1 or ON cells were resuspended in fresh media at 2e6 cells/mL and treated with DMSO, Palm M (10 µM), HDFP (20 µM), or ABD957 (500 nM or 1 µM) and incubated for the indicated times. Cells were pelleted at 500 RCF by centrifugation, washed with PBS, pelleted again and frozen.

Frozen cell pellets were thawed, diluted in PBS and subsequently lysed by probe sonication. The membrane fraction of each cell sample was isolated using ultra-centrifugation (100,000*g* for 45 min at 4ºC) and membrane pellets were resuspended in PBS and diluted to 2.0 mg/mL. Membrane proteomes (2 mg/mL in 1 mL of PBS) were labeled with FP-biotin (10 µM) for 1 h at room temperature while rotating. After labeling, the proteomes were denatured and precipitated using 4:1 $MeOH/CHCl_3$, resuspended in 0.5 mL of 6 M urea in PBS, reduced using tris(2-carboxyethyl)phosphine (TCEP, 10 mM) for 30 min at 37 °C, and then alkylated using iodoacetamide (40 mM) for 30 min at room temperature in the dark. The biotinylated proteins were enriched with PBS-washed avidin-agarose beads (100 µL; Sigma-Aldrich) by rotating at room temperature for 1.5 h in PBS with 0.2% SDS (6 mL). The beads were then washed sequentially with 5 mL 0.2% SDS in PBS (3x), 5 mL PBS (3x) and 5 mL $H_2O$ (3x). On-bead digestion was performed using sequencing-grade trypsin (2 µg; Promega) in 2 M urea in 100 mM triethylammonium bicarbonate buffer with 2 mM $CaCl_2$ for 12–14 h at 37 °C (200 µL). Duplex reductive dimethylation (ReDiMe) was performed as previously described[3]. Briefly, for duplex ReDiMe of WT versus KO tissues either $^{13}CD_2O$ (heavy) or $CH_2O$ (light) was added to each sample (0.15%) followed by addition of $NaBH_3CN$ (22.2 mM). Tissue samples from vehicle- or inhibitor-treated samples were labeled using a triplex ReDiMe protocol where control samples (i.e. vehicle-treated) were labeled with $CH_2O$ (light) and inhibitor-treated samples labeled with $^{13}CD_2O$ (heavy). Light samples were then treated with $NaBH_3CN$ whereas heavy samples were treated with $NaBD_3CN$. Following a 1 h incubation period at room temperature, the reaction was quenched by addition of $NH_4OH$ (0.23%) and formic acid (0.5%). The samples were then combined and analyzed by LC/MS/MS.

**MS-ABPP data analysis**

Nanoflow LC-MS/MS measurements were performed on an Ultimate 3000 (Thermo Scientific) interfaced with an Orbitrap Fusion Lumos Tribrid mass spectrometer (Thermo Scientific) via an EASY-Spray source (Thermo Scientific). Peptides were separated on an EASY-Spray PepMap RSLC C18 column (2 µm particle size, 75 µm × 50 cm; Thermo Scientific, ES801) heated to 55 °C using a flow rate of 400 nl/min. The compositions of LC solvents were A: water and 0.1 % formic acid, and B: 95% acetonitrile, 5% water and 0.1% formic acid. Peptides were eluted over 4 hours using the linear gradient, 2.5-215 min 3-35% B, 215-230 min 25-40% B, 230-231 min 45-70% B, 231-233 min 70-90% B, 233-234 min 5-70% B, 234-236 min 70-90% B, 236-240 min 3% B.

MS data were acquired in data dependent mode (top 20 dependent scans). MS1 profile scans were acquired in the Orbitrap (resolution: 120,000, scan range: 375–1500 m/z, AGC target: 4.0e5, maximum injection time: 50 ms). Monoisotopic peak determination was set to “peptide”. Only charge states 2-5 were included. Dynamic exclusion was enabled (repeat count, *n*: 1, exclusion duration: 60 s, mass tolerance: ppm, low: 10 and high: 10, excluding isotopes). An intensity threshold of 5e3 was set. MS2 spectra were acquired in centroid mode. Precursor ions were isolated using the quadrupole (isolation window: 1.6 m/z), fragmented using CID (collision energy: 35%, activation time: 10 ms, activation Q: 0.25), and detected in the ion trap (scan range mode: auto m/z normal, scan rate: rapid, AGC target: 4.0e3, maximum injection time: 300 ms, injecting ions for all available parallelizable time).

Spectrum raw files were extracted into MS2 files using RawConverter (http://fields.scripps.edu/rawconv/) and searched using the ProLuCID algorithm[4] against a human reverse concatenated nonredundant Uniprot database, with static modifications for cysteine residues to account for alkylation by iodoacetamide (+57.0215 m/z), and standard static modifications for reductive dimethylation: lysine and N-terminus (+28.0313 m/z for light, +34.06312 m/z for heavy). Data was assembled using DTASelect version 2.0,[5] and ratio quantification was performed using in-house CIMAGE software.[6] Peptides were required to be fully tryptic, unique, to have an envelope correlation score of $R^2 \geq 0.5$, and ratios were capped to a maximum value of 20. Proteins were required to have at least two unique peptides, and in cases where proteins had exactly one peptide with a calculated ratio of 20, and at least one other peptide with a ratio below 2, the 20 value was discarded.

#### Syngeneic ON/ONK cells

A pCDH-LMN-GFP lentiviral vector was obtained by cloning the miR30-PGK-NeoR-IRES-GFP cassette from LMN-GFP[7] into a pCDH Expression Lentivector (System Biosciences). A miR30-based shRNA targeting human *NRAS* (sense: 5'- CAGGGTGTTGAAGATGCTTTT -3') was cloned into the vector. The coding sequence of $\textit{Nras}^{G12D}$ was cloned downstream of GFP to create a N-terminal GFP-fused N-$\text{Ras}^{\text{G12D}}$ expression construct with N-Ras endogenous HVR (N-HVR). Alternatively, a chimeric version was cloned were the sequence corresponding to amino acids 166-188 from N-Ras was replaced with that of K-Ras-4B (K-HVR). Lentiviral vectors production and transduction of OCI-AML3 cells were performed as previously described[8]. ON and ONK cells were validated by Western Blot.

#### Dynamic palmitoylation assay

Dynamic palmitoylation assay was modeled after earlier work in our lab[9]. OCI-AML3 cells were grown as described above, then spun down (3 min, 500 RCF) and re-suspended in fresh medium at a density of 2e6 cells/ml (5-10 million cells per sample for gel-based assay and 20e6 cells per sample for mass spectrometry-based assay). Cells were pre-incubated with inhibitor or DMSO for 1 hour, then 17-ODYA (20 μM) was added for 1 hour. Samples were pelleted and snap frozen immediately following the 17-ODYA "pulse" and designated $t_0$, or re-suspended in pre-warmed chase medium which consisted of OCI-AML3 growth media, supplemented with DMSO or inhibitor at the same concentration as in the pre-incubation step and incubated for 1 hour. Cells were then centrifuged, placed on ice, washed in cold PBS, snap frozen, and designated $t_1$.

#### Immunoprecipitation

Cell pellets were re-suspended in cold lysis buffer which consisted of: 1% Triton-X PBS with 1 mM PMSF, 0.2 mM HDSF, 20 μM HDFP and protease inhibitors (complete Ultra EDTA-free mini tablets, 5892791001 Roche), sonicated with a microtip probe sonicator (7 times, 50% rate, power 4), and placed on an end-over-end rotator for 30 minutes at 4 °C. Samples were then hard spun (16,300 RCF, 10 minutes) and the supernatant was transferred to fresh tubes on ice. Protein concentration was measured using a DC assay kit from Bio-Rad and adjusted to 1 mg/mL. Input samples were taken at this point and stored at -80°C.

Enrichment was performed with anti-GFP Sepharose beads (20 μL 50:50 slurry, ab69314, abcam). Antibody conjugated beads were spun down (500 RCF), storage buffer was aspirated with a 26-gauge needle, and beads were washed three times with 1% Triton-X PBS. To each sample, 20 μL of washed bead slurry was added using a cut pipette tip, and samples were placed on an end-over-end rotator for 3 hours at 4 °C. Next, washes were performed by centrifugation/aspiration (3x, 500 RCF) with cold 1% Triton-X PBS containing 500 mM NaCl. After the last wash, supernatant was removed, and a 26-gauge needle was quickly inserted into the bead slurry to remove all remaining liquid. GFP beads were then re-suspended in 50 μL wash buffer. At this point, IP samples were either stored at -80°C or processed further for analysis by SDS-PAGE gel.

**Click chemistry and processing for SDS-PAGE gel**

On-bead click chemistry was performed to conjugate a rhodamine fluorophore reporter to 17-ODYA labeled proteins. To each 50 μL sample, 6 μL of a click chemistry reaction mixture was added. Click reaction mixture was freshly prepared as follows (amounts given are per sample): 1 μL 50 mM $CuSO_4$ (in water; final concentration during reaction of 1 mM), 3 μL 1.7 mM TBTA (4:1 tert-butanol:DMSO; 100 μM final), 1 μL 50 mM TCEP (freshly made in PBS; 1 mM final) and 1 μL Rh-$N_3$ (DMSO; 25 μM final). Reactions were allowed to proceed for 1 hour at room temperature. Samples were then quenched with 4x SDS loading buffer containing 1% BME (described below), boiled for 5 minutes to effect elution and finally 17-ODYA proteins were resolved and imaged on gel.

Loading buffer was prepared as follows (for 100 mL): 3.02 g Tris base was added to 40 mL water. 40 mL of glycerol was added slowly, and the mixture was stirred using a stir bar while the pH was brought to 6.75 using concentrated HCl. 8 g of SDS were then added followed by 20 mL of water and a pinch of bromophenol blue. Loading buffer was stored at room temperature, and 1% (10 μL per 1 mL buffer) β-mercaptoethanol (BME) was added immediately before quenching.

Finally, samples were loaded onto a 10% acrylamide gel, resolved by SDS-PAGE (300V for approx. 2.5 hours, and in-gel fluorescence was visualized using a Bio-Rad ChemiDoc MP flatbed fluorescence scanner.

**Preparation of tandem mass tag (TMT) labeled 17-ODYA enrichment samples**

Samples were prepared as previously described[9], with the following modifications. 17-ODYA labeled ON cells were lysed in DPBS containing protease inhibitors (Roche) and 1 mM PMSF using a Branson Sonifier probe sonicator (3 rounds of 8 pulses, 40% duty cycle, output setting = 4). Protein concentration was determined by DC assay on a microplate reader, and 500 μL of whole cell lysate at 2 mg/mL was added to a mixture of TBTA (30 μL/sample, 1.7 mM in 1:4 DMSO:t-BuOH), $CuSO_4$ (10 μL/sample, 50mM in $H_2O$), TCEP (10 μL/sample, 50mM in DPBS) and Biotin-N3 (10 μL/sample, 10 mM in DMSO). After 1 h, samples were chloroform-methanol precipitated. Hydroxylamine-sensitivity was performed as previously described from ON cells metabolically labeled with 17-ODYA for 2 h[9]. Following precipitation, each sample was solubilized in 1.2% SDS in DPBS via sonication and diluted to 0.2% SDS for enrichment.

Streptavidin beads (Thermo Scientific) 100 µL slurry was added to each sample and rotated at room temperature for 2 h, washed three times with 0.2% SDS, three times with DPBS, and three times with water. Beads were resuspended in 6 M urea in 200 mM EPPS, reduced with 10 mM neutral TCEP for 30 min at 37°C followed by addition of 20 mM iodoacetamide for 30 min at room temperature. Samples were diluted to 2 M urea in 200 mM EPPS, pelleted, and resuspended with 200 µL 2 M urea in 200 mM EPPS containing 2 µg sequence grade porcine trypsin (Promega) and 1 mM $CaCl_2$ and digested overnight at 37°C. Digests were centrifuged to remove the beads, and acetonitrile was added to reach 30% final volume. 6 µL (20 µg/µL) of respective 6-plex TMT tag (Thermo Scientific) was added to digests and incubated at room temperature for 1 h with occasional vortexing. Labeling reaction was quenched with 6 µL of 5% hydroxylamine for 15 minutes, followed by acidification by 15 µL formic acid. Samples were combined and vacuum-centrifuged to dryness to remove acetonitrile, reconstituted in Buffer A (95% $H_2O$, 5% acetonitrile, 0.1% formic acid) and stored at -80°C until MS analysis.

**Mass spectrometry analysis of tandem mass tag (TMT) labeled peptides**

Labeled peptides were prepared for MS analysis as previously described[10]. Briefly, labeled peptides were pressure loaded onto a 250 µm (inner diameter) fused silica capillary column packed with 4 cm C18 resin (Phenomenex, Aqua 5 µm). Samples were analyzed on an Orbitrap Fusion mass spectrometer (Thermo Scientific) coupled to an UltiMate 3000 Series Rapid Separation LC system and autosampler (Thermo Scientific Dionex). Peptides were separated on a 100 µm inner diameter capillary column with a 5 µm tip packed with 10 cm C18 (Phenomenex, Aqua 5 µm) and 3 cm strong cation exchange resin (SCX, Phenomenex)] using a 5-step 'MudPIT' protocol that injects 5 µL 0%, 20%, 50%, 80%, 100% salt bumps of ammonium acetate (500mM) in buffer A (100% $H_2O$, 0.1% formic acid) followed by an increasing gradient of buffer B (100% acetonitrile, 0.1% formic acid) in Buffer A in each step. A MS3-based TMT method was used for data acquisition. MS1 full scan spectrum (resolution: 120,000, scan range: 400−1700 m/z, RF lens 60%, AGC target 2e5, maximum injection time 50 ms, centroid mode) with dynamic exclusion enabled (repeat count 1, duration 15s) was performed. The top ten precursors were then selected for MS2/MS3 analysis. MS2 analysis consisted of quadrupole ion trap analysis, AGC 1.8e4, CID collision energy 35%, Activation Q 0.25, maximum injection time 120 ms, and isolation window at 0.7 m/z. Synchronous Precursor Selection (SPS) was enabled to include up to 10 MS2 fragment ions for the MS3 spectrum. MS3 precursors were fragmented by HCD and analyzed using the Orbitrap (resolution: 15,000, collision energy 55%, AGC 1.5e5, maximum injection time 120 ms). For MS3 analysis, we used charge state–dependent isolation windows. For charge state z = 2, the MS isolation window was set at 1.2 m/z; for z = 3-6, the MS isolation window was set at 0.7 m/z.

Raw files were processed using MaxQuant (v1.6.3.3) using the Homo sapiens reviewed Uniprot FASTA database (October 2018) with N-Ras substituted for GFP-N-$Ras^{G12D}$ to reflect the syngeneic ON cells, with FDR <1% at the peptide spectrum match (PSM) and protein levels, and removed identifications from reverse and MaxQuant contaminants database. Required minimum of 2 unique peptides for identification per MS experiment, included only unique peptides and minimum MS3 reporter ion intensity of 5,000 per control channel (DMSO $t_0$) for each PSM. Relative protein abundance was averaged across all PSMs.

### Proliferation Assay

A 40mM stock solution of ABD957 was serially diluted in DMSO to obtain 2000X concentrations. Next, twice-concentrated medium (2X) was obtained by diluting the drugs 1000-fold in culture medium. A solvent control was obtained by diluting DMSO 1000-fold in culture medium. To test the effect of ABD957 on cellular growth, cells were resuspended in fresh culture medium to a concentration of 6e5 cells/ml and seeded in a 96-well plate (75 µl/plate). Subsequently, 75 µl of 2X drugged medium or control was added. Each treatment was performed in triplicate. After 72 h in culture, 50 µl of each well were transferred to a white, flat-bottom 96 well plate and an equal volume of Cell-TiterGlo reagent was added. After 30 min of shaking at room temperature, luminescence was analyzed using a Tecan Infinite M200PRO plate reader. Raw values were normalized using Prism GraphPad software.

### Western Blot Analysis

A minimum of 2e6 cells per condition were pelleted and resuspended in fresh medium at 1e6 cells/mL containing compounds at the indicated concentration. Compound containing media were obtained by diluting a 1000-fold concentrated drug stock or vehicle (DMSO control) in fresh medium. Unless otherwise indicated, cells were treated for 4 hours, and then pelleted by centrifugation (500 RCF), washed in PBS, and frozen. Cells were resuspended in RIPA buffer (Thermo Scientific) supplemented with PhosSTOP (Sigma-Aldrich) and protease inhibitors (Roche) and lysed using a Branson Sonifier probe sonicator (3 rounds of 8 pulses, 40% duty cycle, output setting = 4). Western blotting was performed as previously described[11]. Briefly, lysates were separated on SDS-PAGE TG gels, and transferred to nitrocellulose membranes at 350 mA for 90 minutes, block and probe with primary and secondary antibodies in 5% milk in 1% TBS-T. Antibodies for western blotting included anti-pan-Ras (Cell Signaling, 3965, 1:500), anti-N-RAS (Santa Cruz, sc-31, 1:1000), anti-Ras (G12D specific mutant) (Cell Signaling, 14429, 1:1000), anti-pERK (Cell Signaling, 4370, 1:1000), anti-ERK (Cell Signaling, 9107, 1:2500), anti-HSP90-HRP (Cell Signaling, 79641, 1:5000), anti-mouse HRP (Cell Signaling, 7076, 1:10,000), anti-rabbit HRP (Cell Signaling 7074 or Santa Cruz sc2030, 1:10,000), Li-cor IRDye 800CW Donkey anti-rabbit (1:10,000), Li-cor IRDye 680RD Goat anti-mouse (1:10,000). Blots were imaged on a Li-cor Odyssey (Model 9120) (pERK and ERK) or detected using chemiluminescence on a Bio-Rad ChemiDoc™ XRS imager. Densitometry analysis was performed using ImageJ software (NIH).

### Live-cell imaging

OCI-AML3 cells were transferred to a 12 mm Nunc glass bottomed dish and diluted with 200 uL medium so that the entire 12 mm glass bottom was covered. Cells were then treated with Cell Light Golgi or ER RFP 2.0 reagent and incubated overnight. The following day, dishes were transferred to a ZEISS LSM 880 laser scanning confocal microscope equipped with an incubator equilibrated to 37°c and 5% $CO_2$. For live cell studies, images were acquired every 60 seconds over five z planes using 25 mW Multi-line Ar laser with a C-Apochromat 63x/1.2 W autocorr M27 Oil objective for 72 cycles. Images were quantified using CellProfiler. Briefly, cellular objects were identified based on GFP signal using standard minimum cross entropy, and Golgi or endoplasmic reticulum were identified based on RFP signal with a similar approach. From these objects, edge intensity of the GFP signal was used to quantify plasma membrane associated N-Ras, and total endomembrane signal was quantified by measuring total intercellular GFP signal of each cellular object. For colocalization studies, Manders' R was calculated from GFP and RFP signal in cells expressing GALNT2-RFP with a 20% threshold of maximum signal intensity.

**Statistical analysis**

Quantitative data expressed in bar and line graphs with mean ± s.d. (standard deviation, shown as an error bar) shown. Differences between two groups were examined using an unpaired two-tailed Student's t-test. Significant P values were indicated (*P < 0.05, **P < 0.01, ***P < 0.001 and ****P < 0.0001).
