## Supplementary material for "ABHD17 enzymes regulate dynamic plasma membrane palmitoylation and N-Ras-dependent cancer growth": Chemistry Methods

### Chemistry General Methods

All commercially available chemicals were obtained from Aldrich, Acros, Fisher, Fluka, Maybridge, or the like and were used without further purification. Palm M and HDFP were synthesized as generally described[1, 2]. Anhydrous solvents and oven-dried glassware were used for synthetic transformations sensitive to moisture and/or oxygen. All reactions are typically carried out under an inert nitrogen atmosphere using oven-baked glassware unless otherwise noted. Flash chromatography is performed using 230–400 mesh silica gel 60 using Isco Combiflash instruments. NMR spectra were generated on either Bruker 300 or Bruker 400 instruments. Chemical shifts are typically recorded in ppm relative to tetramethylsilane (TMS) with multiplicities given as s (singlet), bs (broad singlet), d (doublet), t (triplet), dt (double of triplets), q (quadruplet), qd (quadruplet of doublets), hept (heptuplet), and m (multiplet). Chemical purities were >95% for all final compounds as assessed by LC/MS analysis with detection at 210 and 254 nm.

**N,N-dimethyl-1-(4-(2-morpholino-4-(trifluoromethyl)benzyl)piperazine-1-carbonyl)-1H-pyrazole-4-sulfonamide (2).**

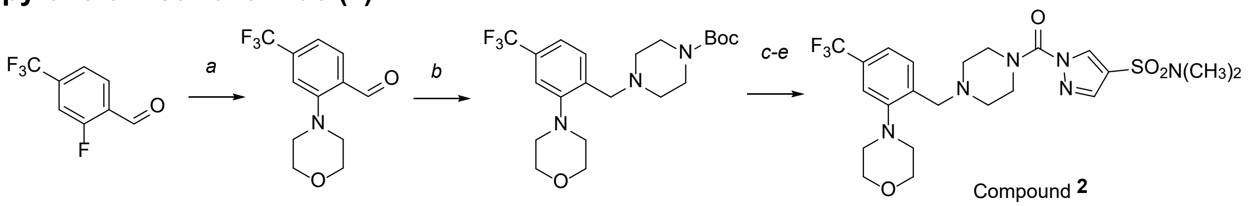

Compound **2**

*Step A: 2-morpholino-4-(trifluoromethyl)benzaldehyde*

A 100-mL round-bottom flask was charged with 2-fluoro-4-(trifluoromethyl)benzaldehyde (400 mg, 2.08 mmol, 1.00 equiv) in DMSO (10 mL). Morpholine (271 mg, 3.11 mmol, 1.50 equiv), potassium carbonate (861 mg, 6.18 mmol, 3.00 equiv) was added under nitrogen. The resulting solution was stirred overnight at 80 °C. The reaction progress was monitored by LCMS and upon completion the reaction was quenched with water (10 mL). The resulting solution was extracted with EtOAc (3 x 10 mL) and the organic layers were combined, washed with brine (2 x 10 mL), dried over anhydrous $Na_2SO_4$, filtered and concentrated under reduced pressure. The residue was chromatographed on a silica gel column with EtOAc/petroleum ether (1/5) to yield 223 mg (41% yield) of 2-(morpholin-4-yl)-4-(trifluoromethyl)benzaldehyde as yellow oil. LCMS (ESI, *m/z*): 260 $[M+H]^+$ .

*Step B: tert-butyl 4-(2-morpholino-4-(trifluoromethyl)benzyl)piperazine-1-carboxylate*

A 100-mL round-bottom flask was charged with 2-(morpholin-4-yl)-4-(trifluoromethyl)benzaldehyde (223 mg, 0.860 mmol, 1.00 equiv) in 1,2-dichloroethane (10 mL), tert-butyl piperazine-1-carboxylate (240 mg, 1.29 mmol, 1.50 equiv), triethylamine (260 mg, 2.57 mmol, 3.00 equiv) was added. The resulting solution was stirred for 30 min at room temperature. Sodium triacetoxyborohydride (547 mg, 2.58 mmol, 3.00 equiv) was added. The resulting solution was stirred overnight at room temperature. The reaction progress was monitored by LCMS and upon completion the reaction was then quenched with water (10 mL). The resulting solution was extracted with DCM (3 x 10 mL) and the organic layers were combined, washed with brine (2 x 10 mL), dried over anhydrous $Na_2SO_4$, filtered and concentrated under reduced pressure. The residue was chromatographed on a silica gel column with DCM/methanol (20/1) to yield 300 mg (81% yield) of tert-butyl 4-[[2-(morpholin-4-yl)-4-(trifluoromethyl)phenyl]methyl]piperazine-1-carboxylate as yellow oil. LCMS (ESI, *m/z*): 430 $[M+H]^+$ .

*Step C: 4-[2-(piperazin-1-ylmethyl)-5-(trifluoromethyl)phenyl]morpholine*

A 100-mL round-bottom flask was charged with tert-butyl 4-[[2-(morpholin-4-yl)-4-(trifluoromethyl)phenyl]methyl]piperazine-1-carboxylate (300 mg, 0.700 mmol, 1.00 equiv) in DCM (10 mL), trifluoroacetic acid (2.50 mL). The resulting solution was stirred for 5 h at room temperature. The reaction progress was monitored by LCMS and upon completion the resulting mixture was concentrated under reduced pressure to yield 230 mg (100% yield) of 4-[2-(piperazin-1-ylmethyl)-5-(trifluoromethyl)phenyl]morpholine as yellow oil and was carried to the next step without further purification. LCMS (ESI, *m/z*): 330 $[M+H]^+$ .

*Step D: 4-[[2-(morpholin-4-yl)-4-(trifluoromethyl)phenyl]methyl]piperazine-1-carbonyl chloride*

A 40-mL round-bottom flask was charged with triphosgene (104 mg, 0.350 mmol, 0.50 equiv) in DCM (10 mL), 4-[2-(piperazin-1-ylmethyl)-5-(trifluoromethyl)phenyl]morpholine (230 mg, 0.700 mmol, 1.00 equiv), N-ethyl-N-isopropylpropan-2-amine (181 mg, 1.40 mmol, 2.00 equiv) was added. The resulting solution was stirred for 3 h at 0 °C. The reaction was then quenched with water (10 mL). The resulting solution was extracted with DCM (3 x 10 mL) and the organic layers were combined, washed with brine (2 x 10 mL), dried over anhydrous $Na_2SO_4$, filtered and concentrated under reduced pressure to yield 300 mg (crude) of 4-[[2-(morpholin-4-yl)-4-(trifluoromethyl)phenyl]methyl]piperazine-1-carbonyl chloride as yellow oil and was carried to the next step without further purification. LCMS (ESI, *m/z*): 392 $[M+H]^+$ .

*Step E: N,N-dimethyl-1-(4-(2-morpholino-4-(trifluoromethyl)benzyl)piperazine-1-carbonyl)-1H-pyrazole-4-sulfonamide (**2**)*

A 40-mL round-bottom flask was charged with 4-[[2-(morpholin-4-yl)-4-(trifluoromethyl)phenyl]methyl]piperazine-1-carbonyl chloride (300 mg, 0.770 mmol, 1.00 equiv) in tetrahydrofuran (10 mL), N,N-dimethyl-1H-pyrazole-4-sulfonamide (161 mg, 0.920 mmol, 1.20 equiv), N,N-diisopropylethylamine (297 mg, 2.30 mmol, 3.00 equiv), 4-dimethylaminopyridine (9.36 mg, 0.0800 mmol, 0.10 equiv) was added. The resulting solution was stirred overnight at 60 °C. The reaction progress was monitored by LCMS and upon completion the reaction was then quenched with water (10 mL). The mixture was extracted with DCM (3 x 10 mL) and the organic layers were combined, washed with brine (2 x 10 mL), dried over anhydrous $Na_2SO_4$, filtered and concentrated under reduced pressure. The crude product (710 mg) was purified by preparative HPLC using the following gradient conditions: 30% $CH_3CN$/80% Phase A increasing to 80% $CH_3CN$ over 10 min, then to 100% $CH_3CN$ over 0.1 min, holding at 100% $CH_3CN$ for 1.9 min, then reducing to 30% $CH_3CN$ over 0.1 min, and holding at 30% for 1.9 min, on a Waters 2767-5 Chromatograph. Column: Xbridge Prep C18, 19*150mm 5um; Mobile phase: Phase A: aqueous $NH_4HCO_3$ (0.05%); Phase B: $CH_3CN$; Detector, UV220 & 254nm. Purification resulted in 118.0 mg (29% yield) of *N,N*-dimethyl-1-[(4-[[2-(morpholin-4-yl)-4-(trifluoromethyl)phenyl]methyl]piperazin-1-yl)carbonyl]-1H-pyrazole-4-sulfonamide as a white solid.

$^1$H NMR (300MHz, $CDCl_3$) δ 8.47 (s, 1H), 7.82 (s, 1H), 7.61 - 7.66 (m, 1H), 7.31 - 7.40 (m, 2H), 3.77 - 3.92 (m, 8H), 3.67 (br, 2H), 2.87 - 3.02 (m, 4H), 2.74 (s, 6H), 2.63 (br, 4H).

$^{13}$C NMR (101 MHz, DMSO-$d_6$) δ 152.36, 149.20, 139.96, 136.85, 134.12, 131.10, 128.50 (q, $J$ = 31.5 Hz), 124.23 (q, $J$ = 272.3 Hz), 119.68 (q, $J$ = 3.5 Hz), 117.88, 116.07 (q, $J$ = 3.6 Hz), 66.50, 56.25, 52.46, 52.33, 37.46.

HRMS calculated for $C_{22}H_{29}F_3N_6O_4S$ $[M+H]^+$ 531.1996, found 531.2003.

**1-(4-(4-chlorobenzyl)piperazine-1-carbonyl)-N,N-dimethyl-1H-pyrazole-4-sulfonamide (1).**

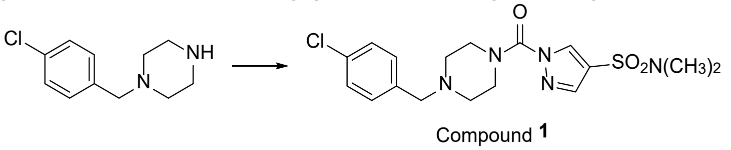


Compound **1**

1-(4-chlorobenzyl)piperazine was reacted with triphosgene and N,N-dimethyl-1H-pyrazole-4-sulfonamide following conditions used in Steps D and E in the synthesis of compound **4** to provide 1-(4-(4-chlorobenzyl)piperazine-1-carbonyl)-N,N-dimethyl-1H-pyrazole-4-sulfonamide (**1**) as a white solid**.**

$^1$H NMR (300 MHz, $CDCl_3$) δ 8.46 (s, 1H), 7.82 (s, 1H), 7.26 - 7.33 (m, 4H), 3.86 (br, 4H), 3.53 (s, 2 H), 2.74 (s, 6H), 2.56 (s, 4H).

$^{13}$C NMR (101 MHz, DMSO-$d_6$) δ 149.18, 139.95, 136.69, 134.13, 131.62, 130.72, 128.20, 117.86, 60.69, 52.02, 37.45.

HRMS calculated for $C_{17}H_{22}ClN_5O_3S$ $[M+H]^+$ 412.1205, found 412.1208.

**N,N-dimethyl-1-(4-(2-morpholino-4-(trifluoromethyl)benzyl)piperazine-1-carbonyl)-1H-pyrazole-3-sulfonamide (3)**

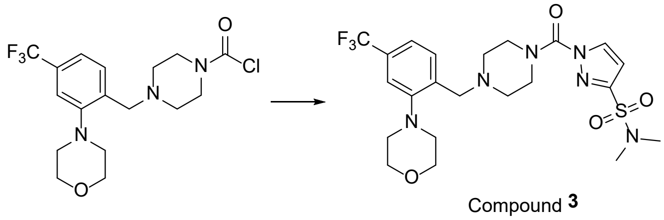


Compound 3

A 50-mL round-bottom flask was charged with N,N-dimethyl-1H-pyrazole-3-sulfonamide (107 mg, 0.610 mmol, 1.10 equiv), tetrahydrofuran (10 mL), 4-dimethylaminopyridine (20.4 mg, 0.170 mmol, 0.30 equiv), N,N-diisopropylethylamine (144 mg, 1.12 mmol, 2.00 equiv), 4-[[2-(morpholin-4-yl)-4-(trifluoromethyl)phenyl]methyl]piperazine-1-carbonyl chloride (218 mg, 0.560 mmol, 1.00 equiv). The resulting solution was stirred overnight at 60 °C and quenched with water (10 mL). The resulting mixture was extracted with DCM (3 x 15 mL) and the organic layers were combined, washed with brine (1 x 50 mL), dried over anhydrous $Na_2SO_4$, filtered and concentrated under reduced pressure. The crude product (450 mg) was purified by preparative HPLC using the following gradient conditions: 20% $CH_3CN$/80% Phase A increasing to 80% $CH_3CN$ over 10 min, then to 100% $CH_3CN$ over 0.1 min, holding at 100% $CH_3CN$ for 1.9 min, then reducing to 20% $CH_3CN$ over 0.1 min, and holding at 20% for 1.9 min, on a Waters 2767-5 Chromatograph. Column: Xbridge Prep C18, 19*150mm 5um; Mobile phase: Phase A: aqueous $NH_4HCO_3$ (0.05%); Phase B: $CH_3CN$; Detector, UV220 & 254nm. Purification resulted in 75.7 mg (26% yield) of N,N-dimethyl-1-[(4-[[2-(morpholin-4-yl)-4-(trifluoromethyl)phenyl]methyl]piperazin-1-yl)carbonyl]-1H-pyrazole-3-sulfonamide as a yellow solid.

$^1$H NMR (300MHz, $CDCl_3$) δ 8.18 (d, $J$ = 2.7 Hz, 1H), 7.62 - 7.60 (m, 1H), 7.36 - 7.32 (m, 2H), 6.74 (d, $J$ = 2.7 Hz, 1H), 3.87 - 3.84 (m, 8H), 3.65 (br, 2H), 2.98 - 2.96 (m, 4H), 2.86 (s, 6H), 2.60 (br, 4H).

$^{13}$C NMR (101 MHz, DMSO-$d_6$) δ 152.35, 149.37, 148.79, 136.89, 134.18, 131.07, 128.49 (q, $J$ = 31.4 Hz), 124.23 (q, $J$ = 272.3 Hz), 119.71 (q, $J$ = 3.7 Hz), 116.08 (d, $J$ = 3.4 Hz), 108.12, 66.50, 56.20, 52.45, 52.29, 37.61.
HRMS calculated for $C_{22}H_{29}F_3N_6O_4S$ $[M+H]^+$ 531.1996, found 531.1986.

**(S)-N,N-dimethyl-1-(2-methyl-4-(2-morpholino-4-(trifluoromethyl)benzyl)piperazine-1-carbonyl)-1H-pyrazole-3-sulfonamide (4)**

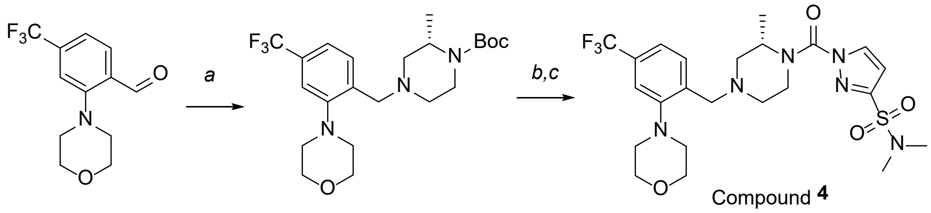


Compound 4

*Step A: tert-butyl (S)-2-methyl-4-(2-morpholino-4-(trifluoromethyl)benzyl)piperazine-1-carboxylate*

A 50-mL round-bottom flask was charged with 2-(morpholin-4-yl)-4-(trifluoromethyl)benzaldehyde (300 mg, 1.16 mmol, 1.00 equiv), tert-butyl (2S)-2-methylpiperazine-1-carboxylate (278 mg, 1.39 mmol, 1.20 equiv), DCM (10 mL). The mixture was stirred for 2 h at room temperature. Sodium triacetoxyborohydride (982 mg, 4.63 mmol, 4.00 equiv) was added. The resulting solution was stirred overnight at room temperature and then quenched with water (50 mL). The resulting mixture was extracted with DCM (3 x 50 mL) and the organic layers were combined, washed with brine (1 x 100 mL), dried over anhydrous $Na_2SO_4$, filtered and concentrated under reduced pressure. The residue was chromatographed on a silica gel column with MeOH/DCM (1/20) to provide 470 mg (92% yield) of tert-butyl (2S)-2-methyl-4-[[2-(morpholin-4-yl)-4-(trifluoromethyl)phenyl]methyl]piperazine-1-carboxylate as yellow oil. LCMS (ESI, *m/z*): 444 $[M+H]^+$.

*Step B: (S)-4-(2-((3-methylpiperazin-1-yl)methyl)-5-(trifluoromethyl)phenyl)morpholine*

A 250-mL round-bottom flask was charged with tert-butyl (2S)-2-methyl-4-[[2-(morpholin-4-yl)-4-(trifluoromethyl)phenyl]methyl]piperazine-1-carboxylate (470 mg, 1.06 mmol, 1.00 equiv), trifluoroacetic acid (5 mL), DCM (15 mL). The resulting solution was stirred overnight at room temperature and concentrated under reduced pressure. The crude product was dissolved in 1M NaOH solution (10 mL) and extracted with DCM (3 x 20 mL). The organic layers were combined, washed with brine (10 mL), dried over anhydrous $Na_2SO_4$, filtered and concentrated under reduced pressure to provide 340 mg (93% yield) of 4-(2-[[(3S)-3-methylpiperazin-1-yl]methyl]-5-(trifluoromethyl)phenyl)morpholine as brown oil and was carried to the next step without further purification. LCMS (ESI, *m/z*): 344 $[M+H]^+$.

*Step C: (S)-N,N-dimethyl-1-(2-methyl-4-(2-morpholino-4-(trifluoromethyl)benzyl)piperazine-1-carbonyl)-1H-pyrazole-3-sulfonamide (4)*

A 50-mL round-bottom flask was charged with triphosgene (103 mg, 0.350 mmol, 0.70 equiv), DCM (5 mL). N,N-Dimethyl-1H-pyrazole-3-sulfonamide (175 mg, 1.00 mmol, 2.00 equiv) was added at 0 °C. N,N-Diisopropylethylamine (256 mg, 1.98 mmol, 4.00 equiv) was added at 0 °C. The mixture was stirred for 2 h at room temperature. 4-(2-[[(3S)-3-methylpiperazin-1-yl]methyl]-5-(trifluoromethyl)phenyl)morpholine (170 mg, 0.500 mmol, 1.00 equiv) was added. The resulting solution was stirred overnight at room temperature and then quenched with water (50 mL). The resulting mixture was extracted with DCM (3 x 50 mL) and the organic layers were combined, washed with brine (1 x 100 mL), dried over anhydrous $Na_2SO_4$, filtered and concentrated under reduced pressure. The crude product (400 mg) was purified by preparative HPLC using the following gradient conditions: 20% $CH_3CN$/80% Phase A increasing to 80% $CH_3CN$ over 10 min, then to 100% $CH_3CN$ over 0.1 min, holding at 100% $CH_3CN$ for 1.9 min, then reducing to 20% $CH_3CN$ over 0.1 min, and holding at 20% for 1.9 min, on a Waters 2767-5 Chromatograph. Column: Xbridge Prep C18, 19*150mm 5um; Mobile phase: Phase A: aqueous $NH_4HCO_3$ (0.05%); Phase B: $CH_3CN$; Detector, UV220 & 254nm. Purification resulted in 98.7 mg (37% yield) of N,N-dimethyl-1-[[(2S)-2-methyl-4-[[2-(morpholin-4-yl)-4-(trifluoromethyl)phenyl]methyl]piperazin-1-yl]carbonyl]-1H-pyrazole-3-sulfonamide as a white solid.

$^1$H NMR (400 MHz, $CDCl_3$) δ 8.17 (d, *J* = 2.7 Hz, 1H), 7.61 (br, 1H), 7.39 – 7.28 (m, 2H), 6.74 (d, *J* = 2.7 Hz, 1H), 4.67 (br, 1H), 4.38 – 4.28 (m, 1H), 3.93 – 3.77 (m, 4H), 3.73 – 3.48 (m, 2H), 3.47 – 3.39 (m, 1H), 3.08 – 2.90 (m, 5H), 2.85 (s, 6H), 2.76 – 2.63 (m, 1H), 2.48 – 2.18 (m, 2H), 1.46 (d, *J* = 6.2 Hz, 3H).
$^{13}$C NMR (101 MHz, DMSO-$d_6$) δ 152.41, 149.37, 148.74, 136.73, 134.18, 131.19, 128.54 (q, *J* = 31.3 Hz), 124.23 (q, *J* = 272.3 Hz), 119.55 (q, *J* = 3.9 Hz), 115.96 (q, *J* = 3.9 Hz), 108.14, 66.51, 57.29, 56.41, 52.45, 52.40, 37.60, 16.09.
HRMS calculated for $C_{23}H_{31}F_3N_6O_4S$ $[M+H]^+$ 545.2152, found 545.2158.

**2-((1R,5S)-8-(2-(((S)-4-(3-(N,N-dimethylsulfamoyl)-1H-pyrazole-1-carbonyl)-3-methylpiperazin-1-yl)methyl)-5-(trifluoromethyl)phenyl)-3,8-diazabicyclo[3.2.1]octan-3-yl)acetic acid (5, ABD957*)***

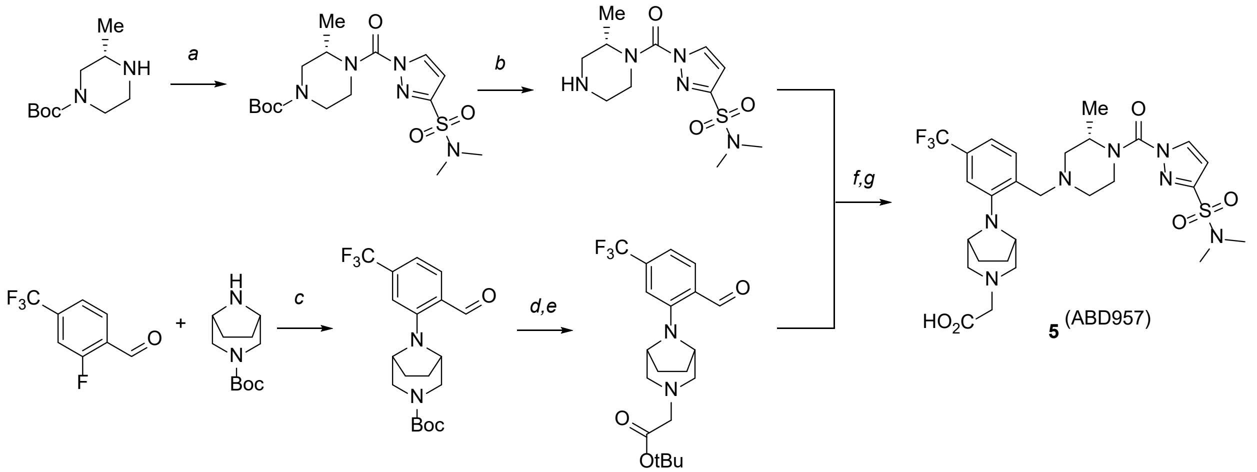


*Step A: tert-butyl (S)-4-(3-(N,N-dimethylsulfamoyl)-1H-pyrazole-1-carbonyl)-3-methylpiperazine-1-carboxylate*

A 40-mL vial was charged with tert-butyl (3S)-3-methylpiperazine-1-carboxylate (600 mg, 3.00 mmol, 1.00 equiv), DCM (10 mL), triphosgene (446 mg, 1.50 mmol, 0.50 equiv). N,N-Diisopropylethylamine (1.16 g, 8.99 mmol, 3.00 equiv) was added dropwise at 0 °C. The resulting solution was stirred for 2 h at room temperature and quenched by water (10 mL). The mixture was extracted with DCM (3 x 10 mL) and the organic layers were combined, washed with brine (3 x 10 mL), dried over anhydrous $Na_2SO_4$, filtered and concentrated under reduced pressure to provide 600 mg (crude) of tert-butyl (3S)-4-(carbonochloridoyl)-3-methylpiperazine-1-carboxylate as yellow oil which was carried to the next step without further purification.

A 40-mL vial was charged with tert-butyl (3S)-4-(carbonochloridoyl)-3-methylpiperazine-1-carboxylate (600 mg, 2.28 mmol, 1.00 equiv), tetrahydrofuran (10 mL), N,N-dimethyl-1H-pyrazole-3-sulfonamide (401 mg, 2.29 mmol, 1.00 equiv), N,N-diisopropylethylamine (886 mg, 6.86 mmol, 3.00 equiv) and 4-dimethylaminopyridine (56.0 mg, 0.460 mmol, 0.20 equiv). The resulting solution was stirred overnight at 60 °C and quenched by water (10 mL). The mixture was extracted with EtOAc (3 x 10 mL) and the organic layers were combined, washed with brine (3 x 10 mL), dried over anhydrous $Na_2SO_4$, filtered and concentrated under reduced pressure. The residue was chromatographed on a silica gel column with EtOAc/petroleum ether (1/1) to provide 900 mg (98% yield) of tert-butyl (3S)-4-[[3-(dimethylsulfamoyl)-1H-pyrazol-1-yl]carbonyl]-3-methylpiperazine-1-carboxylate as yellow oil. LCMS (ESI, *m/z*): 402 $[M+H]^+$.

*Step B: (S)-N,N-dimethyl-1-(2-methylpiperazine-1-carbonyl)-1H-pyrazole-3-sulfonamide*

A 40-mL vial was charged with tert-butyl (3S)-4-[[3-(dimethylsulfamoyl)-1H-pyrazol-1-yl]carbonyl]-3-methylpiperazine-1-carboxylate (900 mg, 2.24 mmol, 1.00 equiv), DCM (10 mL), trifluoroacetic acid (4 mL). The resulting solution was stirred for 2 h at room temperature and concentrated under reduced pressure to provide 660 mg (crude) of N,N-dimethyl-1-[[(2S)-2-methylpiperazin-1-yl]carbonyl]-1H-pyrazole-3-sulfonamide as yellow oil which was carried to the next step without further purification. LCMS (ESI, *m/z*): 302 $[M+H]^+$.

*Step C: tert-butyl 8-(2-formyl-5-(trifluoromethyl)phenyl)-3,8-diazabicyclo[3.2.1]octane-3-carboxylate*

A 40-mL vial was charged with dimethyl sulphoxide (10 mL), 2-fluoro-4-(trifluoromethyl)benzaldehyde (576 mg, 3.00 mmol, 1.00 equiv), tert-butyl 3,8-diazabicyclo[3.2.1]octane-3-carboxylate (623 mg, 2.93 mmol, 1.00 equiv), potassium carbonate (1.24 g, 8.97 mmol, 3.00 equiv) under nitrogen. The resulting solution was stirred overnight at 120 °C and quenched by water (10 ml). The mixture was extracted with EtOAc (3 x 10 mL) and the organic layers were combined, washed with brine (1 x 10 mL), dried over anhydrous $Na_2SO_4$, filtered and concentrated under reduced pressure. The residue was chromatographed on a silica gel column with EtOAc/petroleum ether (1/3) to provide 450 mg (39% yield) of tert-butyl 8-[2-formyl-5-(trifluoromethyl)phenyl]-3,8-diazabicyclo[3.2.1]octane-3-carboxylate as yellow oil. LCMS (ESI, *m/z*): 385 $[M+H]^+$.

*Step D: 2-[3,8-diazabicyclo[3.2.1]octan-8-yl]-4-(trifluoromethyl)benzaldehyde*

A 40-mL vial was charged with tert-butyl 8-[2-formyl-5-(trifluoromethyl)phenyl]-3,8-diazabicyclo[3.2.1]octane-3-carboxylate (150 mg, 0.390 mmol, 1.00 equiv), DCM (10 mL), trifluoroacetic acid (2 mL). The resulting solution was stirred for 2 h at room temperature and concentrated under reduced pressure. The pH value of the solution was adjusted to 9 with sodium hydroxide solution (1 mol/L aq). The mixture was extracted with DCM (3 x 30 mL) and the organic layers were combined, washed with brine (2 x 20 mL), dried over anhydrous $Na_2SO_4$, filtered and concentrated under reduced pressure to yield 100 mg (crude) of 2-[3,8-diazabicyclo[3.2.1]octan-8-yl]-4-(trifluoromethyl)benzaldehyde as yellow oil. LCMS (ESI, *m/z*): 285 $[M+H]^+$.

*Step E: tert-butyl 2-[8-[2-formyl-5-(trifluoromethyl)phenyl]-3,8-diazabicyclo[3.2.1]octan-3-yl]acetate*

A 40-mL vial was charged with 2-[3,8-diazabicyclo[3.2.1]octan-8-yl]-4-(trifluoromethyl)benzaldehyde (100 mg, 0.350 mmol, 1.00 equiv), acetone (10 mL), tert-butyl 2-bromoacetate (137 mg, 0.700 mmol, 2.00 equiv), potassium carbonate (146 mg, 1.06 mmol, 3.00 equiv). The resulting solution was stirred overnight at 80 °C and quenched with water (10 ml). The mixture was extracted with EtOAc (3 x 10 mL) and the organic layers were combined, washed with brine (1 x 10 mL), dried over anhydrous $Na_2SO_4$, filtered and concentrated under reduced pressure. The residue was chromatographed on a silica gel column with EtOAc/petroleum ether (1/3) to provide 125 mg (86% yield) of tert-butyl 2-[8-[2-formyl-5-(trifluoromethyl)phenyl]-3,8-diazabicyclo[3.2.1]octan-3-yl]acetate as yellow oil. LCMS (ESI, *m/z*): 399 $[M+H]^+$.

*Step F: tert-butyl 2-((1R,5S)-8-(2-(((S)-4-(3-(N,N-dimethylsulfamoyl)-1H-pyrazole-1-carbonyl)-3-methylpiperazin-1-yl)methyl)-5-(trifluoromethyl)phenyl)-3,8-diazabicyclo[3.2.1]octan-3-yl)acetate*

A 40-mL vial was charged with 1,2-dichloroethane (10 mL), N,N-dimethyl-1-[[(2S)-2-methylpiperazin-1-yl]carbonyl]-1H-pyrazole-3-sulfonamide (125 mg, 0.410 mmol, 1.00 equiv), tert-butyl 2-8-[2-formyl-5-(trifluoromethyl)phenyl]-3,8-diazabicyclo[3.2.1]octan-3-ylacetate (165 mg, 0.410 mmol, 1.00 equiv), triethylamine (126 mg, 1.25 mmol, 3.00 equiv). The resulting solution was stirred for 1 h at room temperature, then sodium triacetoxyborohydride (264 mg, 1.25 mmol, 3.00 equiv) was added. The resulting solution was stirred overnight at room temperature and then quenched with water (10 mL). The resulting mixture was extracted with DCM (3 x 10 mL) and the organic layers were combined, washed with brine (1 x 10 mL), dried over anhydrous $Na_2SO_4$, filtered and concentrated under reduced pressure. The residue was chromatographed on a silica gel column with EtOAc/petroleum ether (1/1) to provide 100 mg (35% yield) of tert-butyl 2-[8-(2-[[(3S)-4-[[3-(dimethylsulfamoyl)-1H-pyrazol-1-yl]carbonyl]-3-methylpiperazin-1-yl]methyl]-5-(trifluoromethyl)phenyl)-3,8-diazabicyclo[3.2.1]octan-3-yl]acetate as yellow oil. LCMS (ESI, *m/z*): 684 $[M+H]^+$.

*Step G: 2-((1R,5S)-8-(2-(((S)-4-(3-(N,N-dimethylsulfamoyl)-1H-pyrazole-1-carbonyl)-3-methylpiperazin-1-yl)methyl)-5-(trifluoromethyl)phenyl)-3,8-diazabicyclo[3.2.1]octan-3-yl)acetic acid (**5**, ABD957)*

A 40-mL vial was charged with tert-butyl 2-[8-(2-[[(3S)-4-[[3-(dimethylsulfamoyl)-1H-pyrazol-1-yl]carbonyl]-3-methylpiperazin-1-yl]methyl]-5-(trifluoromethyl)phenyl)-3,8-diazabicyclo[3.2.1]octan-3-yl]acetate (100 mg, 0.150 mmol, 1.00 equiv), DCM (10 mL), trifluoroacetic acid (4 mL). The resulting solution was stirred for 2 h at room temperature and concentrated under reduced pressure. The crude product (100 mg) was purified by preparative HPLC using the following gradient conditions: 38% $CH_3CN$/62% Phase A increasing to 60% $CH_3CN$ over 7 min, then to 100% $CH_3CN$ over 0.1 min, holding at 100% $CH_3CN$ for 1.9 min, then reducing to 38% $CH_3CN$

over 0.1 min, and holding at 38% for 1.9 min, on a Waters 2767-5 Chromatograph. Column: Xbridge Prep $C_{18}$, 19*150mm 5um; Mobile phase: Phase A: aqueous $NH_4HCO_3$ (0.05%); Phase B: $CH_3CN$; Detector, UV220 & 254nm. Purification resulted in 20.4 mg (22% yield) of 2-[8-(2-[[(3S)-4-[[3-(dimethylsulfamoyl)-1H-pyrazol-1-yl]carbonyl]-3-methylpiperazin-1-yl]methyl]-5-(trifluoromethyl)phenyl)-3,8-diazabicyclo[3.2.1]octan-3-yl]acetic acid as a white solid.

$^1$H NMR (300 MHz, MeOD) δ 8.29 (d, $J$ = 2.7 Hz, 1H), 7.66 (d, $J$ = 8.0 Hz, 1H), 7.31 (d, $J$ = 7.9 Hz, 1H), 7.18 (s, 1H), 6.82 (d, $J$ = 2.7 Hz, 1H), 4.56 (br, 1H), 4.30 (d, $J$ = 17.2 Hz, 2H), 4.20 (d, $J$ = 13.8 Hz, 1H), 3.75 – 3.23 (m, 9H), 2.96 (d, $J$ = 10.4 Hz, 1H), 2.83 (s, 6H), 2.79 (s, 1H), 2.43 (dd, $J$ = 11.6, 3.7 Hz, 1H), 2.38 – 2.26 (m, 1H), 2.15 (s, 4H), 1.45 (d, $J$ = 6.8 Hz, 3H).
$^{13}$C NMR (101 MHz, DMSO-$d_6$) δ 171.56, 150.33, 149.38, 148.73, 134.18, 133.86, 131.64, 128.23 (q, $J$ = 30.7 Hz), 124.32 (q, $J$ = 272.1 Hz), 117.11 (q, $J$ = 3.4 Hz), 113.13 (q, $J$ = 3.3 Hz), 108.14, 59.53, 59.34, 58.32, 58.26, 57.75, 57.61, 57.33, 52.55, 37.62, 26.93, 16.09.
HRMS calculated for $C_{27}H_{36}F_3N_7O_5S$ $[M+H]^+$ 628.2523, found 628.2534.

Compound **1**

1H NMR (300 MHz, CDCl3) δ 8.46 (s, 1H), 7.82 (s, 1H), 7.26 - 7.33 (m, 4H), 3.86 (br, 4H), 3.53 (s, 2 H), 2.74 (s, 6H), 2.56 (s, 4H).

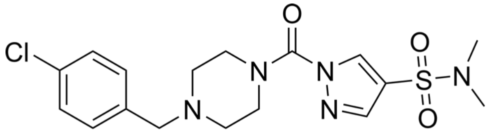


Compound **1**

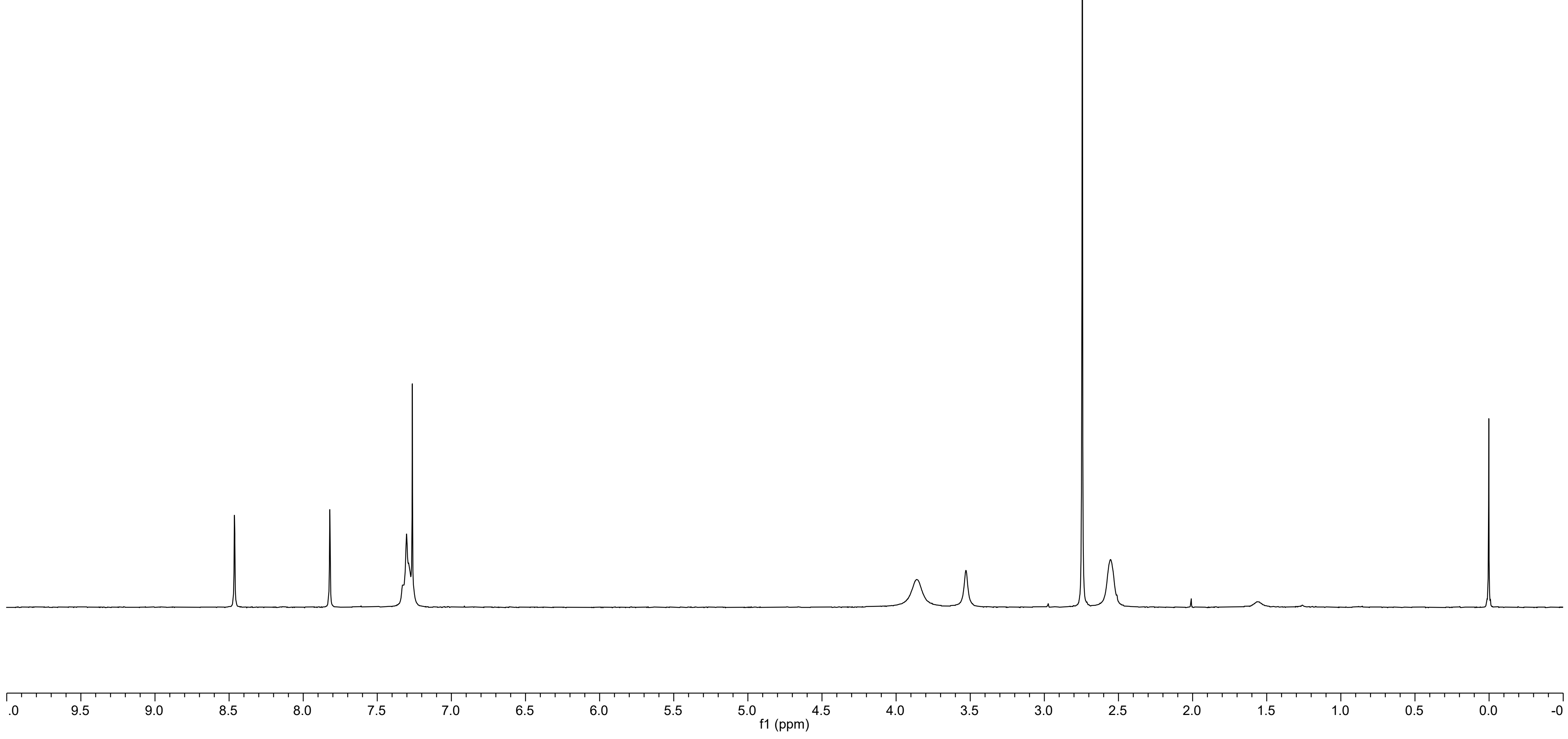

Compound **1**

13C NMR (101 MHz, DMSO-d6) δ 149.18, 139.95, 136.69, 134.13, 131.62, 130.72, 128.20, 117.86, 60.69, 52.02, 37.45.

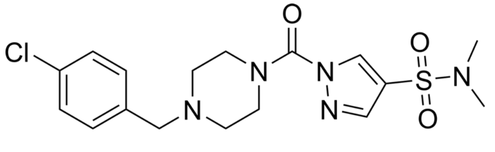


Compound **1**

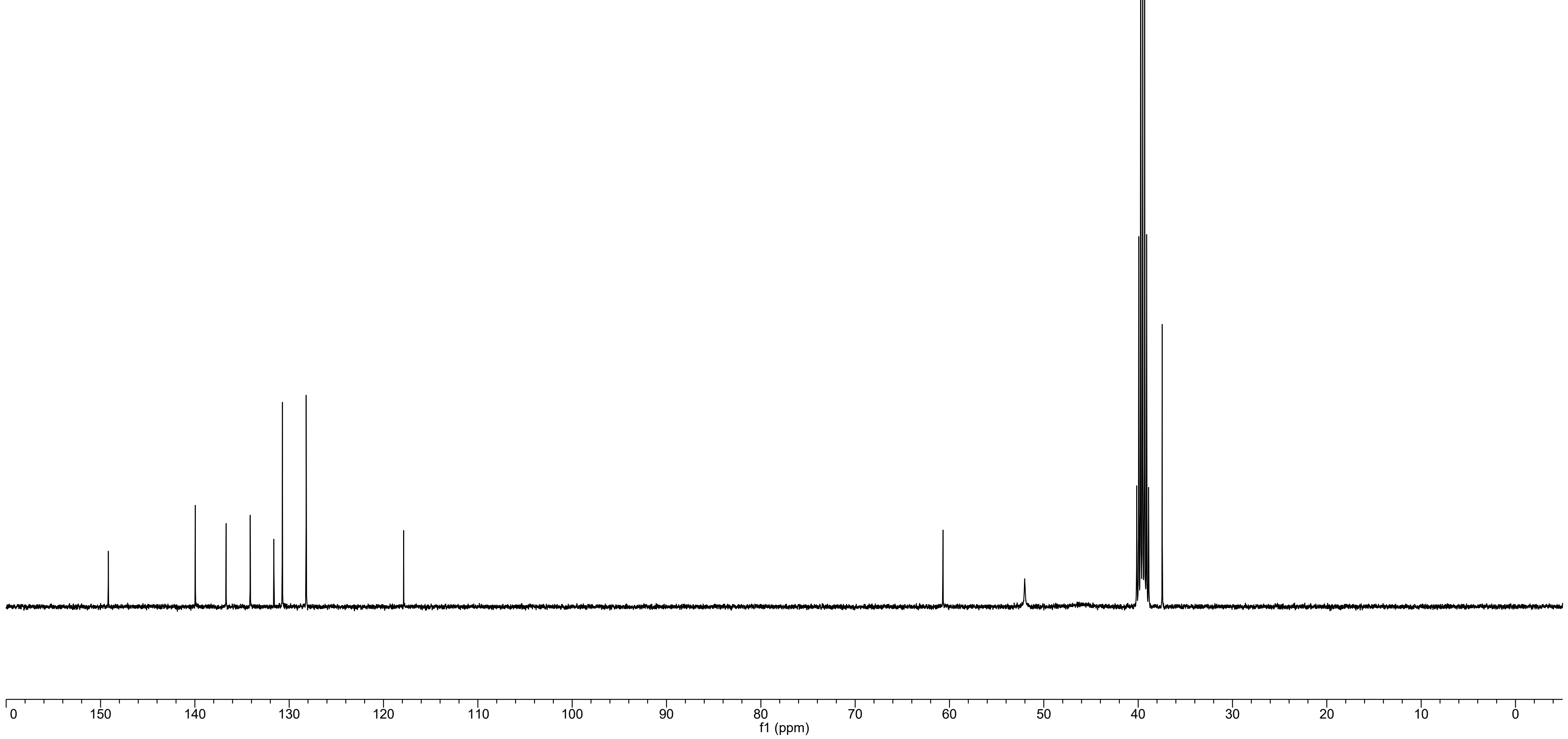

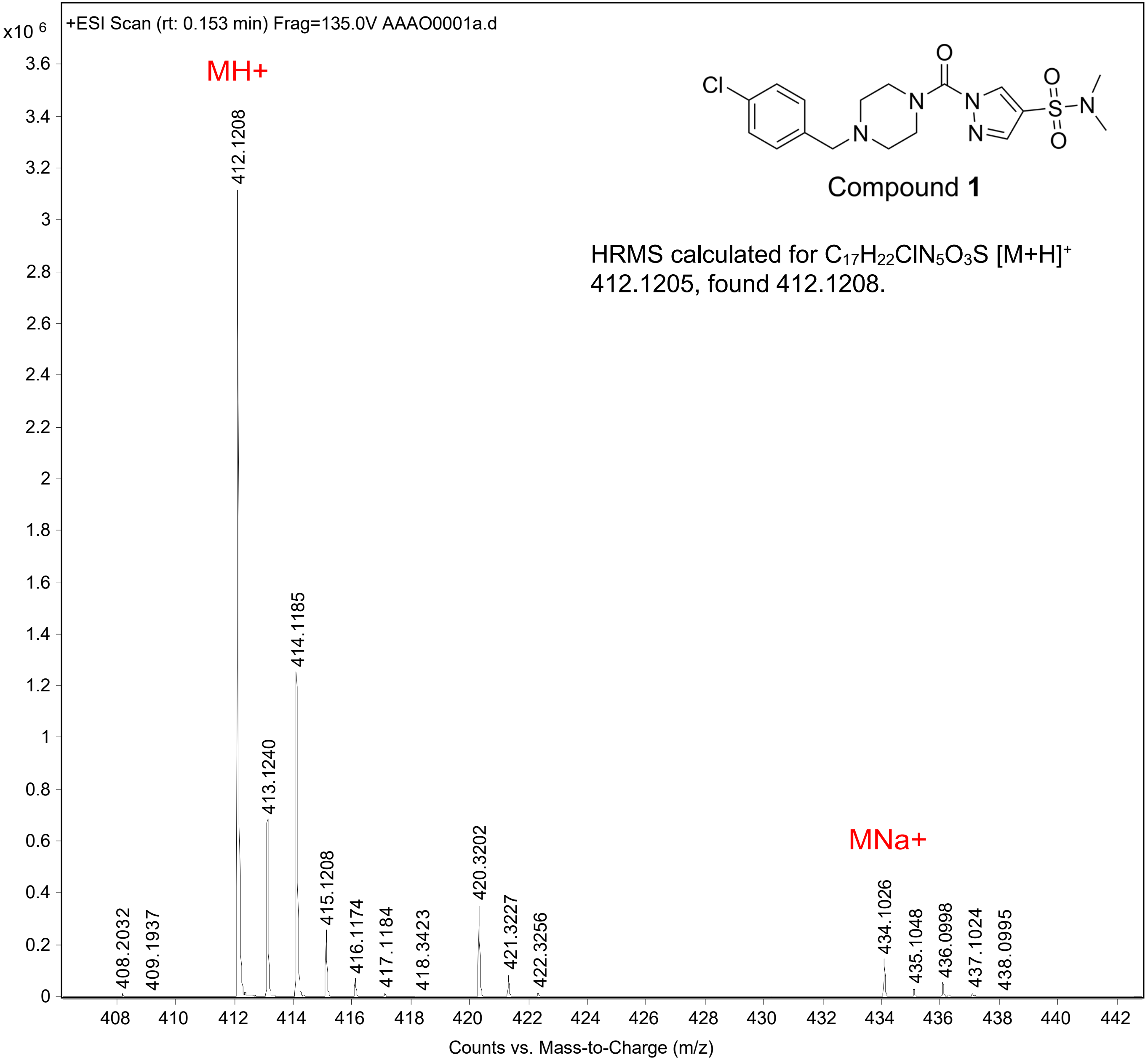
+ESI Scan (rt: 0.153 min) Frag=135.0V AAAO0001a.d
x10 6
MH+
MNa+
Compound 1
HRMS calculated for $C_{17}H_{22}ClN_5O_3S$ $[M+H]^+$ 412.1205, found 412.1208.
412.1208
413.1240
414.1185
415.1208
416.1174
417.1184
418.3423
420.3202
421.3227
422.3256
408.2032
409.1937
434.1026
435.1048
436.0998
437.1024
438.0995
Counts vs. Mass-to-Charge (m/z)

Compound **2**

1H NMR (300MHz, CDCl3) δ 8.47 (s, 1H), 7.82 (s, 1H), 7.61 - 7.66 (m, 1H), 7.31 - 7.40 (m, 2H), 3.77 - 3.92 (m, 8H), 3.67 (br, 2H), 2.87 - 3.02 (m, 4H), 2.74 (s, 6H), 2.63 (br, 4H).

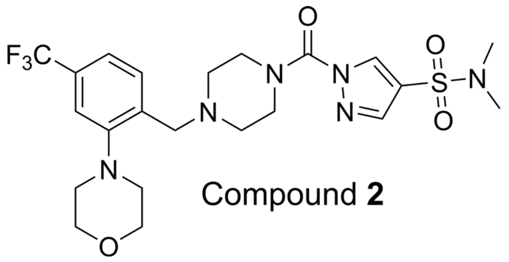


Compound **2**

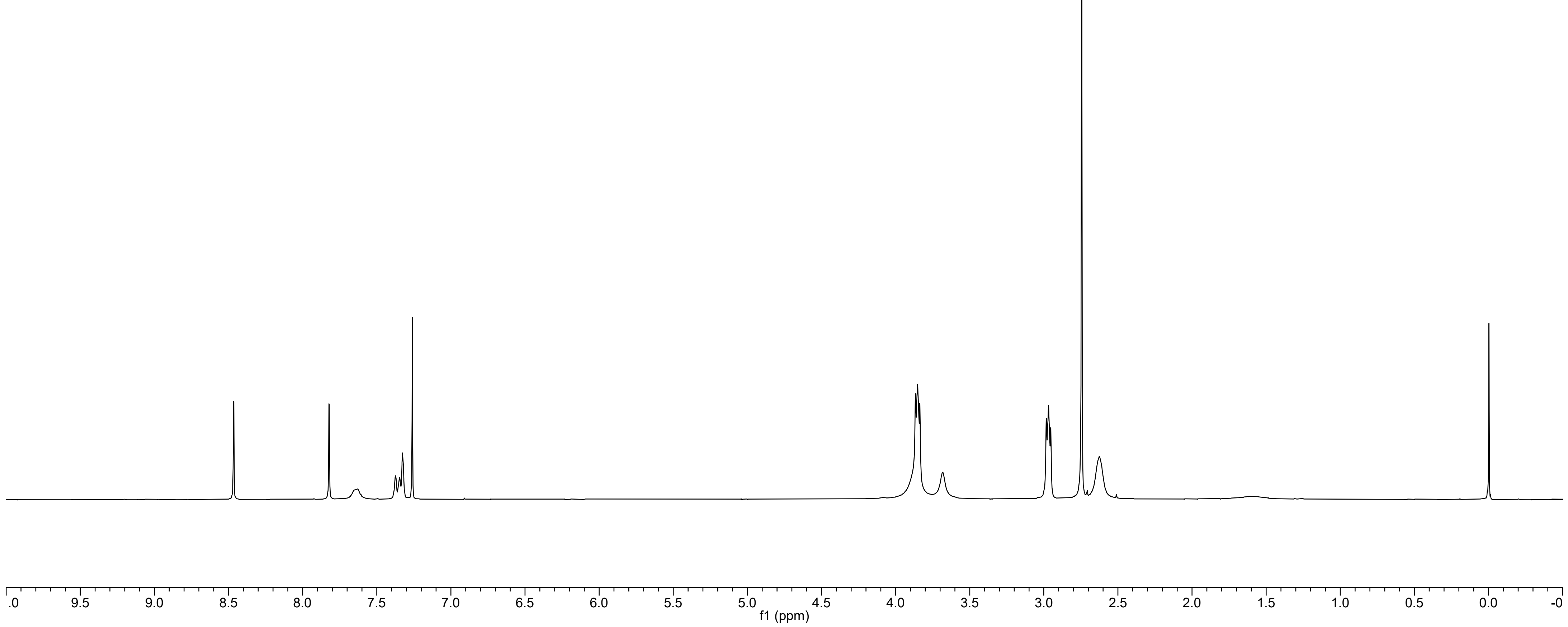

Compound **2**

13C NMR (101 MHz, DMSO-d6) δ 152.36, 149.20, 139.96, 136.85, 134.12, 131.10, 128.50 (q, J = 31.5 Hz), 124.23 (q, J = 272.3 Hz), 119.68 (q, J = 3.5 Hz), 117.88, 116.07 (q, J = 3.6 Hz), 66.50, 56.25, 52.46, 52.33, 37.46.

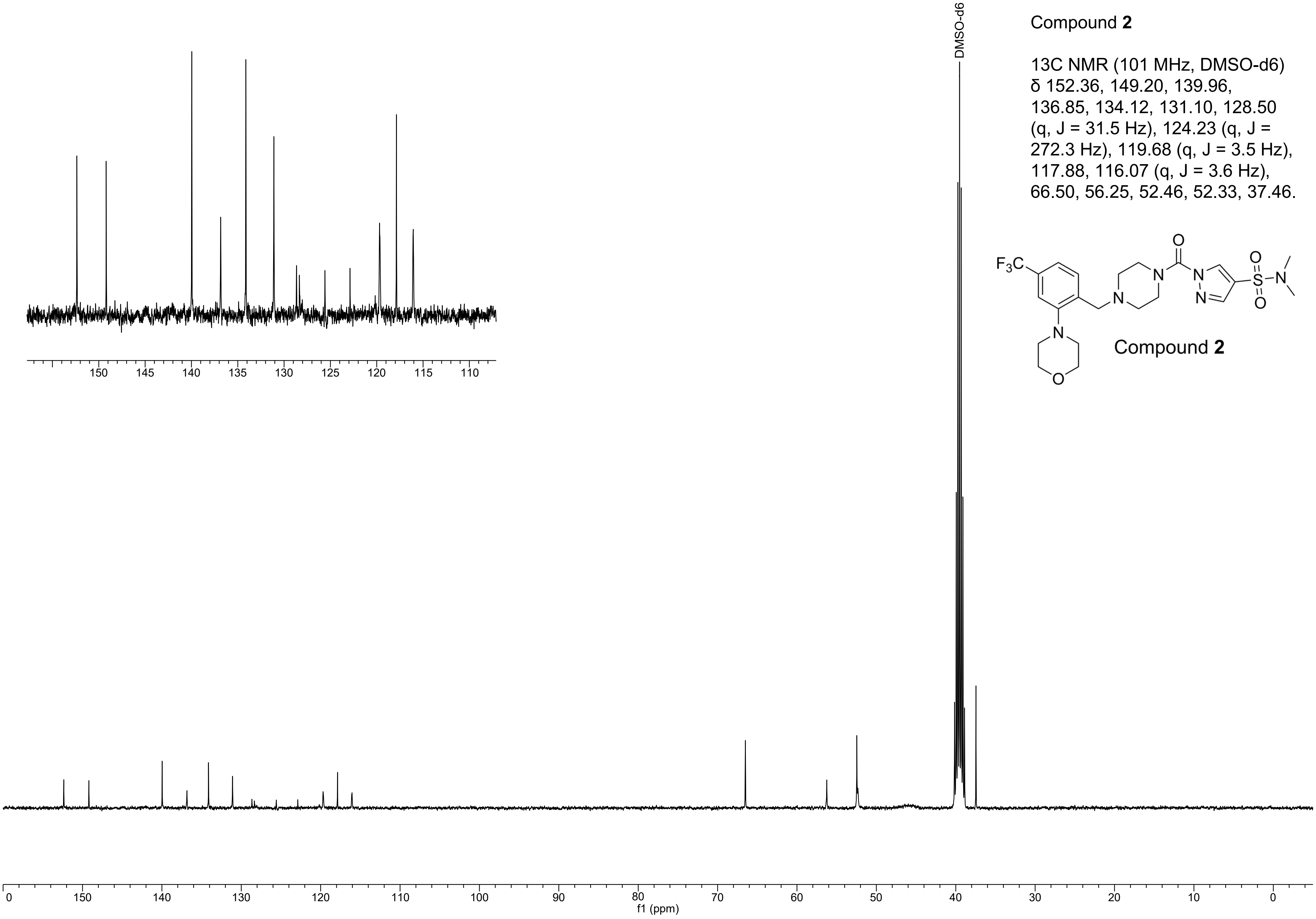


Compound **2**

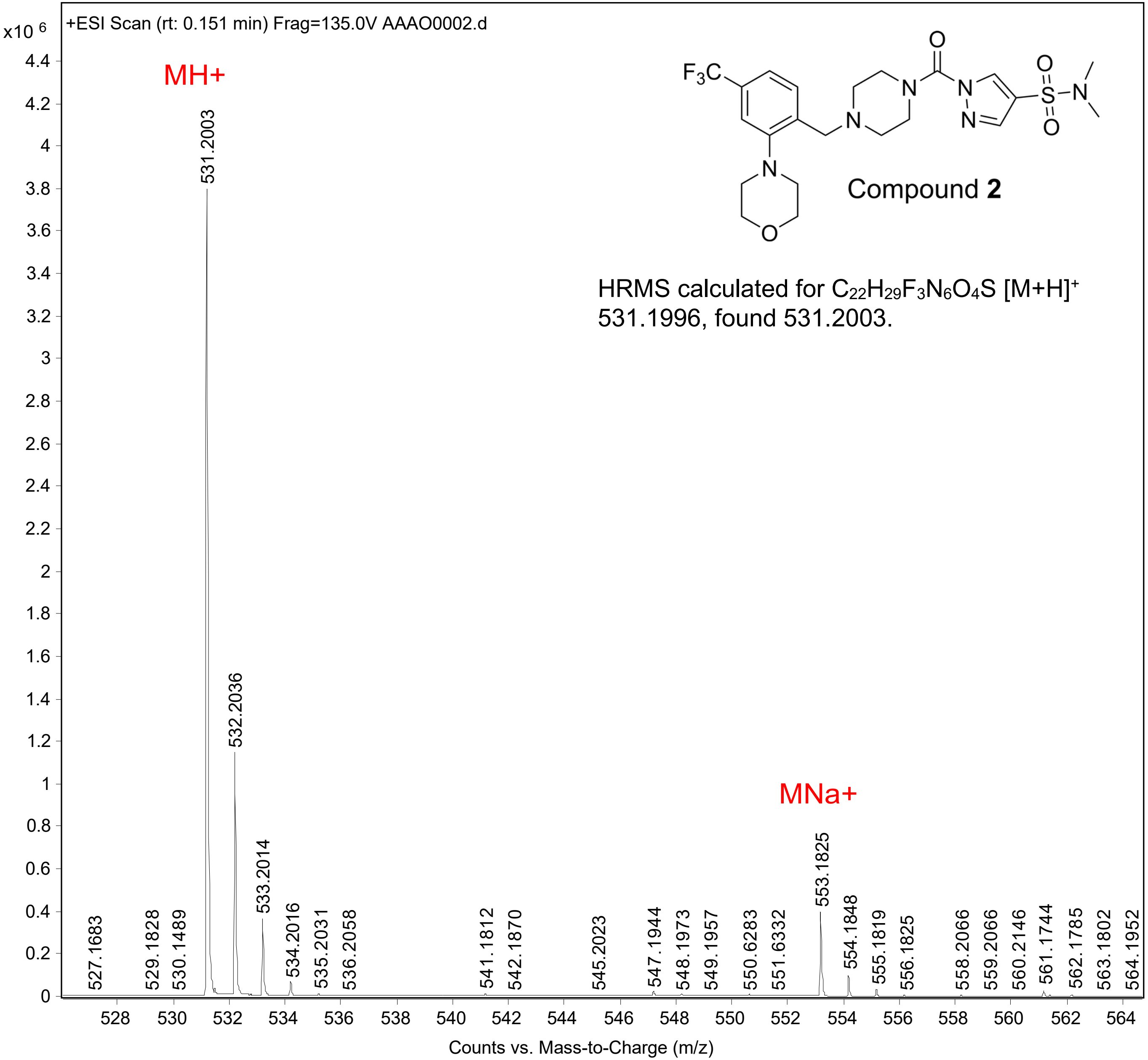

+ESI Scan (rt: 0.151 min) Frag=135.0V AAAO0002.d
x10 6
MH+
531.2003
532.2036
533.2014
534.2016
535.2031
536.2058
527.1683
529.1828
530.1489
541.1812
542.1870
545.2023
547.1944
548.1973
549.1957
550.6283
551.6332
MNa+
553.1825
554.1848
555.1819
556.1825
558.2066
559.2066
560.2146
561.1744
562.1785
563.1802
564.1952
F3C
Compound 2
HRMS calculated for C22H29F3N6O4S [M+H]+ 531.1996, found 531.2003.
Counts vs. Mass-to-Charge (m/z)

Compound **3**

1H NMR (300MHz, CDCl3) δ 8.18 (d, J = 2.7 Hz, 1H), 7.62 - 7.60 (m, 1H), 7.36 - 7.32 (m, 2H), 6.74 (d, J = 2.7 Hz, 1H), 3.87 - 3.84 (m, 8H), 3.65 (br, 2H), 2.98 - 2.96 (m, 4H), 2.86 (s, 6H), 2.60 (br, 4H).



Compound **3**

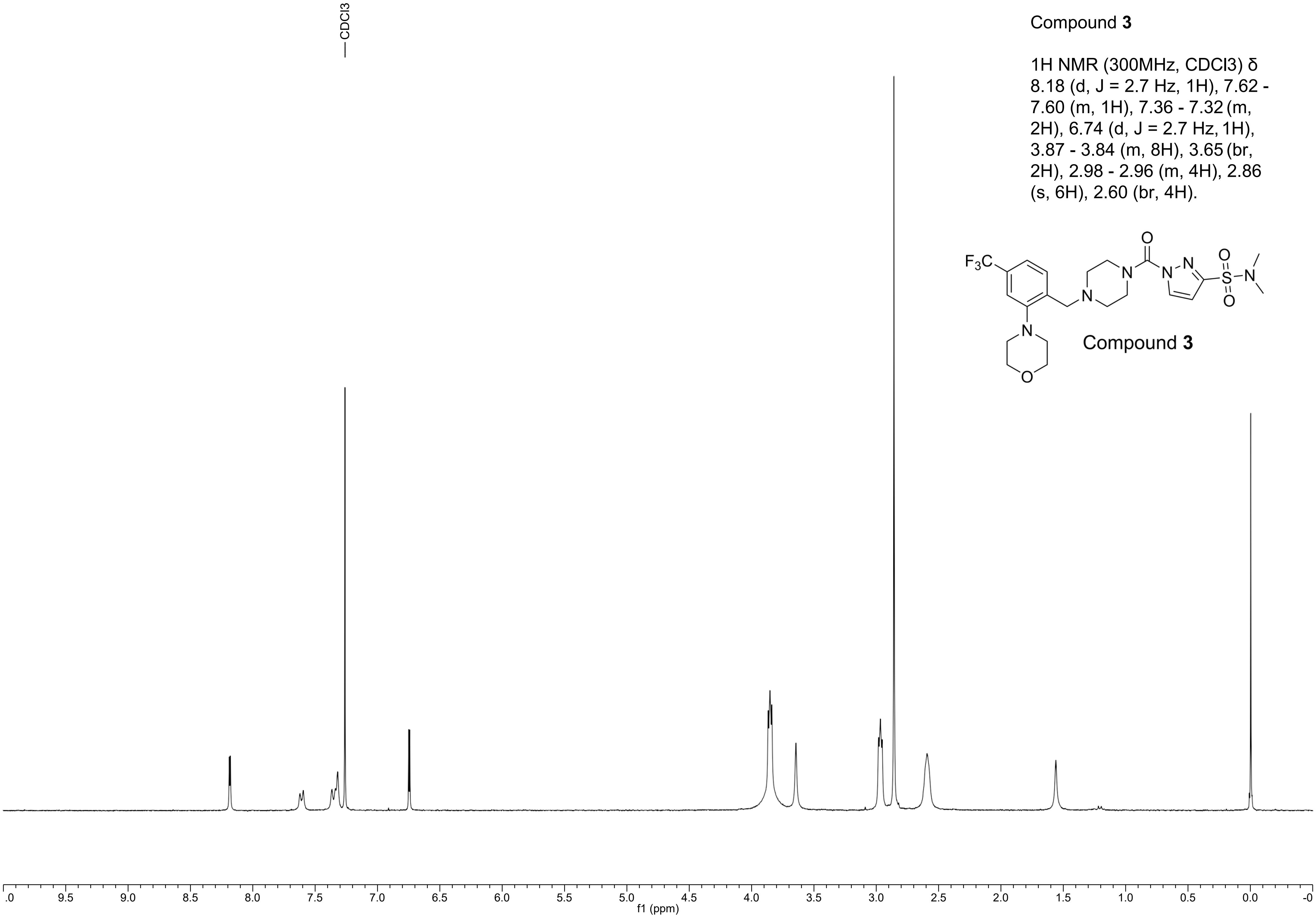

Compound **3**

13C NMR (101 MHz, DMSO-d6) δ 152.35, 149.37, 148.79, 136.89, 134.18, 131.07, 128.49 (q, J = 31.4 Hz), 124.23 (q, J = 272.3 Hz), 119.71 (q, J = 3.7 Hz), 116.08 (d, J = 3.4 Hz), 108.12, 66.50, 56.20, 52.45, 52.29, 37.61.

Compound **3**

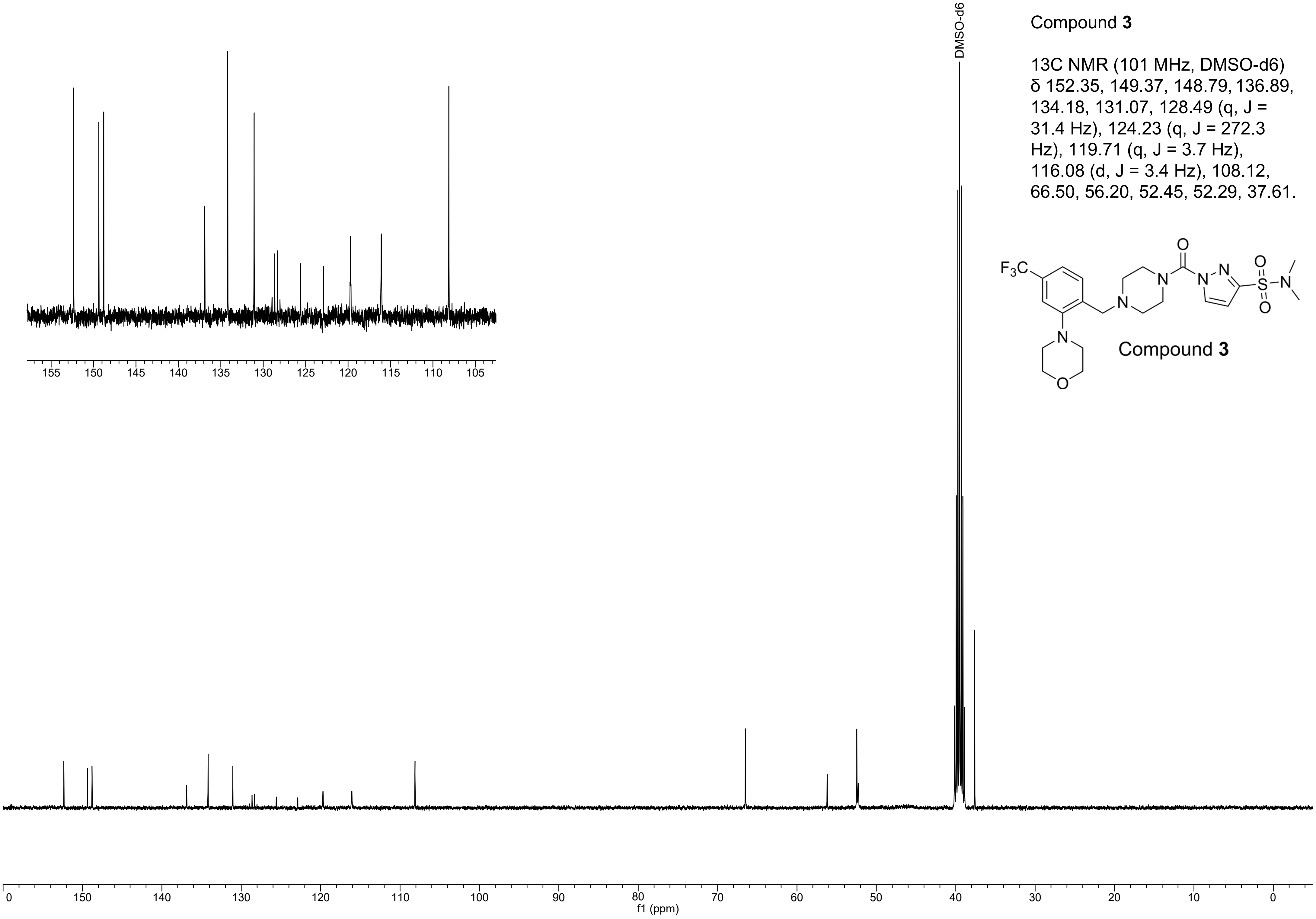

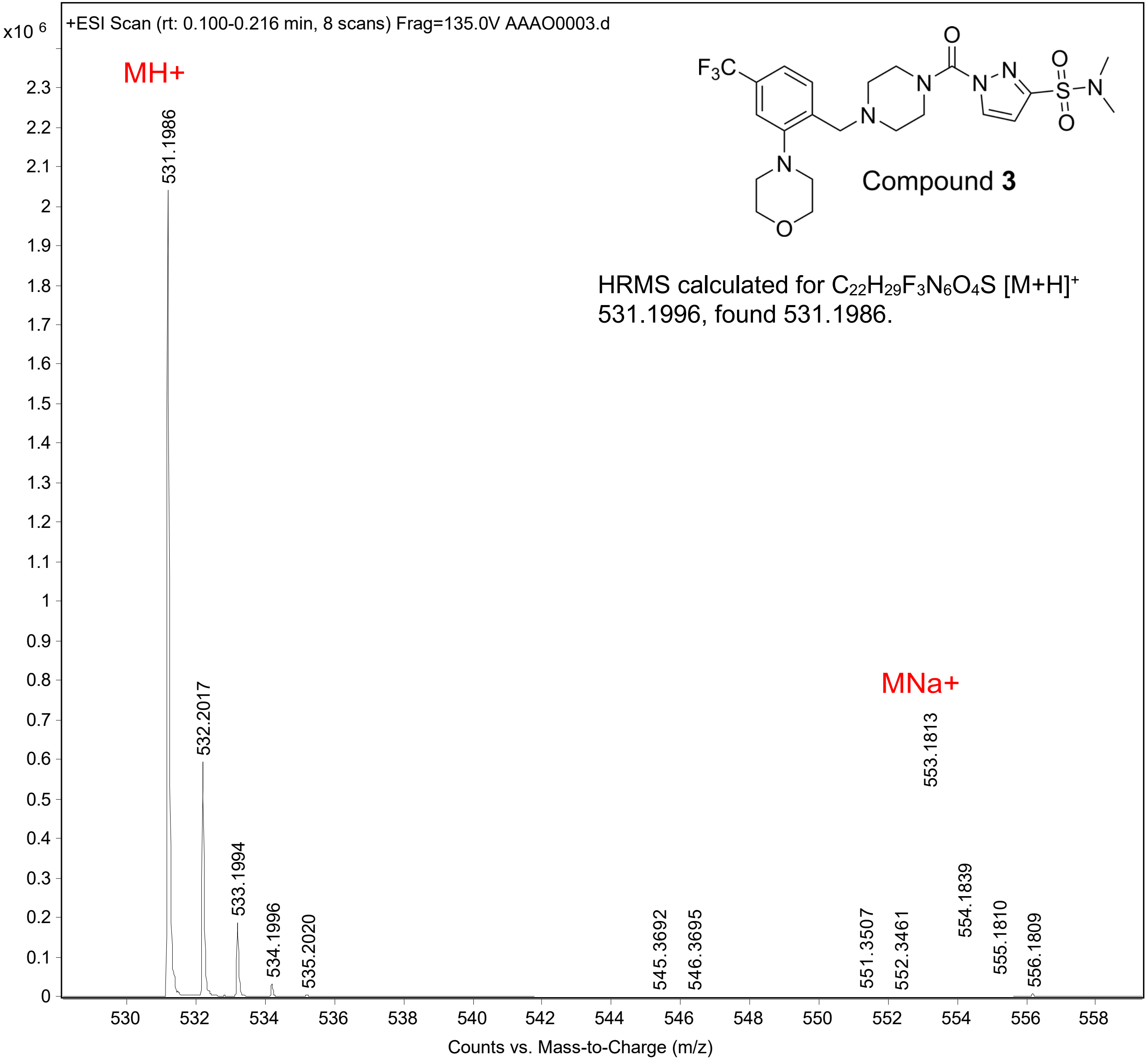

x10 6
+ESI Scan (rt: 0.100-0.216 min, 8 scans) Frag=135.0V AAAO0003.d
MH+
MNa+
531.1986
532.2017
533.1994
534.1996
535.2020
545.3692
546.3695
551.3507
552.3461
553.1813
554.1839
555.1810
556.1809
Compound 3
HRMS calculated for C22H29F3N6O4S [M+H]+ 531.1996, found 531.1986.
Counts vs. Mass-to-Charge (m/z)

Compound **4**

1H NMR (400 MHz, CDCl3) δ 8.17 (d, J = 2.7 Hz, 1H), 7.61 (br, 1H), 7.39 – 7.28 (m, 2H), 6.74 (d, J = 2.7 Hz, 1H), 4.67 (br, 1H), 4.38 – 4.28 (m, 1H), 3.93 – 3.77 (m, 4H), 3.73 – 3.48 (m, 2H), 3.47 – 3.39 (m, 1H), 3.08 – 2.90 (m, 5H), 2.85 (s, 6H), 2.76 – 2.63 (m, 1H), 2.48 – 2.18 (m, 2H), 1.46 (d, J = 6.2 Hz, 3H).

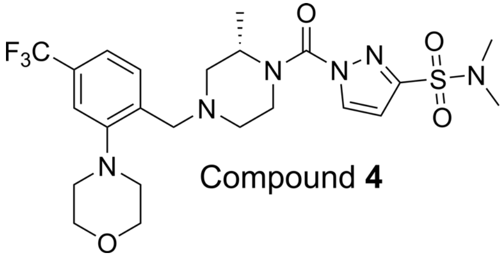


Compound **4**

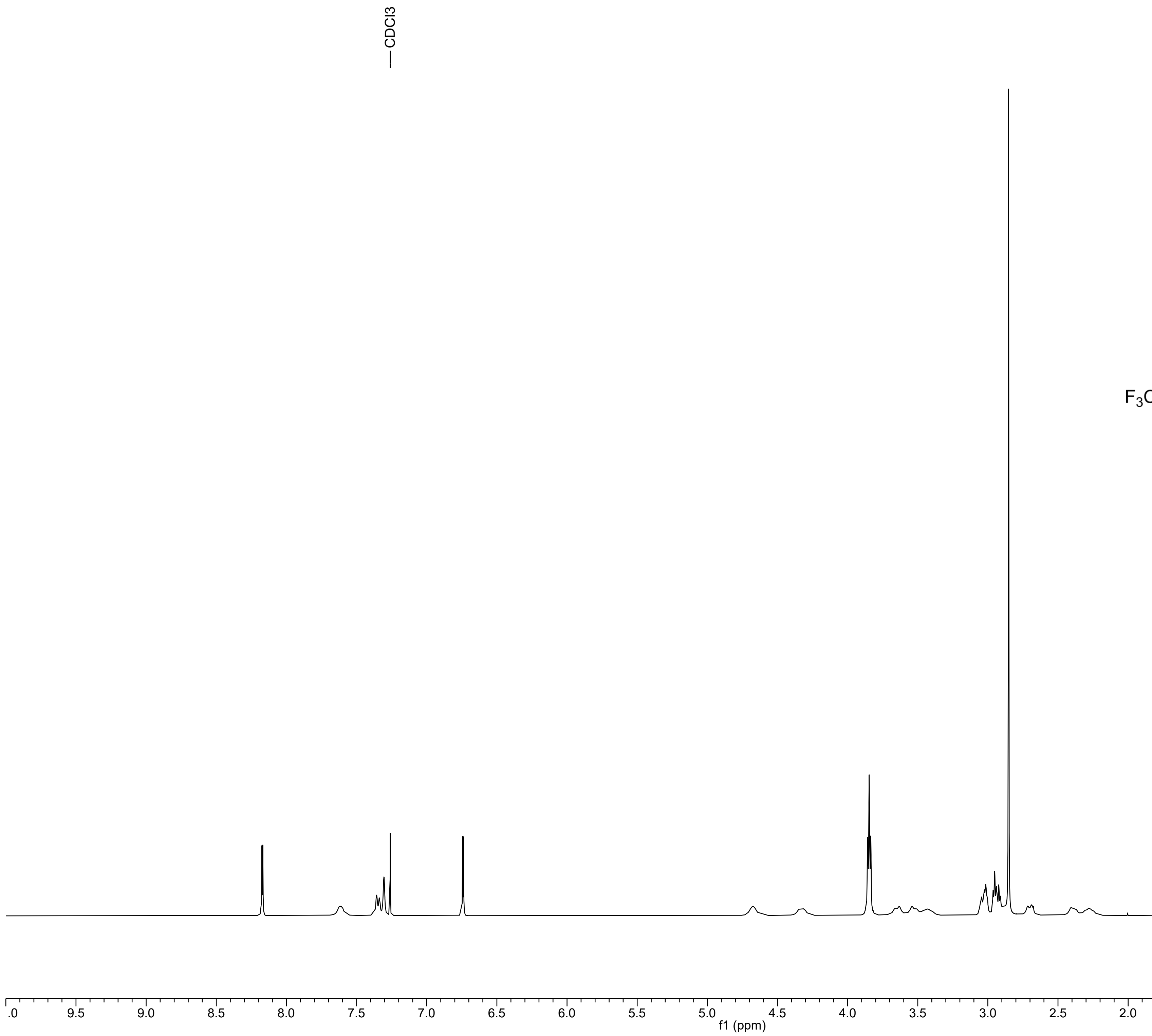

Compound **4**

13C NMR (101 MHz, DMSO-d6) δ 152.41, 149.37, 148.74, 136.73, 134.18, 131.19, 128.54 (q, J = 31.3 Hz), 124.23 (q, J = 272.3 Hz), 119.55 (q, J = 3.9 Hz), 115.96 (q, J = 3.9 Hz), 108.14, 66.51, 57.29, 56.41, 52.45, 52.40, 37.60, 16.09.

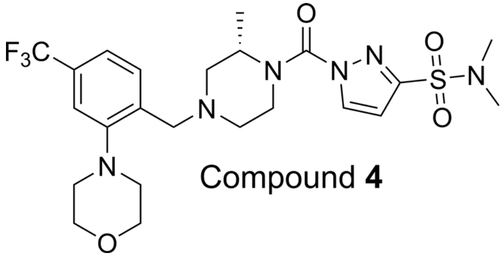

Compound **4**

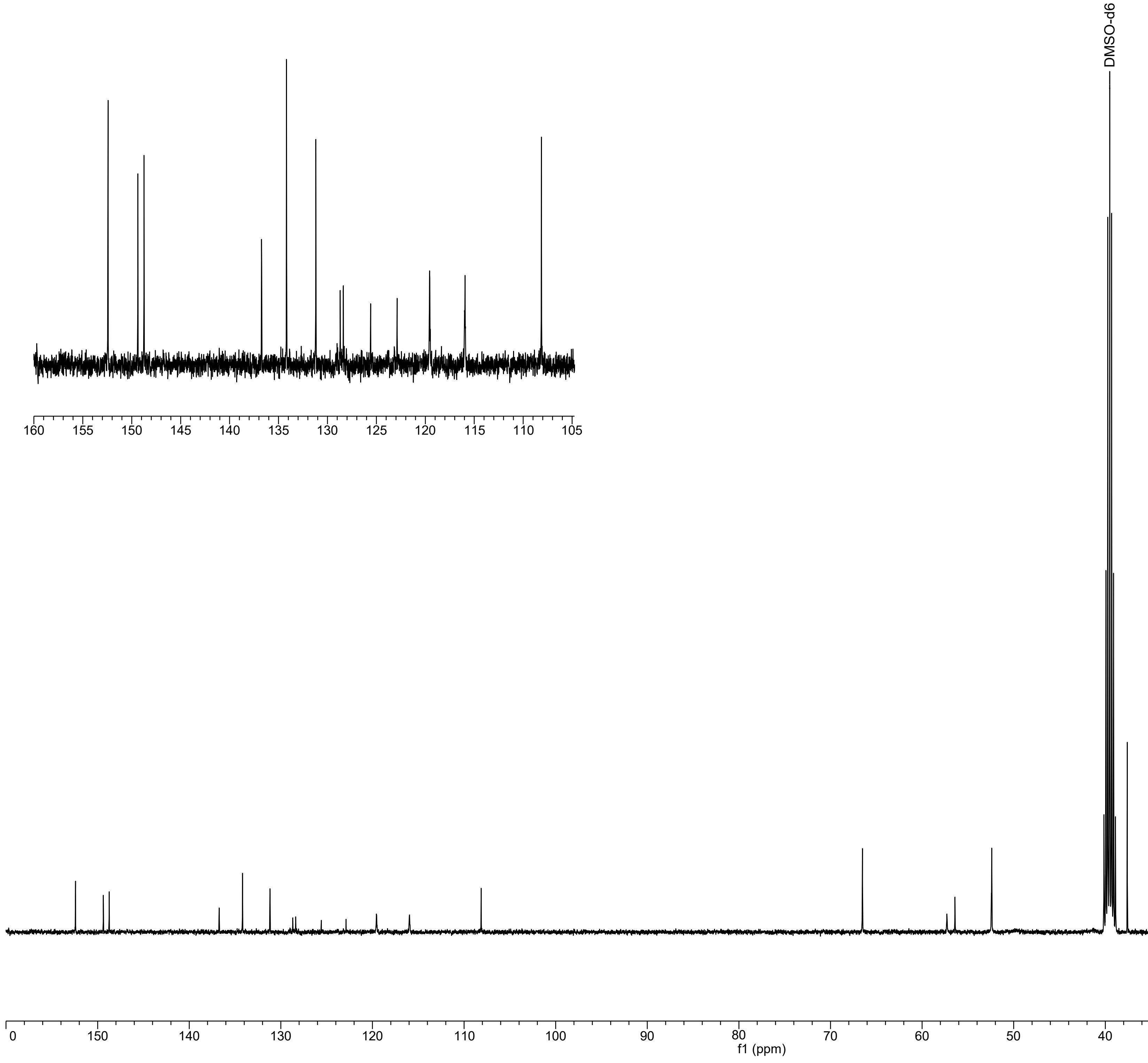

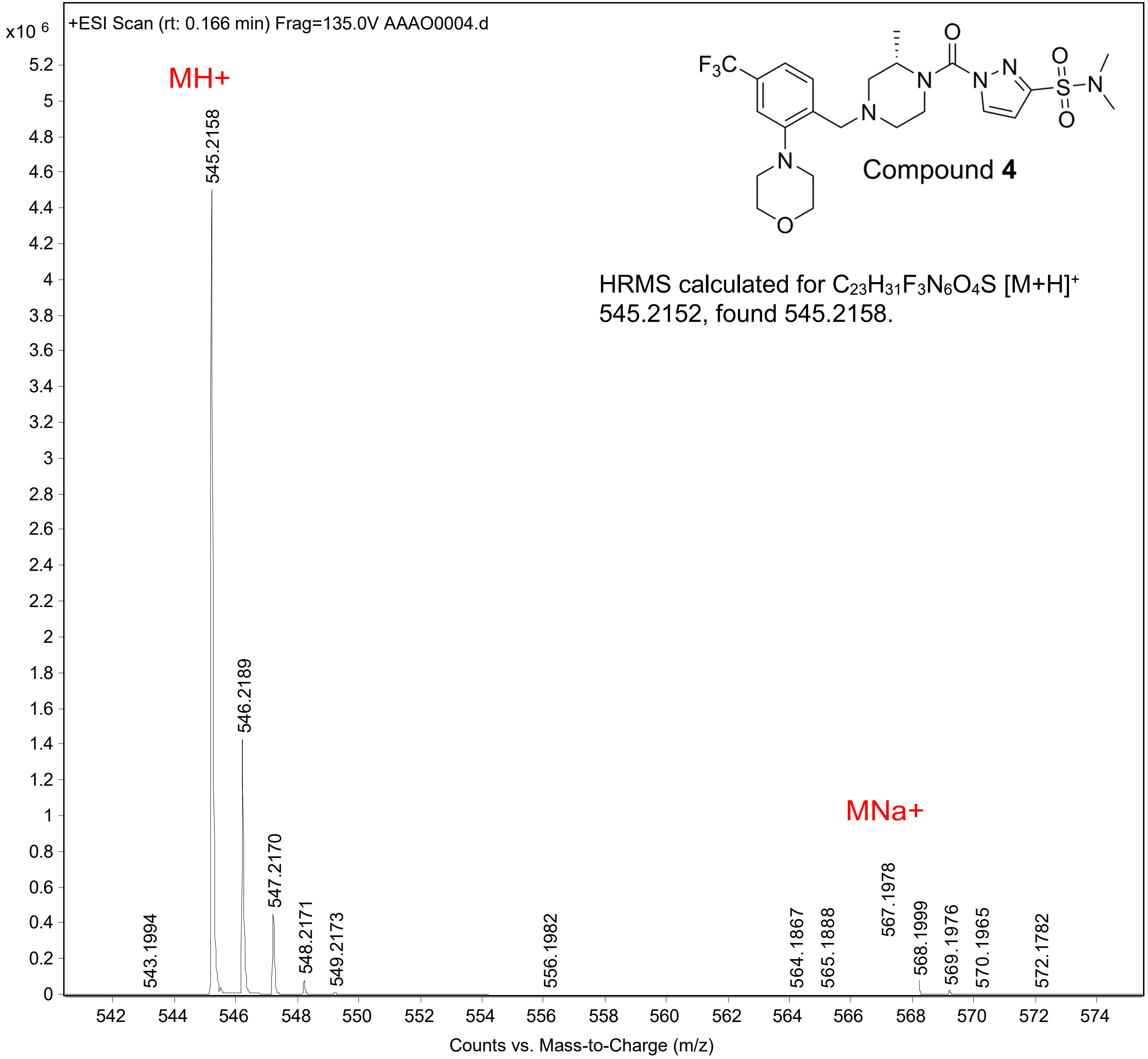
+ESI Scan (rt: 0.166 min) Frag=135.0V AAAO0004.d
x10 6
MH+
545.2158
546.2189
547.2170
548.2171
549.2173
543.1994
556.1982
564.1867
565.1888
567.1978
568.1999
569.1976
570.1965
572.1782
MNa+
$F_3C$
Compound **4**
HRMS calculated for $C_{23}H_{31}F_3N_6O_4S$ $[M+H]^+$ 545.2152, found 545.2158.
Counts vs. Mass-to-Charge (m/z)

Compound **5** (**ABD957**)

1H NMR (300 MHz, MeOD) δ 8.29 (d, J = 2.7 Hz, 1H), 7.66 (d, J = 8.0 Hz, 1H), 7.31 (d, J = 7.9 Hz, 1H), 7.18 (s, 1H), 6.82 (d, J = 2.7 Hz, 1H), 4.56 (br, 1H), 4.30 (d, J = 17.2 Hz, 2H), 4.20 (d, J = 13.8 Hz, 1H), 3.75 – 3.23 (m, 9H), 2.96 (d, J = 10.4 Hz, 1H), 2.83 (s, 6H), 2.79 (s, 1H), 2.43 (dd, J = 11.6, 3.7 Hz, 1H), 2.38 – 2.26 (m, 1H), 2.15 (s, 4H), 1.45 (d, J = 6.8 Hz, 3H).

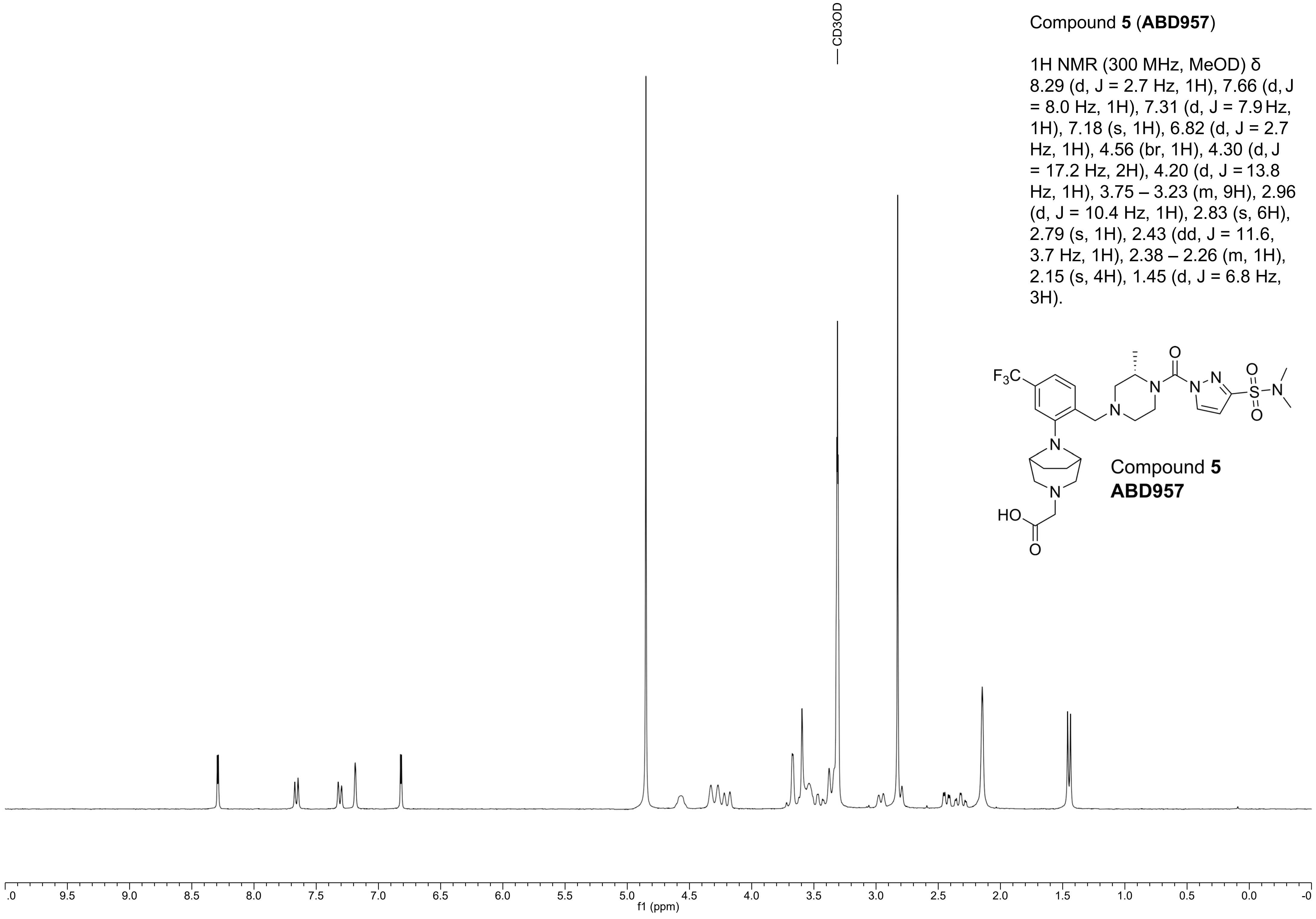


Compound **5**
**ABD957**

Compound **5** (**ABD957**)

13C NMR (101 MHz, DMSO-d6) δ 171.56, 150.33, 149.38, 148.73, 134.18, 133.86, 131.64, 128.23 (q, J = 30.7 Hz), 124.32 (q, J = 272.1 Hz), 117.11 (q, J = 3.4 Hz), 113.13 (q, J = 3.3 Hz), 108.14, 59.53, 59.34, 58.32, 58.26, 57.75, 57.61, 57.33, 52.55, 37.62, 26.93, 16.09.

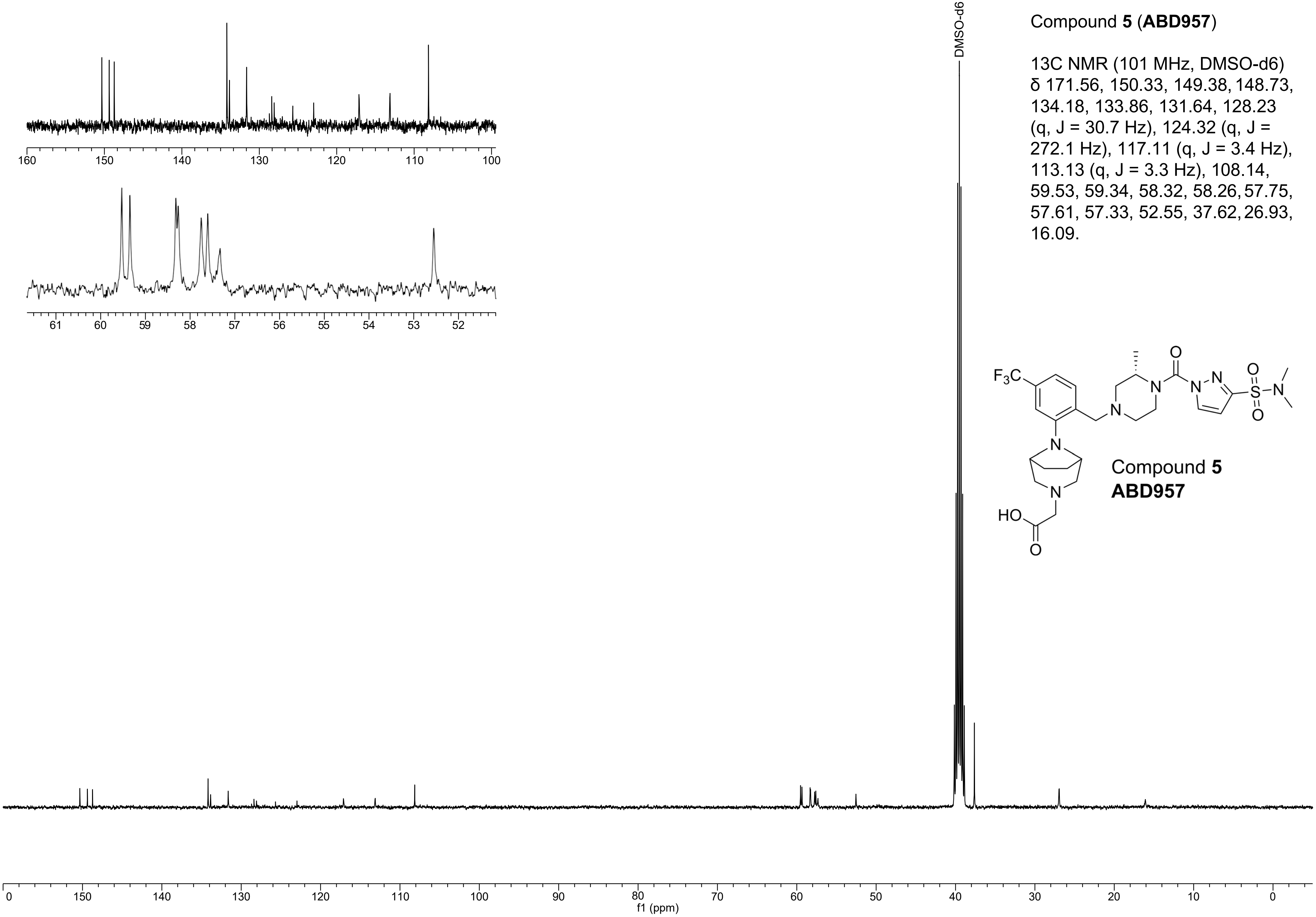


Compound **5**
**ABD957**

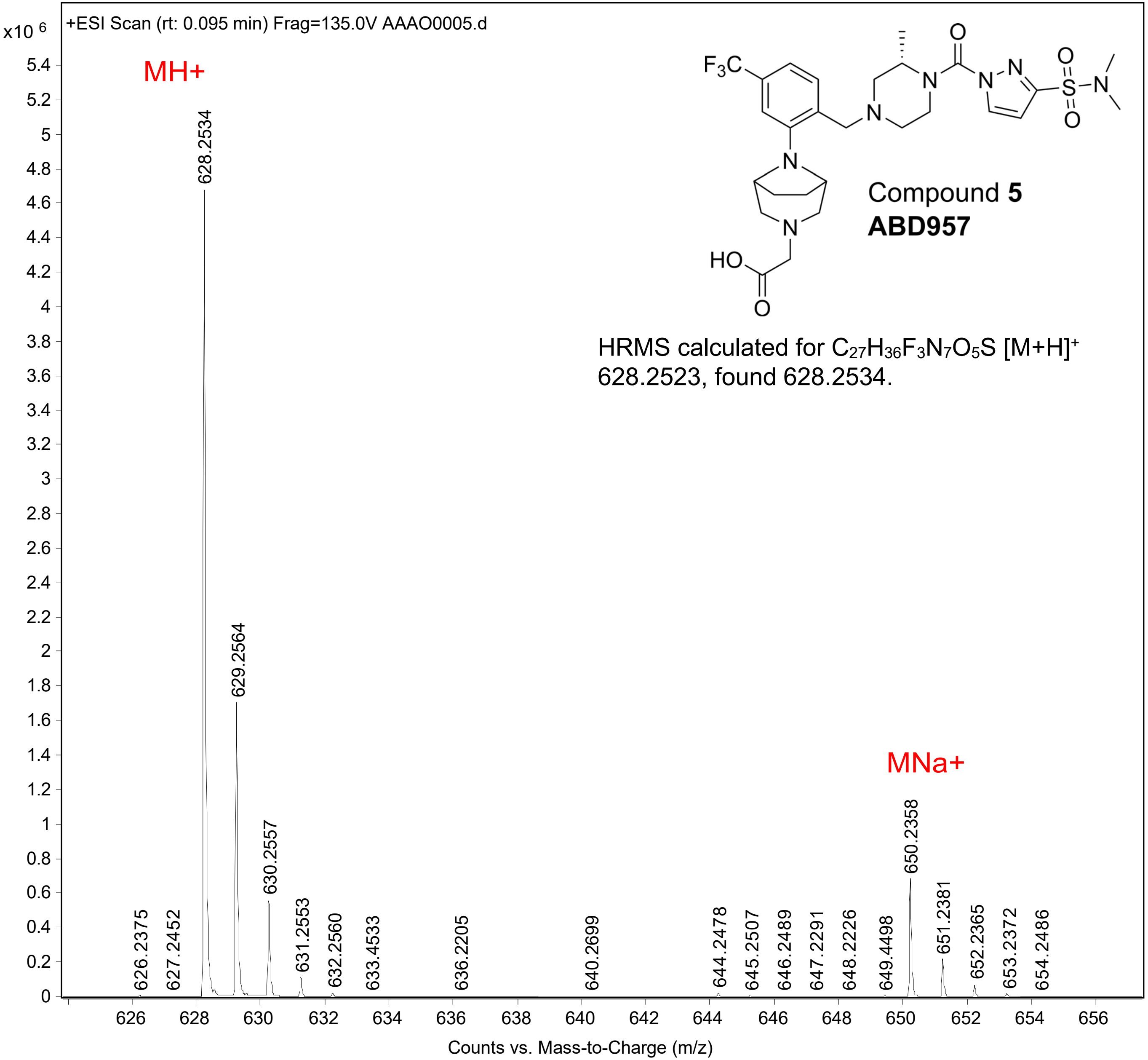

+ESI Scan (rt: 0.095 min) Frag=135.0V AAAO0005.d
x10 6
MH+
MNa+
628.2534
629.2564
630.2557
631.2553
632.2560
633.4533
636.2205
640.2699
644.2478
645.2507
646.2489
647.2291
648.2226
649.4498
650.2358
651.2381
652.2365
653.2372
654.2486
626.2375
627.2452
Compound 5
ABD957
HRMS calculated for $C_{27}H_{36}F_3N_7O_5S$ $[M+H]^+$ 628.2523, found 628.2534.
Counts vs. Mass-to-Charge (m/z)
